# Screening macrocyclic peptide libraries by yeast display allows control of selection process and affinity ranking

**DOI:** 10.1101/2024.08.24.609237

**Authors:** Sara Linciano, Ylenia Mazzocato, Zhanna Romanyuk, Filippo Vascon, Lluc Farrera Soler, Edward Will, Yuyu Xing, Shiyu Chen, Yoichi Kumada, Marta Simeoni, Alessandro Scarso, Laura Cendron, Christian Heinis, Alessandro Angelini

## Abstract

Macrocyclic peptides provide an attractive modality for drug development due to their ability to bind challenging targes, their small size, and amenability to powerful *in vitro* evolution techniques such as phage or mRNA display. While these technologies proved capable of generating and screening extremely large libraries and yielded ligands to already many targets, they often do not identify the best binders within a library due to the difficulty of monitoring performance and controlling selection pressure. Furthermore, only a small number of enriched ligands can typically be characterised due to the need of chemical peptide synthesis and purification prior to characterisation. In this work, we address these limitations by developing a yeast display-based strategy for the generation, screening and characterisation of structurally highly diverse disulfide-cyclised peptides. Analysis and sorting by quantitative flow cytometry enabled monitoring the performance of millions of individual macrocyclic peptides during the screening process and allowed us identifying macrocyclic peptide ligands with affinities in the low micromolar to high picomolar range against five highly diverse protein targets. X-ray analysis of a selected ligand in complex with its target revealed optimal shape complementarity, large interaction surface, constrained peptide backbones and multiple inter- and intra-molecular interactions, rationalising the high affinity and exquisite selectivity. The novel technology described here offers a facile, quantitative and cost-effective alternative to rapidly and efficiently generate and characterise fully genetically encoded macrocycle peptide ligands with sufficiently good binding properties to even therapeutically relevant targets.

## Introduction

Macrocyclic peptides are increasingly proving to be a valuable molecular format for drug development.^1^ At present, there are around eighty peptide therapeutics on the global market, of which more than forty are macrocyclic peptides, with a growing number of macrocyclic peptide drugs approved per year.^2,3^ Macrocyclic peptides have a number of favourable properties that make them an attractive modality for the development of therapeutic agents.^4^ They can bind to macromolecular targets with high affinity and selectivity. Moreover, they often display good proteolytic stability, and in some cases even display membrane permeability. Furthermore, they usually show a low inherent toxicity or antigenicity. Additionally, macrocyclic peptides can be efficiently produced by chemical synthesis and possess ease of modification. Besides, their modular structure and facile access to hundreds of different commercially available amino acid building blocks simplify the development of macrocyclic peptide variants with tailored properties. All these qualities well position macrocyclic peptides to bridge the gap between small molecule drugs and larger biologics such as antibodies.^4^

While numerous macrocyclic peptides continue to originate from the investigation and exploitation of naturally occurring peptides, recent technological advances and major breakthroughs in molecular biology paved the way for the development of *de novo* generated macrocyclic peptide ligands with desired qualities.^5^ Indeed, the advent of powerful combinatorial technologies such as phage display,^6,7^ mRNA display,^8,9^ bacteria display,^10,11^ and the split-intein based approach SICLOPPS ^12^ have exponentially accelerated the generation of macrocyclic peptide ligands to diverse protein targets for which no natural peptide ligands have been discovered. All these *in vitro* directed evolution approaches rely on a physical linkage between the ‘phenotype’ (expressed cyclic peptide) and ‘genotype’ (encoding DNA or RNA sequence). By applying such technologies, macrocyclic peptide ligands with desired properties are usually evolved following a similar scheme, comprising the generation of large combinatorial libraries of random genetically encoded macrocyclic peptides, multiple iterative cycles of selection, amplification and diversification. Recently, the development of innovative strategies for post-translational chemical and enzymatic modifications have greatly expanded the number of macrocyclic peptide formats that, in addition to the canonical disulfide-tethered peptides, can be examined by the aforementioned combinatorial technologies.^13–16^ These, coupled with the latest technological advances in screening procedures and automatisation, the emergence of new reagents and tools, and the affirmation of next-generation sequencing techniques, have enabled the construction of larger libraries and the isolation of macrocyclic peptide ligands with exquisite binding properties towards increasingly challenging targets.^17–19^

Though these technologies have been proven capable of generating and screening extremely large sizes of sequences and structurally diverse repertoires, thus enabling the rapid identification of macrocyclic peptide ligands against virtually any target, they still rely on rather difficult to control procedures as the performance of individual selected clones or populations cannot be easily monitored during the high-throughput screening. The success of a selection campaign is typically only seen after several weeks of work, once the isolated macrocyclic peptide molecules are identified. Moreover, the biophysical characterization of selected ligands often requires chemical synthesis or sub-cloning and recombinant production followed by purification, all additional phases which ultimately slow down and make the downstream process more complex and often costly.^15,20,21^ Finally, the number of selected macrocyclic peptide ligands that can be synthesised, purified and characterised is typically limited to 10 – 100 molecules.

Herein, we present a novel approach based on yeast surface display technology that addresses the abovementioned concerns. Since its invention, yeast surface display has proven to be a transformative tool for the direct evolution of multiple immunoglobulin- and non-immunoglobulin-based proteins.^22,23^ This technique was initially validated to enhance the binding affinity of an existing antibody fragment, but later proved highly effective also for isolating *de novo* proteins with fine-tuned binding affinities and specificities toward a wide range of targets from naïve combinatorial libraries.^24,25^ An advantage of yeast surface display technology coupled to fluorescence-activated cell sorting (FACS) is that it offers real-time monitoring of the iterative screening of large combinatorial libraries of diverse macrocyclic peptide ligands. By applying flow cytometry, information on the affinity, specificity, stability, and enrichment of individual yeast clones encoding diverse macrocyclic peptide ligands can be monitored in a continuous and quantitative manner through successive rounds of sorting.^22,26^ Moreover, by applying different labelling approaches, including equilibrium, kinetic and competition binding approaches, yeast display combined with flow cytometry can allow accurate adjustment of the selection stringency, enables rapid and fine epitope mapping, and favours quantitative discrimination between single macrocyclic peptide variants directly as cell surface fusions without the need for their chemical synthesis or recombinant expression and purification.^24,25,27^

Yeast display was applied before for the screening of peptide libraires but mostly for the engineering of naturally occurring linear and cystine knot peptides ^28^ or the affinity maturation or binding characterisation of existing molecules.^29,30^ Recently, first studies reported the exploitation of yeast display for the *de novo* generation of cyclic peptide ligands, such as the generation of binders to lysozyme and human interleukin-17 with dissociation constant (*K*_D_) values ranging from 300 nM to 3 μM.^31^ Similarly, screening of library of post-translationally enzymatically modified octapeptides yielded cyclic peptide ligands capable of binding two distinct domains of the yes-associated protein (YAP) with apparent *K*_D_ values ranging from 700 nM to 1.5 μM.^32^ However, the sizes of the libraries were still limited as well as the structural diversity of the peptides that were restricted to monocyclic peptides of 7 and 8 amino acids.

Hence, in this work we attempted to thoroughly explore the full potential of yeast surface display for the isolation and characterisation of macrocyclic peptide ligands with fine binding properties. Toward this goal we generated large and structurally diverse naïve combinatorial libraries encoding macrocycle peptide ligands with different topologies. To provide a more robust procedure we turned to disulfide-cyclised peptide ligands. The spontaneous intra-molecular oxidation of cysteine residues, present in the genetically encoded peptide sequences, is significantly easier and more reliable than post-translational cyclisation. In this regard, we foresee the eukaryotic expression machinery of yeast cells to support precise disulfide isomerization of cysteine-rich sequences ultimately enabling the generation of unique disulfide-tethered patterns that may be otherwise refractory to other systems.^22,26,28^ A key step in our work was to use a different yeast surface protein than the commonly applied α-agglutinin Aga1 and Aga2 proteins, where the ligand of interest is usually expressed as a fusion to Aga2 that is covalently linked to the membrane-anchored Aga1 via two inter-molecular disulfide bonds. In our work, we turned to a cysteine-free glycosylphosphatidylinositol (GPI) cell-surface anchor protein and expressed the disulfide-linked macrocyclic peptide ligands on the surface of yeast cells as aminoterminal fusion to the amino-terminal of the GPI anchor protein.^33^ We showed that by applying quantitative flow cytometry-based selections yeast displayed macrocyclic peptides with high structural diversity and fine binding properties against a panel of distinct protein targets can be rapidly identified. The technology appears capable to effectively pick and enrich rare macrocyclic peptides with different motifs and topologies for each target tested, even though from combinatorial libraries 100 times smaller than those attainable using other *in vitro* evolution tools. Further X-ray analysis of a selected ligand in complex with its target revealed optimal shape complementarity, large interaction surface, constrained peptide backbones and multiple inter- and intra-molecular interactions, explaining its high affinity and exquisite selectivity.

## Results and discussion

### Generation of yeast-encoded macrocyclic peptide libraries with large backbone diversity

We generated large and structurally diverse macrocyclic peptide libraries displayed on the surface of *Saccharomyces cerevisiae* cells (**Figure 1a**). Yeast displayed macrocyclic peptides are obtained by forming intra-molecular disulfide bridges between cysteine residues present in the genetically encoded peptide sequences. To avoid the formation of undesired inter-molecular disulfide bonds we appended the cysteine-rich peptide sequences at the N-terminus of a cysteine-free GPI cell-surface anchor that has been previously developed to tethered small proteins on the surface of yeast cells.^33^

**Figure 1.**
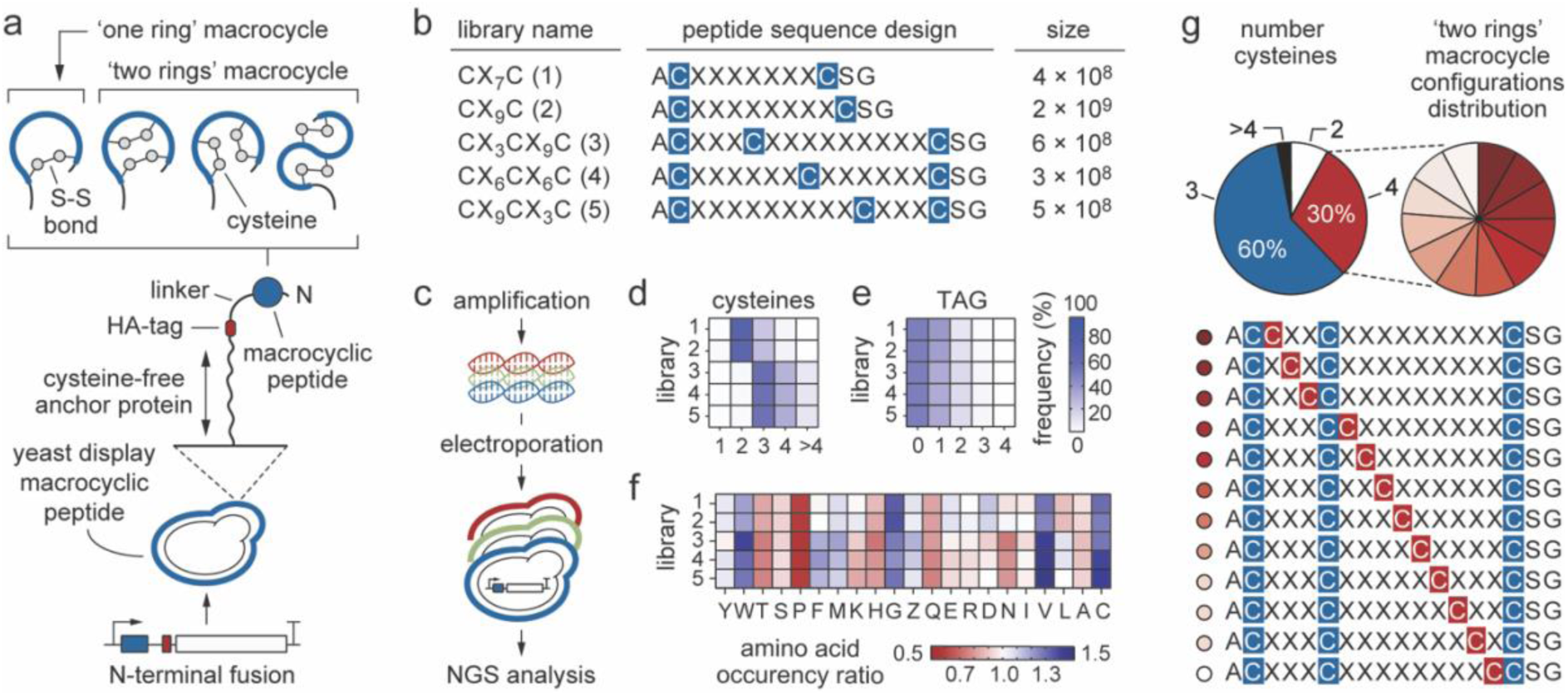
Yeast-encoded macrocyclic peptide libraries. **a**) Schematic representation of the yeast surface display macrocyclic peptide system developsed in this work. The cysteine-rich peptide sequence of interest (blue) is expressed as N-terminal fusion of the cysteine-free glycosylphosphatidylinositol (GPI) cell-surface anchor protein (black line). A long and flexible (G_4_S)_3_ linker is placed between the macrocyclic peptide and the immunofluorescence hemagglutinin (HA) tag (^N^YPYDVPDYA^C^, red). Bridging of one pair of cysteines by disulfide bridge yields macrocycle peptides with one ring and a unique topology while bridging two pairs of cysteines by disulfide bridges yields macrocycle peptides with two rings and three different topologies that can be sampled for binding to protein targets; **b**) Macrocyclic peptide yeast display library design and size comparison. Five naïve peptide libraries were generated to include either two or three cysteines (C, blue) at fixed positions and between 7 to 12 random amino acids (X). The naïve peptide libraries were numbered as follows: library 1 = CX_7_C, library 2 = CX_9_C, library 3 = CX_3_CX_9_C, library 4 = CX_6_CX_6_C and library 5 = CX_9_CX_3_C. The size of each library (number of unique peptide sequences) is reported and was determined as previously described;^22,63^ **c**) The five yeast-displayed macrocyclic peptide naïve libraries were constructed using degenerated primers that allow all 20 amino acids in the randomized positions (X = NNK) and homologous recombination-based methods.^22,63^ Heat map indicating the frequency of the number of cysteines (**d**) and the unique ‘TAG’ amber stop codon (**e**) in each naïve library. Frequencies were determined by next generation sequencing (NGS) of single naïve library. The intensity of the colour correlates with the frequency of a given cysteine residue or TAG stop codon in the peptide sequence. High and low frequency values are shown in dark and light colours, respectively; **f**) Heat map representing the experimental frequency of each individual amino acid determined by NGS analysis compared to the theoretical expected values for a NNK codon. The individual amino acids (20) are indicated by a one-letter code. The amber stop codon is indicated by the letter Z. The intensity of the colour correlates with the frequency ratio. Enriched amino acids in naïve libraries are shown in dark blue, whereas depleted amino acids are shown in dark red. Amino acids with an experimental/theoretical ratio of one are shown in white; **g**) Frequency of the occurrence and distribution of additional cysteine in library 3. The three constant cysteine (C) residues are shown in grey, while the fourth, which appears randomly, is highlighted in red.

The use of this cysteine-free GPI anchor protein to display disulfide-tethered macrocyclic peptide ligands on the surface of yeast cells should not only minimise the risk of undesired intermolecular disulphide bridges formation between cysteine residues present within the peptide ligands and those existing in both Aga1 and Aga2 proteins but could also facilitate the use of the system in different yeast strains, ultimately enhancing the flexibility of the system. Indeed, while the commonly applied galactose-inducible heterodimeric Aga1-Aga2 yeast display system requires the use of engineered yeast strains bearing the Aga1 gene stably integrated into the yeast chromosome, the gene encoding the disulfide-tethered macrocyclic peptide ligands fused to the monomeric cysteine-free GPI anchor protein cloned into an episomal plasmid can be directly covalently connected to the cell wall of any yeast strain. To secure distance and minimise potential steric hindrance with detection reagents, we placed a flexible glycine-serine (G_4_S)_3_ linker between the peptide sequence and the immunofluorescence hemagglutinin (HA) tag (**Supplementary figure 1**). Detection of the HA tag using fluorescently labelled antibody combined with flow cytometry allows not only rapid quantification of the expression level of macrocyclic peptide ligands on yeast cells, but also normalization of the binding signal to surface expression.

We generated peptide libraries preferentially encoding either ‘one ring’ or ‘two rings’ macrocycle topologies with sequence diversity comprised between 3 × 10^8^ and 2 × 10^9^ (**Figure 1b** and **Supplementary table 1**). We designed yeast-encoded ‘one ring’ macrocyclic peptide libraries of the format C(X)*_n_*C (X = any amino acid, *n* = 7 or 9), in which all sequences contain predominantly two fixed cysteines. Similarly, we created yeast-encoded ‘two rings’ macrocyclic peptide libraries of the format C(X)*_n_*C(X)*_m_*C in which all peptides contain predominantly three constant cysteines spaced by different numbers of random amino acids (X) wherein *n* = 3, 6 or 9 and *m* = 9, 6 or 3 (**Figure 1b**). The random amino acid positions in all libraries were encoded by ‘NNK’ codons resulting in a ∼3% possibility of finding a cysteine residue in each of the randomised position (**Supplementary table 2**). The theoretical probability of encountering an additional cysteine residue in the ‘two rings’ C(X)*_n_*C(X)*_m_*C macrocyclic peptide libraries was estimated to be 37% (**Supplementary table 3**). We reasoned that a first disulfide bond can be formed between any of the two of the three fixed cysteines and a second disulfide bridge could be formed between the third and the fourth cysteine appearing in any of the randomised positions. In total, as many as 3(*l* – 4)!/2(*l* – 6)! different macrocyclic peptide topologies with a length of *l* amino acids could be formed (**Supplementary figure 2**).^34^ Such a large structural diversity should increase the chance of finding ‘two rings’ macrocycle peptides that have a shape complementary to the surface of protein targets. Next generation sequencing (NGS) analysis of the naïve libraries confirmed that the macrocyclic peptides present the expected number of cysteines, amino acid frequency and topological diversity (**Figure 1c-g**, **Supplementary figure 3** – **5** and **Supplementary table 4** – **8)**.

### Screening of yeast-encoded macrocyclic peptides toward a wide range of protein targets

To validate our technology, we screened yeast-encoded macrocyclic peptide libraries towards five highly diverse protein targets (PT) namely aldolase (PT1), streptavidin (PT2), carbapenemase GES-5 (PT3), carbonic anhydrase (PT4) and α-chymotrypsin (PT5; **Figure 2a-b** and **Supplementary table 9**).

**Figure 2.**
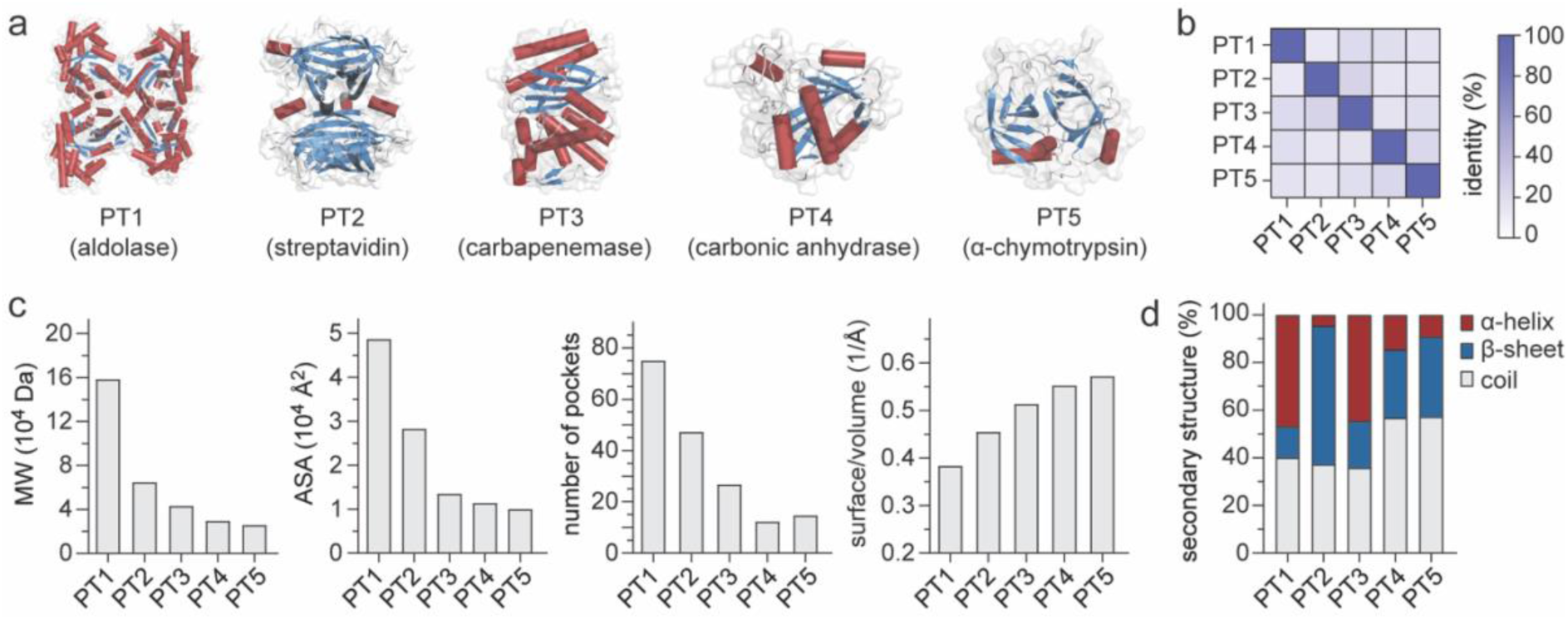
Biochemical properties of the chosen protein targets. **a**) Three-dimensional structure of five different protein targets (PT) used during selection screening: aldolase (PT1, PDB identification code 1QO5), streptavidin (PT2, PDB identification code 7EK8), carbapenemase GES-5 (PT3, PDB identification code 6TS9), carbonic anhydrase (PT4, PDB identification code 1VE9) and α-chymotrypsin (PT5, PDB identification code 6DI9). The overall target secondary structure is shown as cartoon and coloured by secondary structure (α-helices = firebrick, β-sheet = sky-blue, random coil = white). Proteins are ordered according to their molecular weight (MW), from largest (left) to smallest (right); **b**) Heat map showing the amino acid identity among different PTs. The intensity of the colour correlates with the identity percentage. High and low identities are shown in dark blue and white, respectively; **c**) From left to right, bar chart showing the distribution of the MW, solvent accessible surface area (A.S.A.), number of pockets ^60^ and surface/volume ratio of each PT; **d**) Columns graph reporting the percentage of secondary structure content (α-helices = firebrick, β-sheet = sky blue, random coil = light grey) for each PT tested.

These proteins were chosen based on their low-sequence identities, and different structural and biochemical properties (**Figure 2c-d** and **Supplementary table 10**). The use of such distinct proteins is key because it allows us to evaluate whether the technology can be widely applicable or if it is instead biased for certain types of protein targets. For instance, the presence of multiple copies (10^4^ – 10^5^) of a macrocycle peptide ligand displayed on the surface of a yeast cell could lead to undesired polyvalent interactions that might synergize to enhance the apparent binding affinity. This effect is commonly referred to as ‘avidity’ and can occur in the presence of a multivalent soluble target that can be simultaneously recognized by multiple copies of macrocyclic peptide ligands present on the surface of yeast.^25,27,35^ To better comprehend this potential issue, we included in our screening campaign two tetrameric proteins (PT1 and PT2) in addition to the monomeric ones (PT3 – PT5; **Supplementary table 10**). Furthermore, to better appreciate the potentiality of our technology, we picked two protein targets (PT2 and PT5) toward which macrocyclic peptide ligands have previously been isolated using well established *in vitro* directed evolution tools such as phage and mRNA display (**Supplementary figure 6)**. The five selected PTs were chemically biotinylated and their purity, degree of monodispersity and multimeric state confirmed using size-exclusion chromatography (**Supplementary figure 7**).

To increase the likelihoods of isolating macrocyclic peptide ligands with desired binding properties against each PT, we applied highly avid magnetic bead (MB) separations followed by multiple rounds of fluorescence-activated cell sorting (FACS; **Figure 3a-b**).

**Figure 3.**
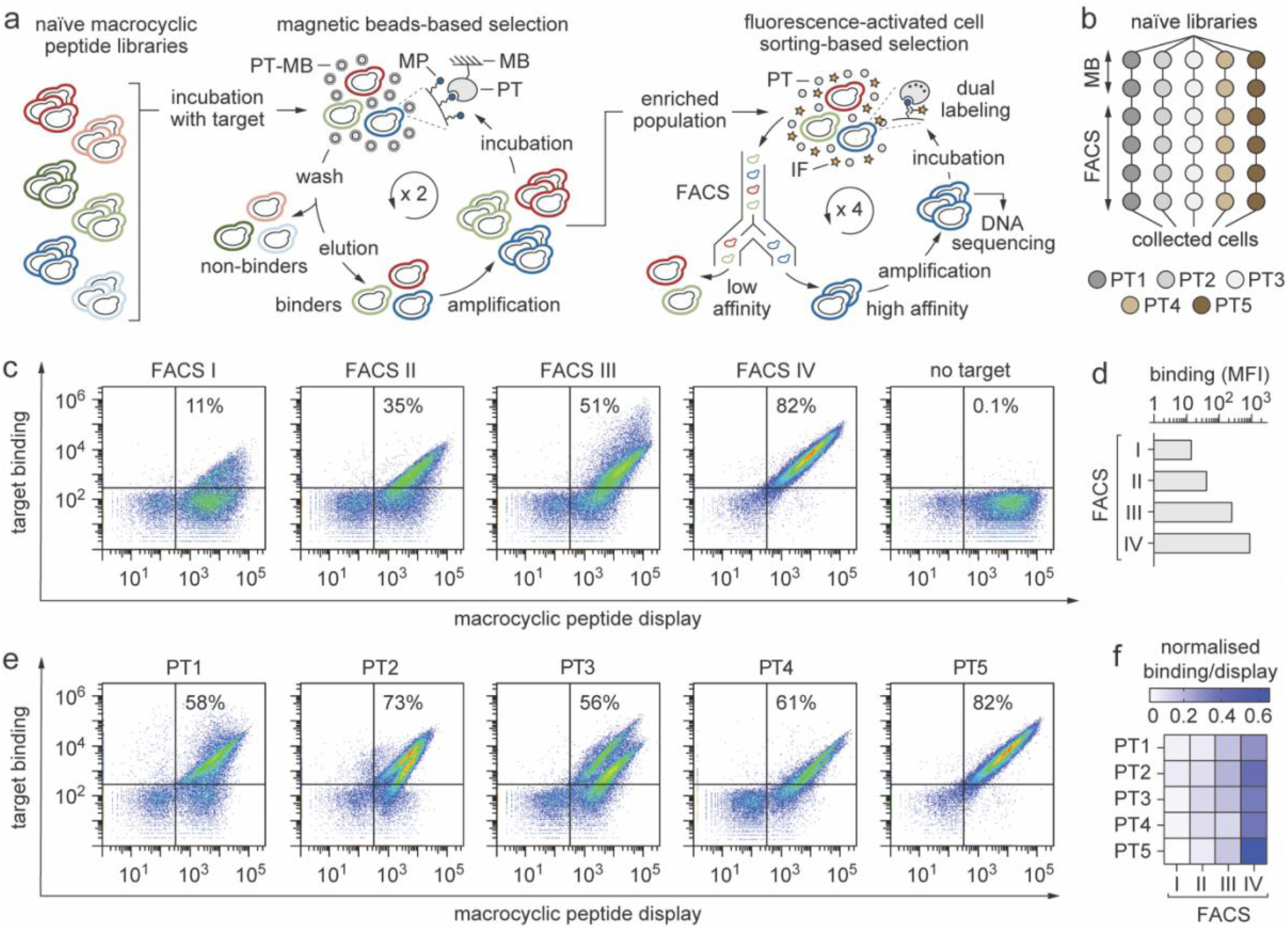
Selection of yeast-encoded macrocyclic peptides towards multiple protein targets. **a**) General flowchart applied to identify yeast-encoded macrocyclic peptide binders towards five highly diverse PTs. The five yeast-encoded macrocyclic peptide libraries (CX_7_C, CX_9_C, CX_3_CX_9_C, CX_6_CX_6_C and CX_9_CX_3_C) were pooled together and then incubated separately with each PT through two rounds of magnetic bead (MB) separation followed by four rounds of flow cytometry (FACS) based selection. In the flow chart of MB-based selection, the biotinylated PT immobilised on MB (PT-MB) is represented as a grey circle (PT) surrounded by a dotted ring (MB). In FACS-based selection, the soluble biotinylated PT is represented as a grey circle (PT) while the fluorescently labelled anti-HA antibody (IF) is depicted as an orange star; **b**) Schematic representation of the iterative selection pathways applied to isolate yeast-encoded macrocyclic peptide binders against five different PTs. Two cycles of MB-based screening followed by four cycles of FACS sorts were applied; **c**) Density plots of a representative polyclonal population of yeast cells encoding different macrocyclic peptides against PT5 that has been enriched from 11% to 82% through four cycles (I, II, III and IV) of FACS. Each dot represents two fluorescent signals of a single yeast cell. The fluorescence intensity on the *y*-axis is a measure of the amount of biotinylated PT bound to the surface of a yeast cell (DyLight 650; ‘target binding’) whereas the fluorescence intensity on the *x*-axis is a measure of the number of macrocyclic peptide molecules expressed on the surface of a yeast cell (DyLight 488; ‘macrocyclic peptide display’); **d**) Columns graph reporting the geometric mean fluorescence (MFI) measured for the polyclonal population of yeast cells encoding different macrocyclic peptides against PT2 through four cycles of FACS; **e**) Density plots of the enriched polyclonal population of yeast cells encoding different macrocyclic peptides against the respective PT after the fourth and last FACS cycle; **f**) Heat map displaying the ratio between the binding and display MFI of the polyclonal population after each round of FACS. The intensity of the colour correlates with MFI value. High and low intensity are shown in dark blue and white, respectively.

The initial use of magnetic bead-based screening allows the isolation of macrocyclic peptides within a relatively short period of time and in a high-throughput combinatorial manner.^22^ Moreover, we assumed the multivalency of the yeast display system combined to the multiple copies of the biotinylated PT immobilized on streptavidin-coated magnetic beads would facilitate highly avid interactions, thus promoting the isolation of weak macrocyclic peptide binders otherwise difficult to identify by flow cytometry.^26,35^ In this study we applied two iterative cycles of MB-based selections for each PT (**Figure 3a-b**). Each MB-based screening comprised growth of yeast cells, expression of the macrocyclic peptide on the yeast surface, binding to the immobilized biotinylated PT, washing, and expansion of the isolated bound yeast cells. Prior to ‘positive selection’ against each single PT, we also performed a ‘negative selection’ process to deplete macrocyclic peptide-binding nude streptavidin-coated MB. Notably, rather than exposing each naïve library separately with each different PT, we decided to mix the five libraries all together and let the individual PT pick the most suitable macrocyclic peptide topology.

We then promoted the enrichment of yeast cells isolated after the MB separations by allowing them to evolve through sequential cycles of equilibrium based FACS selections (**Figure 3a-b**).^22^ To avoid enrichment of potential macrocyclic peptide binders against labelling reagents we alternated the use of neutravidin and streptavidin during FACS sorts. By applying a two-colour labelling scheme, with one fluorescent probe to monitor macrocyclic peptide expression on the surface of yeast cells and another to monitor binding of macrocyclic peptide ligand to biotinylated PT, we fostered selection and enrichment of yeast-encoded macrocyclic peptides with desired affinity, specificity, and stability concomitantly (**Figure 3c-d**). Particularly, in this work we decided to maintain a constant and relatively high concentration (1 μM) of each PT during all rounds of FACS-based selection (**Supplementary figure 8**). This allowed us not only to identify macrocyclic peptides with a wider range of binding affinities, but also to obtain a more comprehensive view of the extent of amino acid sequence and topological diversity we could attain for each PT.

Overall, such evolutionary strategy enabled isolation and enrichment of yeast-encoded macrocyclic peptide ligands against all five distinct PTs employed in this work (**Figure 3e-f** and **Supplementary figure 8**). Though PTs containing large binding pockets, such as proteases, are apparently easier to target than proteins with featureless surfaces, here we have shown that the chances of identifying macrocyclic peptides against varying surface structures increases if screening is performed in an unbiased manner using large and structurally diverse combinatorial libraries.

### Selected yeast-encoded macrocyclic peptides revealed differences in amino acid sequences and topologies

To uncover the identity of the selected macrocyclic peptide ligands, we applied both Sanger and next generation sequencing (NGS). We started with Sanger sequencing because it allows us to prepare DNA samples from the same yeast cells that will be further used to characterise the macrocyclic peptide variant as a cell surface fusion, letting us to concomitantly obtain information on both the amino acid composition and binding properties of each ligand in just few days. To prepare DNA samples for Sanger sequencing, we plated a fraction of each flow cytometry sorted population on selective solid media, picked individual colonies, and further extracted and purified the DNA plasmids present within the yeast cells. Sequence analysis of approximately fifteen unique clones derived from each collected cell population revealed the presence of thirteen unique macrocyclic peptide ligands (MP) with different topologies and varying amino-acid compositions and ring sizes (**Figure 4a**). Notably, we did not detect cross-contamination among sequences derived from different populations, further proving the technology’s ability to effectively separate and enrich unique binders against different protein targets in parallel. Though rapid and cost-effective, Sanger sequencing allows analysis of a very limited portion of each flow cytometry sorted population (<0.0001% of the total collected cells).

**Figure 4.**
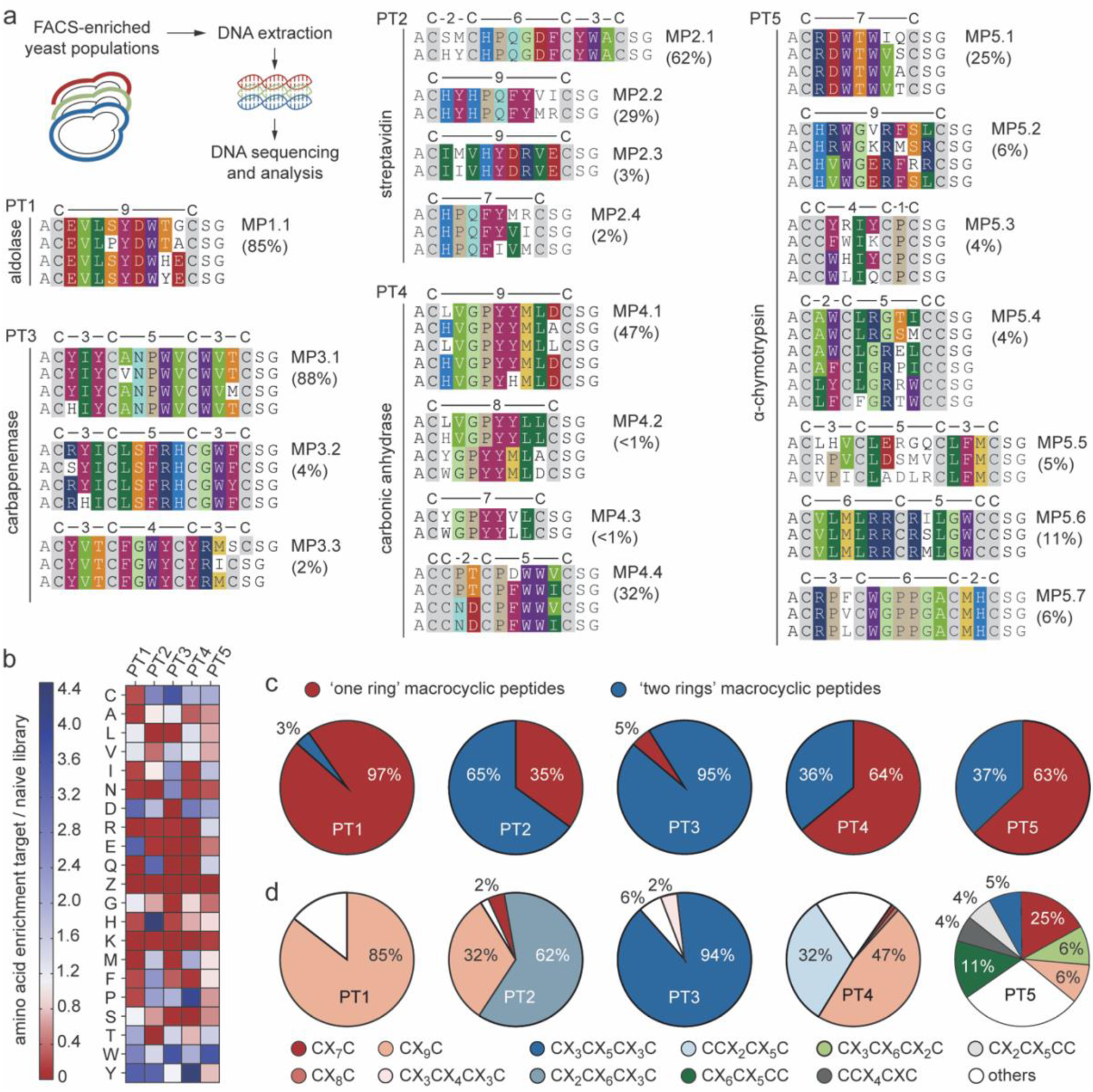
High-throughput sequencing analysis of selected target-specific macrocyclic peptides encoded by yeast. **a**) DNA extracted from FACS enriched yeast cell populations was sequenced using both Sanger and next generation sequencing (NGS) methodologies. DNA samples for high-throughput sequencing analaysis were amplified by PCR, indexed using target-specific barcodes, processed using NovaSeq Illumina NGS technology and the obtained FASTQ files analysed using MatLab scripts. The NGS analyses workflow of selected libraries consist of in *i*) subdivision based on barcode sequence and quality filter, *ii*) translation of peptide sequence, *iii*) depletion of unspecific motif and *iv*) clusterisation in families sharing a homology equal to or higher than 75%. Amino acid sequences of yeast-encoded macrocyclic peptides selected against each protein target (PT) are shown. The amino acid sequences are arranged in groups according to sequence similarities. The amino acids are indicated as one letter code. Identical or similar amino acids between different peptide sequences are highlighted in colours (C: grey; E and D: red; G: light green; V and A: intense light green; I and L: dark green; S and T: light orange; Y and F: purple; W: violet; H: indigo; P: light brown; Q and N: light blue; R and K: dark blue; M: yellow). Within a single macrocyclic peptide family, the amino acid sequences were listed starting from the clone with the highest abundance (top) to the one with the lower (bottom). Only sequences with a percentage of abundance >0.1% are reported; **b**) Heat map visualization of the type of amino acid residues that were enriched or depleted during selection when compared to naïve libraries. The amino acids are indicated as one letter code. The colour intensity correlates with the occurrence of each amino acid, with enriched or depleted residues shown as dark blue and dark red, respectively; **c**) Pie chart visualization of the different macrocyclic peptide topologies enriched during selection against the five different PTs. Macrocyclic peptides with ‘one’ and ‘two’ rings are shown in red and blue respectively; **d**) Pie chart visualization of the relative abundance of the most frequently selected macrocyclic peptide topologies across different PTs: CX_7_C (dark red), CX_8_C (dark salmon), CX_9_C (light salmon), CX_3_CX_5_CX_3_C (dark blue), CX_2_CX_6_CX_3_C (blue-grey), CCX_2_CX_5_C (light blue), CX_3_CX_4_CX_3_C (pink), CCX_4_CXC (dark grey), CX_6_CX_5_CC (light grey), CX_6_CX_5_CC (dark green), CX_2_CX_6_CX_2_C (light green) and others (white).

To gain a better insight into the genetic diversity and abundance of the target-specific macrocyclic peptide ligands present in each population of collected yeast cells, we applied NGS analysis on pool of DNA plasmids extracted directly from flow cytometry sorted polyclonal yeast cells (**Figure 4a** and **Supplementary figure 3**). We implemented sequence data filters to reduce biases originating from the sequencing method and applied sequence correction algorithms to prevent identification of false consensus motifs. Next, we compared the obtained peptide sequence datasets to each other in order to *i*) identify target specific peptide binding motifs, *ii*) uncover the topology and ring size distribution, *iii*) determine the frequency of each amino acid within each peptide loop and *iv*) cluster the most abundant peptides in consensus families (**Supplementary figure 3** and **Supplementary table 11** – **14**). NGS sequence analysis *i*) confirmed the presence of all thirteen out of nineteen (∼70%) unique and most abundant MPs previously identified by Sanger sequencing, *ii*) revealed the existence of at least three unique peptide families for four out of the five PTs tested, *iii*) uncovered six new macrocyclic peptide families (MP2.3, MP3.3, MP4.2, MP4.3, MP5.3 and MP5.5) and *iv*) provided a better view of the sequence diversity and abundance distribution among peptide sequences present within each macrocyclic peptide family (**Figure 4a** and **Supplementary table 15**). Each family contains several sequences that differ in one or more amino acids. The abundance of single macrocyclic peptide sequences present within each family varies. We cannot exclude that some low abundant sequences might be the result of artefacts occurring during NGS preparation and processing rather than peptide ligands encoded by the yeast. However, a dominant family with relative abundance greater than 45% was identified for each PT. Moreover, the amino acid sequences of the isolated macrocyclic peptides differ from target to target (**Figure 4a-b**). These differences appear to persist even at the level of macrocyclic peptides that bound the same PT, although some shared consensus motifs that have been identified for two targets (**Figure 4a**). For example, three of the four macrocyclic peptide families isolated against PT2 (streptavidin) contain the ‘HPQ’ motif, well-known as peptide binder of this target (**Supplementary figure 6)**, though localized within diverse ring sizes and topologies (‘one ring’ and ‘two rings’ macrocyclic peptides; **Supplementary figure 9**). Similarly, three out of the four peptide families selected against PT4 present a conserved ‘GPYY’ motif. Interestingly, this sequence motif has been found within ‘one ring’ macrocyclic peptides with different sizes, including an eight-amino acid loop that was not included in the initial design. Furthermore, sequence analysis allowed us to better appreciate what kind of topologies best suited each PT. While protein targets PT1 and PT4 appear to be favourably recognised by ‘one ring’ macrocyclic structures, ‘two rings’ topologies were preferentially enriched during selections against PT2, PT3 and PT5, although with a different relative abundance distribution (**Figure 4c**). In general, ‘one ring’ macrocyclic peptides with a larger ring (9-amino acid ring: CX_9_C) appear to be more frequent than those with a smaller ring (7-amino acid ring: CX_7_C). However, a family of ‘one ring’ macrocyclic peptides with a 7-amino acid ring that has a 5-fold higher abundance than those with 9 amino acids has been identified solely for PT5 (**Figure 4a**, **c** and **d**). Interestingly, the sequences of MP4.2, MP4.4, MP5.3, and MP5.4 peptide families were unexpected and are probably the result of either impurities in oligonucleotide synthesis or the introduction of two additional cysteines into the monocyclic peptides. Notably, among the seventy different ‘two rings’ macrocyclic peptide topologies available in the naïve libraries (**Supplementary figure 4** and **5**), only eight of them (CX_3_CX_5_CX_3_C, CX_2_CX_6_CX_3_C, CCX_2_CX_5_C, CCX_4_CXC, CX_2_CX_5_CC, CX_6_CX_5_CC and CX_3_CX_6_CX_2_C, CX_3_CX_4_CX_3_C) were effectively enriched during selection (**Figure 4d**). Except for the CX_3_CX_5_CX_3_C topology, which is present in a family of macrocyclic peptides selected against both PT3 and PT5, the other six ‘two rings’ topologies are exclusive to individual protein targets, with CX_2_CX_6_CX_3_C being identified only in a peptide family targeting PT2, CCX_2_CX_5_C in a peptide family binding PT4, while CCX_4_CXC, CX_2_CX_5_CC, CX_6_CX_5_CC and CX_3_CX_6_CX_2_C topologies have been detected only in peptide families enriched against PT5 (**Figure 4d**). Notably, no peptide sequences containing three cysteines were isolated even though they were highly represented in the naive libraries (∼60% total clones). This further demonstrates the capability of our tool to generate and enrich exclusively ‘one ring’ and ‘two rings’ macrocyclic peptides.

In summary, sequencing analysis proved the ability of our technology to effectively isolate, fine discriminate and amplify very rare yeast-encoding disulfide-tethered macrocyclic peptides with different motifs and topologies for each target tested, even from combinatorial libraries 100 times smaller than those attainable using other *in vitro* evolution tools.

### The majority of selected yeast-encoded macrocyclic peptides exhibit binding affinities in the nanomolar range

To determine the binding affinities of the isolated macrocyclic peptides, we titrated yeast-displayed macrocyclic peptides into solutions with varying concentrations of the corresponding biotinylated PT (**Figure 5a**). This technique allows selected macrocyclic peptide ligands to be rapidly characterized directly on the surface of yeast cells, eliminating the need for additional chemical synthesis and purification steps. By normalizing the mean fluorescence intensity from the binding signal to the mean fluorescence intensity from the display signal, as a function of PT concentration, we rapidly determined the apparent equilibrium dissociation constant (*K*_D_^app^) of each individual selected clone. We characterised solely the most abundant macrocyclic peptide of each consensus family (**Figure 5b-f** and **Supplementary table 16**). The determined *K*_D_^app^ values span 10000-fold, ranging from 0.2 nM to 1 µM. Particularly, for PT1, PT2 and PT5 (three out of the five protein targets tested), all identified macrocyclic peptides showed *K*_D_^app^ values below 10 nM (**Figure 5b**, **c**, **f** and **Supplementary table 16**). For the other two targets examined (PT3 and PT4), the *K*_D_^app^ values were on average higher (0.04 to 3 µM) with the only exception of PT4, for which a macrocyclic peptide (MP3.2) with *K*_D_^app^ value of 2.2 nM was also isolated (**Figure 5d**, **e** and **Supplementary table 16**). Notably, the *K*_D_^app^ values found for different macrocyclic peptides binding a specific PT are often very close to each other (**Figure 5g**). We assume that rather than the topology of the macrocyclic peptide, it is the propensity of the PT itself that determines the attainable binding affinity.

**Figure 5.**
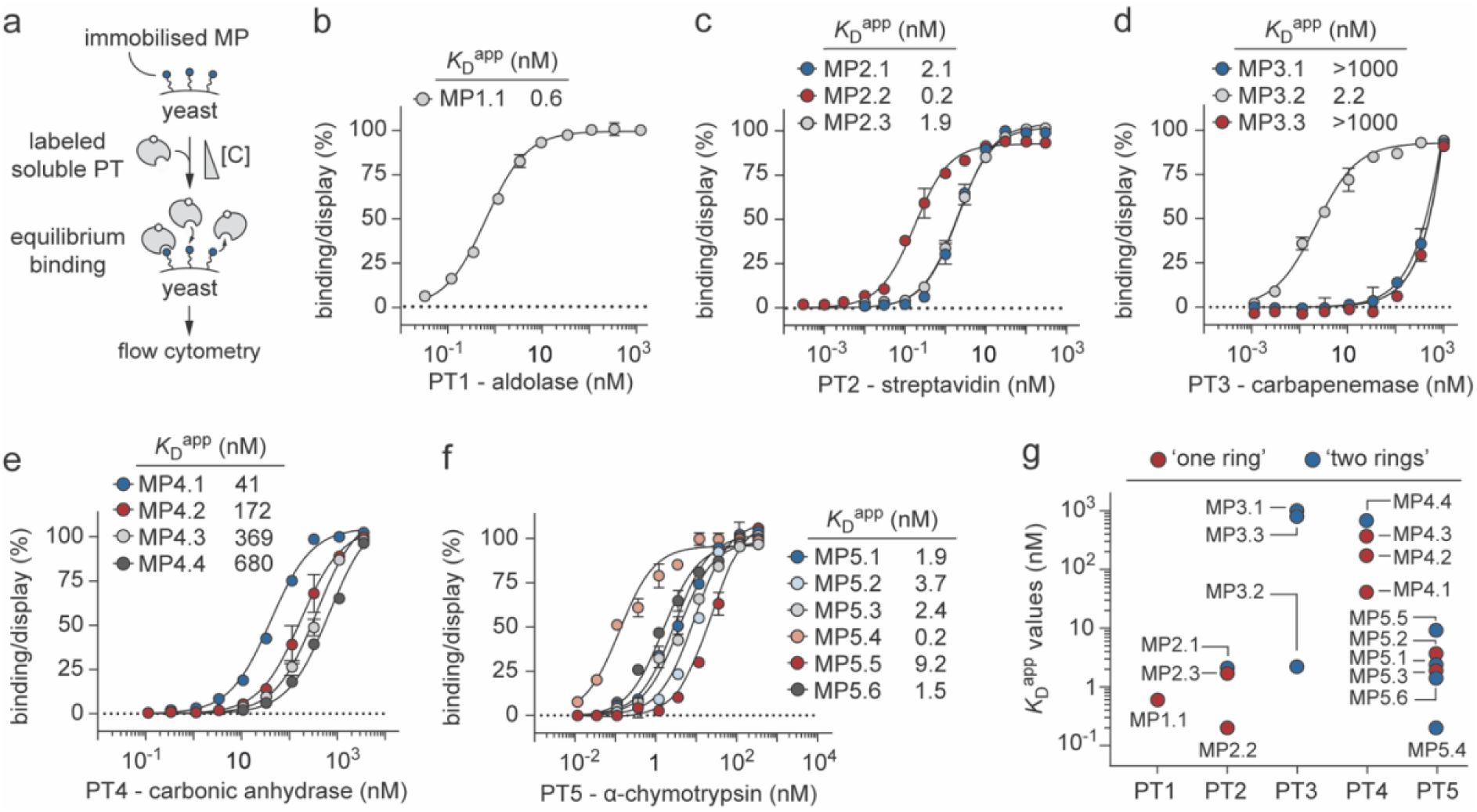
Determination of the binding affinities of cell-anchored macrocyclic peptides against soluble protein targets. **a**) Schematic representation of the apparent binding affinity determination using yeast surface titration. Yeast cells expressing the desired macrocyclic peptide (MP) on the cell surface are incubated with varying concentrations of soluble biotinylated PT. The binding is reported as median fluorescence intensity, and it is proportional to the amount of PT bound to the macrocyclic peptides expressed on the surface of the yeast cell. Binding isotherms of the most abundant yeast-displayed macrocyclic peptide clones to soluble biotinylated PT1 (**b**), PT2 (**c**), PT3 (**d**), PT4 (**e**) and PT5 (**f**) are shown. The apparent equilibrium dissociation constant (*K*_D_^app^) of each individual selected clone was determined by normalizing the mean fluorescence intensity from the binding signal to the median fluorescence intensity from the display signal (*y* axis), as a function of PT concentration (*x* axis). The indicated *K*_D_^app^ values are the results of three independent experiments and are presented as mean (dots) ± s.e.m. (bars); **g**) Plot reporting apparent binding affinity values of the 17 characterised yeast-encoded ‘one ring’ (red-coloured filled circles) and ‘two rings’ (blue-coloured filled circles) macrocyclic peptides to soluble biotinylated PT. The indicated values represent the means of at least three independent experiments presented as *K*_D_^app^ (nM).

Next, we compared the different macrocyclic peptide sequences selected for each PT and checked whether there was a correlation between the determined binding affinities and the topology or length of the isolated ligands (**Figure 5g** and **Supplementary figure 10**). Contrary to what one might intuitively expect, we often found that ‘one ring’ macrocyclic peptides bound the protein targets tighter than those with ‘two rings’. For example, the 9-amino acid ‘one ring’ macrocyclic peptides MP2.2 and MP2.3 appear to bind PT2 with *K*_D_^app^ values 10-fold lower (*K*_D_^app^ = 200 pM) or comparable (*K*_D_^app^ = 1.9 nM) to that of the longer 13-amino acid ‘two rings’ macrocyclic peptide MP2.1 (*K*_D_^app^= 2.1 nM), respectively (**Figure 5c**, **g** and **Supplementary figure 10**). Furthermore, the 9-amino acid ‘one ring’ macrocyclic peptide MP4.1 (*K*_D_^app^ = 41 nM) shows 16-fold higher binding affinity for PT4 than the similar length ‘two rings’ macrocyclic peptide MP4.4 (*K*_D_^app^ = 680 nM; **Figure 5e**, and **Supplementary figure 10**). Likewise, the 7-amino acid ‘one ring’ macrocyclic peptide MP5.1 (*K*_D_^app^ = 900 pM) bound 10-fold tighter PT5 than the 13-amino acid ‘two rings’ macrocyclic peptide MP5.5 (*K*_D_^app^ = 9.2 nM; **Figure 5f, g** and **Supplementary figure 10**). Loop length also does not always appear to be proportionally related to binding affinity. For example, the 7-amino acid ‘one ring’ macrocyclic peptide MP5.1 bound 4-fold better PT5 than the longer 9-amino acid ‘one ring’ macrocyclic peptide MP5.2 (*K*_D_^app^ = 3.7 nM; **Figure 5g** and **Supplementary figure 9**). Similarly, the short 7-amino acid MP5.3 (*K*_D_^app^ = 2.4 nM) and 9-amino acid MP5.4 (*K*_D_^app^ = 200 pM) ‘two rings’ macrocyclic peptides bound 3- and 46-fold tighter PT5 than the longer 13-amino acid bicyclic peptide MP5.5 (*K*_D_^app^ = 9.2 nM), respectively (**Figure 5g** and **Supplementary figure 9**).

To validate the *K*_D_^app^ values measured using yeast surface titrations, we determined the binding affinities of some macrocyclic peptides using a complementary technique such as surface plasmon resonance (SPR). While yeast surface titrations are performed using soluble biotinylated PT against the membrane-anchored macrocyclic peptides, the SPR measurements are executed using soluble macrocyclic peptides against chip-immobilized PTs (**Figure 6a**). This opposite orientation would allow us to rule out potential overestimation of binding affinity caused by undesirable multivalent binding phenomena, which can often occur on yeast when using biotinylated PTs that form multimers in solution. Toward this goal, three ‘one ring’ (MP1.1, MP2.2 and MP4.2) and two ‘two rings’ (MP3.2 and MP5.4) macrocyclic peptides that showed the highest binding affinities for their respective PTs were produced using solid phase peptide synthesis, cyclised, purified by reversed-phase high performance liquid chromatography, their molecular mass determined by electrospray ionization mass spectrometry (**Supplementary figure 11** – **13**) and their binding kinetics toward immobilised PTs assessed using SPR (**Figure 6b-d**, **Supplementary figure 14** and **Supplementary table 17**). The *K*_D_ values determined using SPR were on average comparable to those measured using yeast surface titrations (**Figure 6f**). A large discrepancy (∼500-fold) was observed only for the 9-amino acid ‘one ring’ macrocyclic peptide MP1.1 that appears to bind PT1 with a *K*_D_ value of 600 pM and 326 nM when using yeast surface titration and SPR, respectively (**Figure 6f**). Such difference is not surprising and may reflect the known tetrameric structure of PT1 interacting with multiple yeast-displayed macrocyclic peptides, thus confusing avidity effects with higher affinity.

**Figure 6.**
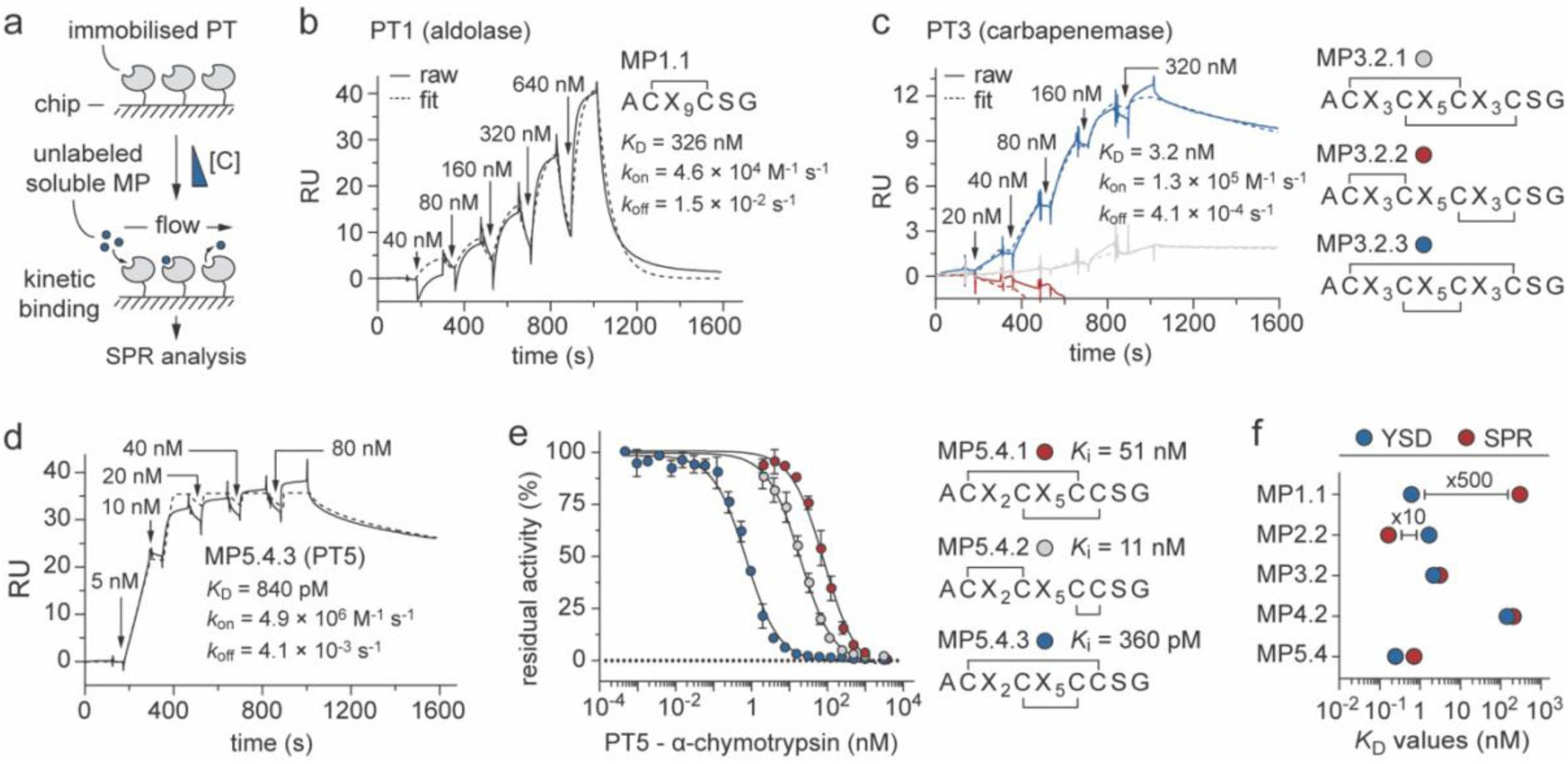
Determination of the binding affinities of soluble macrocyclic peptides against immobilised protein targets. **a**) Schematic representation of the binding affinity determination using surface plasmon resonance (SPR). Protein target (PT) covalently immobilised on the surface of a chip is incubated with varying concentrations of soluble chemically synthesised macrocyclic peptides (MPs). SPR sensorgram traces for the interaction of soluble macrocycle peptides MP1.1 (‘one ring’; **b**), MP3.2 (‘two rings’, three isomers: MP3.2.1, MP3.2.2 and MP3.2.3; **c**) and MP5.4 (‘two rings’, one isomer: MP5.4.3; **d**) with the immobilised biotinylated PT1, PT3 and PT5, respectively. Sensorgram traces presented with injection and flow fill steps removed were fitted with the 1:1 binding model. Kinetic constants *k*_on_ (association constant), *k*_off_ (dissociation constant) and *K*_D_ (equilibrium binding affinity) are presented as geometric mean average values. Raw data are shown as solid lines, while fitting curves are shown as dashed lines.; **e**) Residual activities of PT5 (α-chymotrypsin) measured at different concentrations of ‘two rings’ macrocyclic peptide isomers MP5.4.1, MP5.4.2 and MP5.4.3. The inhibitory activities of all macrocycle variants towards PT5 protease were determined at 25 °C and physiological pH (7.4) using the chromogenic N-Succinyl-Ala-Ala-Pro-Phe *p*-nitroanilide substrate at a concentration of 100 μM. The *K*_m_ value of PT5 protease (α-chymotrypsin) was determined by standard Michaelis-Menten kinetics and used in the calculation of the reported *K*_i_ values. The shown values are the means of three independent experiments. Data are presented as means (symbol). S.E., standard error; **f**) Plot reporting the *K*_D_ values determined using yeast surface titration (YSD; blue-coloured filled circles) and those measured using SPR (red-coloured filled circles).

In addition, SPR also allowed us to identify which of the three possible ‘two rings’ macrocyclic peptide isomers, potentially attainable from each sequence containing four cysteines, interacts tighter with the given PT. In the case of macrocyclic peptide MP3.2, out of the three possible isomers only MP3.2.3 bound PT3 with good affinity (*K*_D_ = 3.2 nM), while no or very weak interactions were detected for ‘two rings’ macrocyclic peptide isomers MP3.2.2 and MP3.2.1, respectively (**Figure 6c** and **Supplementary table 17**). Notably, the *K*_D_ value determined for MP3.2.3 using SPR (*K*_D_ = 3.2 nM) is similar to that measured using yeast surface titrations (*K*_D_ = 2.2 nM). Similarly, of the three isomers of macrocyclic peptide MP5.4, only MP5.4.3 bound PT5 with good affinity (*K*_D_ = 840 pM) while no or very weak interactions were detected for the ‘two rings’ macrocyclic peptide isomers MP5.4.1 and MP5.4.2, respectively (**Figure 6d**). To identify which MP5.4 isomer bound tighter to PT5 we employed a complementary approach and determined the inhibitory constants values (*K*_i_) of each chemically synthesised isomer (MP5.4.1, MP5.4.2 and MP5.4.3) by measuring the residual activity of enzyme PT5 using a colorimetric substrate at physiological pH and room temperature. The ‘two rings’ macrocyclic peptide MP5.4.3 showed a *K*_i_ value of 360 pM, approximately 30- and 140-fold better than those assessed for isomers MP5.4.2 (*K*_i_ = 11.3 nM) and MP5.4.1 (*K*_i_ = 50.8 nM), respectively (**Figure 6e** and **Supplementary table 18**). The good correlation between the determined *K*_i_ and *K*_D_ values further supports the effectiveness of yeast surface display technology for rapid and quantitative characterisation of individual macrocyclic peptide variants as cell-surface fusions.

Overall, our data revealed that macrocyclic peptides with fine binding affinities towards a panel of distinct target proteins can be rapidly identified even from low-diversity combinatorial libraries if screening is performed using a quantitative flow cytometry-based technology that allows continuous and precise monitoring of the evolution process.

### Selected yeast-encoded macrocyclic peptides exhibit fine binding specificities

To assess the extent of specificity of the selected macrocyclic peptides, we measured the binding of each yeast-displayed macrocyclic peptide ligand directly as cell surface fusions using flow cytometry and a panel of diverse proteins. We initially ruled out potential non-specific polyreactivity by assessing the binding of our macrocycle peptides toward the same highly diverse PTs used in our screening which share <10% sequence identity. No or very weak binding signals were detected for all macrocyclic peptides analysed, confirming their specificity toward the PT which they have been selected for (**Figure 7a**). Next, we assessed the binding of some selected macrocyclic peptides toward homologue proteins (PH) that share >30% sequence identity. We initially determined binding specificities of three different yeast-displayed macrocyclic peptides that were selected against PT2 (streptavidin) by titrating them into solutions with varying concentrations of two PHs of streptavidin namely strep-tactin (PH2.1, 98% identity) and neutravidin (PH2.2, 32% identify). No binding was detected for all three macrocyclic peptides MP2.1, MP2.2 and MP2.3 toward the most diverse PH2.2 while a 2- to 10-fold difference in affinity was observed for the highly similar PH2.1 (**Figure 7b**, **Supplementary figure 15** and **Supplementary table 19**).

**Figure 7.**
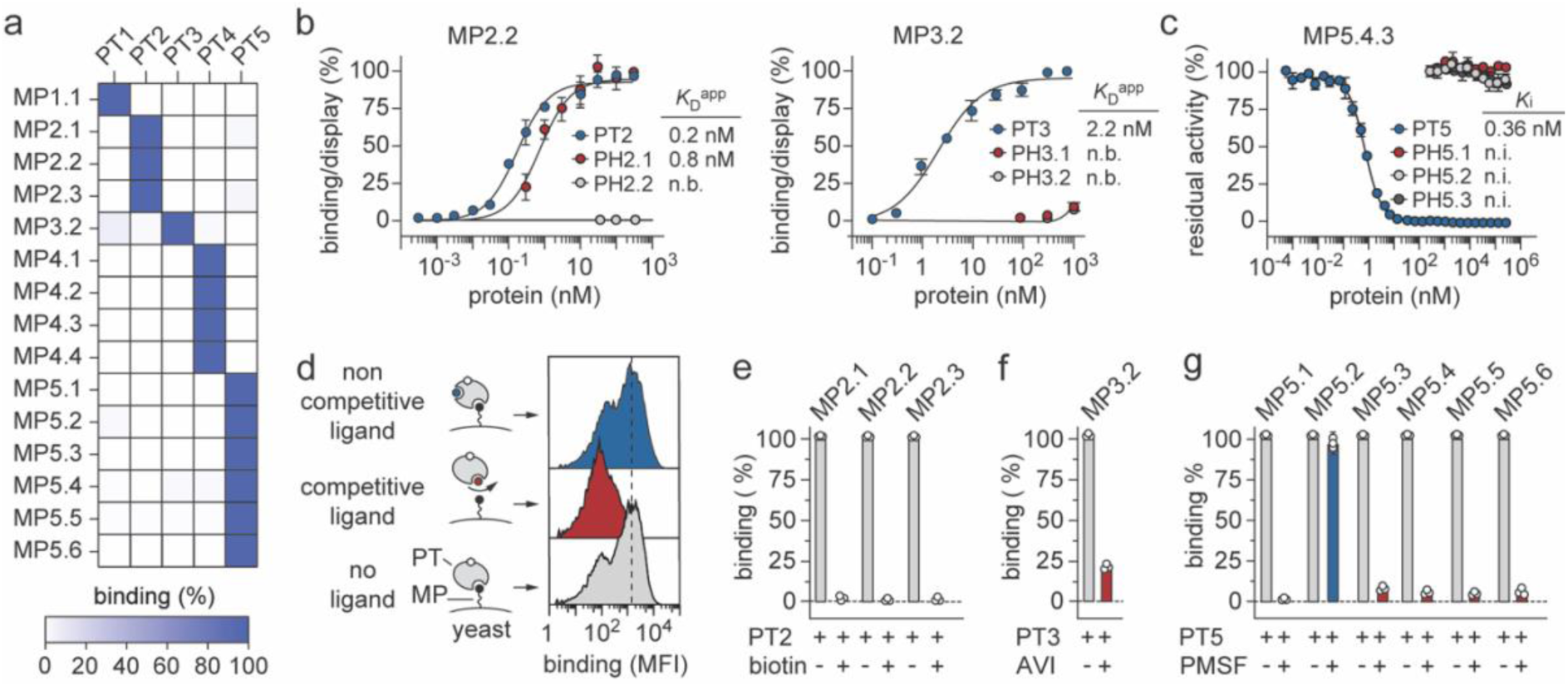
Determination of the binding specificities of yeast-encoded macrocyclic peptides. **a**) Heat map indicating the percentage of residual binding of all five diverse protein targets (PTs) against fifteen selected yeast-encoded macrocyclic peptides (MPs). Binding was assessed by flow cytometry using high concentration (1 μM) of soluble PTs against yeast-displayed MPs. Normalized binding/display signal intensities range from light to dark blue colours indicating low and high titres, respectively; **b**) Left, binding isotherms of yeast-displayed MP2.2 peptide towards streptavidin (PT2, blue), strep-tactin (PH2.1, red) and neutravidin (PH2.2, grey). Right, binding of yeast-displayed MP3.2 peptide towards carbapenemase GES-5 (PT3, blue), KPC-2 (PH3.1, red) and CTX-M-15 (PH3.2, grey). Apparent equilibrium binding affinity (*K*_D_ ^app^) values are reported. Data are presented as mean (dots) ± s.e.m. (bars). n.b., not binding; **c**) Residual activities of α-chymotrypsin (PT5, blue), trypsin (PH5.1, red), plasmin (PH5.2, light grey) and thrombin (PH5.3, dark grey) incubated with the ‘two rings’ macrocyclic peptide isomer MP5.4.3 were determined at 25 °C and physiological pH 7.4. The shown values are the means of three independent experiments. Data are presented as means (symbol) and S.E., standard error. n.i., not inhibition; **d**) Schematic representation of the competitive binding assay of yeast-encoded MPs for binding to their respective PT in the presence or absence of a known site-specific PT binding soluble ligand. Binding was assessed by flow cytometry-based assay. The determined residual fluorescence binding levels are indicated in percentage (%) and represent the mean values of three independent experiments. When known site-specific PT binding soluble ligand recognises the same PT site of MP, a decrease in fluorescence is expected (red plots; ‘competitive binding’). In the absence of known site-specific PT binding soluble ligand (grey plot; ‘no ligand’) or if it binds a PT site other than the one recognised by MP (blue plot; ‘non-competitive binding’), no decrease in fluorescence should be observed; **e**) Competitive binding assay of yeast-encoded macrocyclic peptides MP2.1, MP2.2 and MP2.3 for binding to PT2 (streptavidin) in the presence (+) or absence (-) of biotin; **f**) Competitive binding assay of yeast-encoded macrocyclic peptides MP3.2 for binding to PT3 (carbapenemase GES-5) in the presence (+) or absence (-) of the non-β-lactam β-lactamase avibactam (AVI); **g**) Competitive binding assay of yeast-encoded macrocyclic peptides MP5.1, MP5.2, MP5.3, MP5.4, MP5.5 and MP5.6 for binding to PT5 (bovine α-chymotrypsin) in the presence (+) or absence (-) of PMSF.

Interestingly, though the two macrocyclic peptides MP2.1 and MP2.3 retained a higher binding affinity for PT2 than PH2.1, macrocycle peptide MP2.2 bound PH2.1 4-fold more tightly (*K*_D_^app^ = 0.8 nM) than PT2 (*K*_D_^app^ = 0.2 nM; **Figure 7b**, and **Supplementary table 19**). Although not a striking difference, it is still surprising to observe that there can be up to a 10-fold variation in binding affinity between proteins that share >95% sequence identity, further highlighting the exquisite specificity of our yeast-encoded macrocyclic peptides. To additionally confirm the specificity of these interactions, we performed kinetic competition experiments using biotin, a well-characterized small molecule that binds with exceptionally high affinities (*K*_D_ = 10 fM) all three different PHs (**Figure 7d** and **Supplementary figure 16**). Exposure of yeast-displayed macrocyclic peptides to solutions of PT2 and PH2.1 pre-incubated with a molar excess of biotin resulted in a loss of binding for all three binders tested (**Figure 7e**). The ability of biotin to prevent the interaction of our macrocyclic peptides with both PT2 and PH2.1 also allows us to unveil the binding site. Similar exquisite specificity was observed when we determined the binding affinity of MP3.2, a yeast-displayed ‘two rings’ macrocyclic peptide selected against PT3, a carbapenemase GES-5 from *Klebsiella pneumoniae* ^36^ that was titrated into solutions with varying concentrations of two other class A β-lactamases, namely the carbapenemase KPC-2 ^37^ from *Klebsiella oxytoca* (PH3.1) and the extended-spectrum β-lactamase CTX-M-15 ^38^ from *Escherichia coli* (PH3.2; **Figure 7b**, **Supplementary figure 16** and **Supplementary table 20**). Further kinetic competition studies using a molar excess of avibactam, a non-β-lactam inhibitor of GES-5,^39^ revealed that the ‘two rings’ macrocyclic peptide MP3.2 recognises the same catalytic pocket (**Figure 7e**). To assess the specificity of MP5.4.3, a ‘two rings’ macrocyclic peptide that inhibits chymotrypsin with a *K*_i_ value of 360 pM, we determined its inhibition constants against a group of structurally and functionally related serine proteases, namely trypsin (PH5.1), plasmin (PH5.2) and thrombin (PH5.3). Notably, no inhibition of either PH5.1, PH5.2 or PH5.3 was detected even at the highest concentration tested (300 μM), revealing >100000-fold selectivity (**Figure 7c**, **Supplementary figure 16** and **Supplementary table 21**). To assess whether all six macrocyclic peptides (MP5.1 – MP5.6) identified against PT5 bound the same active site, we conducted binding experiments using a site-specific chemically modified PT5 variant. Exposure of yeast-displayed macrocyclic peptides to solutions of PT5 pre-incubated with phenylmethylsulfonyl fluoride (PMSF), a serine protease inhibitor known to irreversibly sulfonate the Oγ atom of the catalytic serine residue and therefore obliterating the active site of PT5, resulted in a loss of binding for five out of six macrocyclic peptides tested (**Figure 7g**). Similar results were obtained when MP5.2 and MP5.4 were exposed to solutions of PT5 pre-incubated with the bulkier aprotinin, a natural cyclic peptide serin protease inhibitor. MP5.2 retained a partial binding (75%) to chymotrypsin, while MP5.4 binding was completely inhibited (**Supplementary figure 15**).

Taken together, these data indicate that yeast-encoded macrocyclic peptides appear to be highly specific for the protein target they have been selected for. If present, cross-reactivity is often limited to highly similar proteins, while no binding occurs with low identity or unrelated ones. Further competition experiments not only confirmed the ability of our macrocyclic peptides to bind PTs with good specificity, but also highlighted the possibility of using the technology to rapidly ensure recognition of properly folded PTs as well as unveiled the binding site without the need of chemically synthesise and purify the selected ligand.

### Yeast-encoded macrocyclic peptide reveals large contact interface with a protein target

In order to investigate the binding mode of one yeast-encoded macrocyclic peptide to its target, we applied X-ray crystallography and determined the structure of PT5 (bovine α-chymotrypsin) in complex with the ‘two rings’ macrocyclic peptide MP5.4.3 at 2.42 Å maximum resolution (PDB identification code: 9F6H; **Figure 8a** and **Supplementary Table 22**). A single molecule of PT5 in complex with MP5.4.3 is present in the asymmetric unit. The overall structure of PT5 does not show any striking rearrangements of the main backbone (root mean square deviations of the Cα-atoms never exceed 0.5 Å) if compared to another crystal structure of PT5 belonging to the same space group that has been determined in the apo form (PDB identification code: 4CHA).

**Figure 8.**
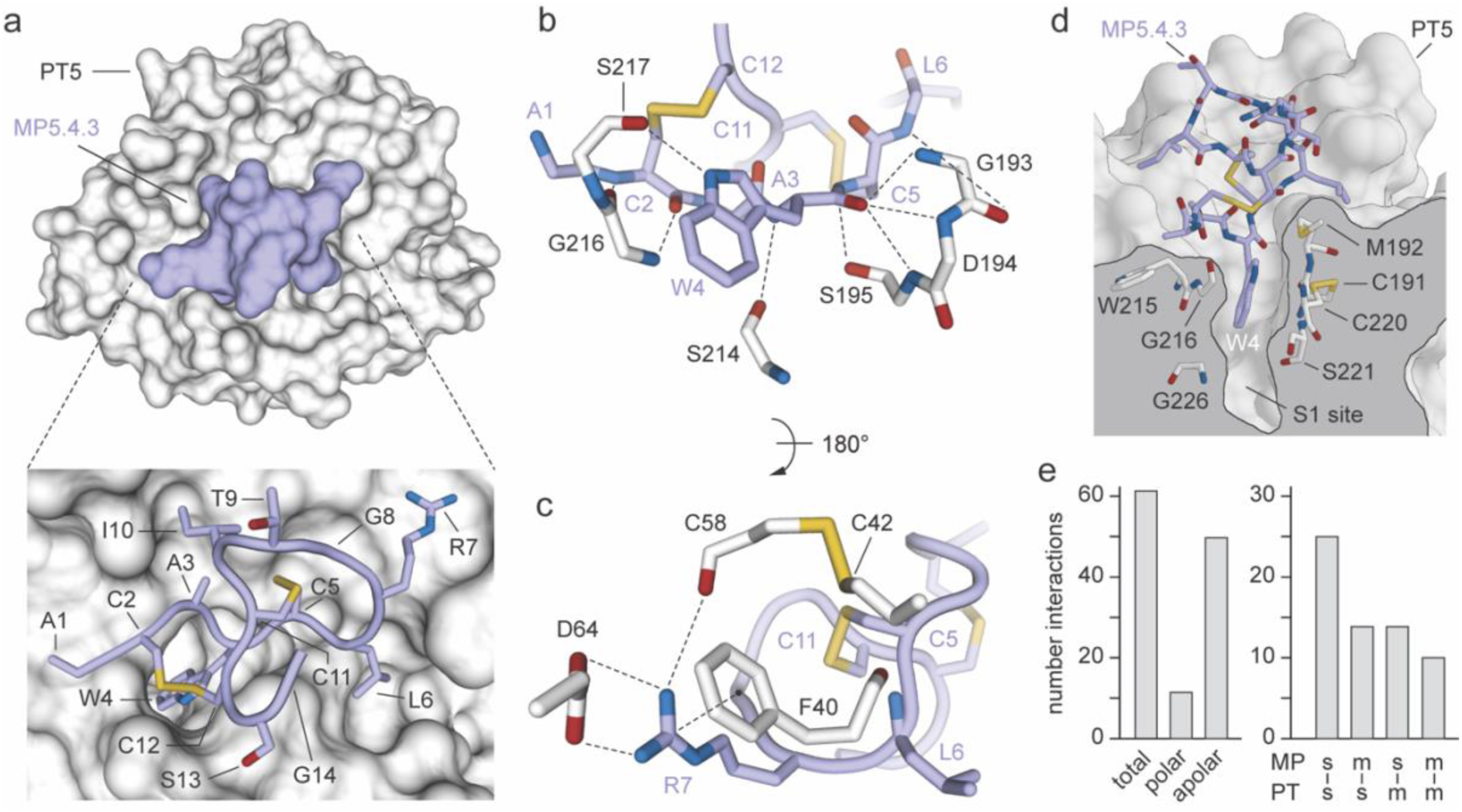
Crystal structure of α-chymotrypsin in complex with a ‘two rings’ macrocyclic peptide. **a**) Molecular surface representation of bovine α-chymotrypsin (PT5; grey) in complex with the ‘two rings’ macrocyclic peptide MP5.4.3 (light blue). A large surface (784 Å^2^) of PT5 is covered by MP5.4.3. Detailed view of the first (**b**, top) and second (**c**, bottom) ring of MP5.4.3 (light blue) bound to PT5 (white) shown in two orientations (180° rotation). The side chains of the residues are shown as sticks. Only the residues involved in key inter-molecular interactions (black dashed lines) are shown. Amino acid side chains are shown as sticks and coloured by atom type (carbon: light blue, oxygen: firebrick, nitrogen: deep blue, sulphur: yellow-orange); **d**) Zoomed-in view of the protein complex structure showing the residues of PT5 (white), known to define the primary substrate-binding pocket called S1 binding pocket (S189 – S195, S214 – C220 and P225 – Y228),^46,81^ surrounding and forming multiple interactions with the side-chain of Trp4 (W4) of MP5.4.3; **e**) Left, columns graph (grey) reporting the total number of polar (both direct and H_2_O-mediates) and non-polar intermolecular interactions established by PT5 with the ‘two rings’ macrocyclic peptide MP5.4.3. Right, columns graph (grey) reporting the total number of intermolecular interactions mediated by either side chain atoms (s) and/or main chain atoms (m) of MP5.4.3 (top) with either side chain atoms (s) and/or main chain atoms (m) of PT5 (bottom).

The electron density of MP5.4.3 is well defined, allowing an unambiguous assignment of side chain orientations (**Supplementary figure 17**). No classical secondary structure elements are found in the macrocyclic peptide. The sixty-three non-covalent intra-molecular interactions present are mainly mediated by main-chain to main-chain contacts and appear to confer structural rigidity to the macrocyclic peptide (**Figure 8a-c**, **Supplementary figure 17** and **Supplementary tables 23** – **25**). The ‘two rings’ macrocyclic peptide MP5.4.3 forms an extended structure that fits well into the cleft formed by the active site and the surrounding substrate pockets covering a total surface of 784 Å^2^. The high affinity and specificity of the bicyclic peptide can be explained by the large number of polar and non-polar inter-molecular contacts mediated by both catalytic (His57, Asp102, Ser195) and surrounding residues of PT5 (**Figure 8**, **Supplementary figure 17** and **Supplementary tables 23** – **25**). Both MP5.4.3 peptide rings interact directly with PT5. The first ring (residues from Cys2 to Cys12) is more rigid and forms a greater number of interactions than the second one (residues from Cys5 to Cys11) and hence contributes more to the overall binding (**Supplementary figure 17**). The residues of MP5.4.3 contacting PT5 through both main and side chain hydrogen bond interactions are eight (Ala1, Cys2, Trp4, Cys5, Leu6, Arg7 and Gly14). Most of these interactions are mediated by the main chain oxygen of Trp4 that establishes three hydrogen bonds with the main chain nitrogen of Gly193, Asp194 and Ser195 of PT5, with the first and last residue being key in the formation of the oxyanion hole (**Figure 8**, **Supplementary figure 17** and **Supplementary tables 24**, **25**). Additional important polar interactions are established by the side chain of Arg7 that forms a hydrogen bond with the main chain oxygen of Cys58, a salt bridge with the side chain of Asp64 and a cation-π interaction with the side chain of Phe41 (**Figure 8**, **Supplementary figure 17** and **Supplementary tables 24, 25**). The side chain of Trp4 occupies the primary specificity S1 hydrophobic pocket of chymotrypsin, as expected for natural substrates of this protease (**Figure 8d**, **Supplementary figure 18** and **Supplementary tables 24, 25**). However, the backbone does not have the required conformation for protease cleavage at this site, and the macrocyclic peptide is not cleaved by the protease. Notably, 37 out of 63 inter-molecular interactions are mediated by peptide main-chain atoms, thus explaining the weaker binding of the MP5.4.1 and MP5.4.2 isomers and further supporting the key role of the backbone conformation (**Figure 8e**, **Supplementary figure 17** and **Supplementary tables 24, 25**).

Overall, the crystal structure of the macrocyclic peptide MP5.4.3 in complex with the protease PT5 resembled those of protein complexes, with large interfaces of interaction, constrained peptide backbones and multiple directional hydrogen and electrostatic bonds from both loops of the peptide, leading to high binding affinity and exquisite selectivity.

## Conclusions

In summary, we have shown that yeast surface display, an *in vitro* directed evolution technology that has proved transformative for the discovery and engineering of multiple antibodies and diverse protein scaffolds, can also be successfully applied to generate macrocycle peptide ligands with desired binding properties. In this proof-of-concept study, we showed that large naïve combinatorial libraries encoding sequence and structural diverse disulfide-tethered macrocycle peptide ligands can be rapidly generated and effectively expressed on the surface of yeast cells. The size of the generated libraries (∼10^9^) is at the high end of those previously reported using yeast surface display but still a few orders of magnitude smaller than those attainable using other *in vitro* evolution techniques (>10^11^). The small size of the libraries has often been considered an Achilles’ heel of yeast surface display technology. However, while this drawback is central when engineering large proteins such as antibodies (>100 amino acids), it might become less critical for short peptide sequences (<10 amino acids) for which the diversity to be covered is smaller.

Next, we demonstrated the robustness and simplicity of our approach by rapidly isolating high binding affinity disulfide-tethered macrocycle peptide ligands against five highly diverse protein targets. By combining magnetic bead separations followed by multiple rounds of fluorescence-activated cell sorting we successfully identified macrocyclic peptide ligands against all five protein targets tested. Isolated macrocyclic peptide ligands showed different amino acid sequences, topologies and ring size distribution.

We demonstrated the ability of yeast surface display technology to enable rapid and efficient characterisation of isolated macrocyclic peptide ligands directly on the cell surface, thus eliminating the need for costly and time-consuming chemical synthesis and purification steps. All identified macrocyclic peptide ligands showed binding affinity values of less than 1 µM. Interestingly, for four out of the five protein targets tested we isolated at least one macrocyclic peptide ligand with binding affinity values below 10 nM. Importantly, yeast-encoded macrocyclic peptides showed high specificity exclusively for the protein target they have been selected for. These exquisite binding properties are remarkable considering that macrocyclic peptide ligands were rapidly isolated starting from relatively small naïve combinatorial libraries without the need of affinity maturation processes. Interestingly, there appears to be no correlation between the binding affinities determined and the topology or the size of the isolated macrocyclic peptide ligands. Contrary to what one might intuitively expect, we found that ‘one ring’ macrocyclic peptides can bind a protein target tighter than those with ‘two rings’. We speculate that, rather than the topology and the size of the macrocyclic peptide ligand, it is the structure of the target protein itself that determines the propensity and the strength of the binding achievable. This somewhat supports the importance of using multiple unbiased library designs to increase the chances of identifying macrocyclic peptide ligands with the desired binding properties.

Interestingly, by performing kinetic competition experiments in the presence of well-characterised soluble ligands with known binding affinities, we could also rapidly unveil the binding site of macrocyclic peptide ligands directly on the yeast cell surface, without the need for time-consuming and painstaking downstream synthesis and purification processes.

The use of sequence and structural distinct protein targets allowed us to better evaluate the capabilities of the technology as well as address some potential limitations that still exist. For instance, aware of the possible ‘avidity’ issues that could arise when using yeast surface display technology in the presence of multivalent soluble targets,^25,27,35^ we included multimeric protein targets in the screening and determined the binding affinities of the selected macrocyclic peptide ligands using opposite orientations and complementary techniques. For the tetrameric protein streptavidin, we observed no difference between the binding affinity values measured by yeast surface titrations (cell-anchored macrocyclic peptide against soluble protein target) and those determined using surface plasmon resonance (chip-anchored protein target against soluble macrocyclic peptide ligand), whereas for the tetrameric protein aldolase, we observed a ∼500-fold discrepancy. One possible explanation for such difference between aldolase and streptavidin is that the extent of ‘avidity’ effect depends not only on the number of identical binding sites present on the protein target to be detected, but also on the size of the protein itself and thus the distance and the density of the epitope that must be concomitantly recognised by different copies of macrocyclic peptide ligands displayed on the surface of yeast cells. Nevertheless, the binding affinity values measured for tetrameric protein aldolase using surface plasmon resonance are still in the nanomolar range. Moreover, we cannot exclude the possibility that avidity may actually prove advantageous when working with otherwise undetectable weak macrocyclic peptide ligands, especially during the first rounds of screening, when each member of the library is represented in small numbers. If necessary, multivalent binding can be overcome during FACS sorts by applying kinetic strategies in the presence of larger excess of a competitive unlabelled protein target able to interfere in a concentration-dependent manner.^24^ All these strategies, that have been previously developed and optimised for the engineering of different protein scaffolds and have contributed to make yeast display technology a versatile and highly successful tool, can now be rapidly recapitulated for the engineering of macrocyclic peptide ligands.

To better appreciate the ability of yeast display technology to efficiently isolate macrocyclic peptide ligands from naïve combinatorial libraries, we included in the screening two protein targets, streptavidin and α-chymotrypsin, toward which macrocyclic peptide ligands had previously been isolated using well-established *in vitro* directed evolution tools. Notably, the macrocyclic peptide ligands selected against streptavidin reported in this work have at least 10-fold better binding affinities than those previously discovered using phage and mRNA display tools and much larger library diversities (**Supplementary figure 6**).^40–44^ This discrepancy is even greater when comparing the binding affinities of macrocyclic peptide ligands selected against α-chymotrypsin with those previously isolated using phage display (**Supplementary figure 6**).^45^ Interestingly, the binding affinity values of some of the selected macrocyclic peptide ligands against α-chymotrypsin described in this work are in the pico-molar range and comparable to those of small protein-based inhibitors (**Supplementary figure 6**).^46,47^ Remarkably, despite their small size, the binding mode of macrocyclic peptide ligands to α-chymotrypsin resemble that of protein-based inhibitors, with large interfaces of interaction, constrained peptide backbones and multiple directional hydrogen and electrostatic bonds, thus explaining their exquisite binding affinity and selectivity (**Supplementary figure 18**).

Though the aim of this work was solely to demonstrate the ability of yeast surface display to efficiently generate and isolate high-affinity and specificity disulphide-bound macrocyclic peptide ligands from large, topologically diverse combinatorial libraries, yeast surface display can also be combined with other *in vitro* directed evolution tools to further harness the unique advantages of each, thus enabling previously unexplored applications. These include the evaluation and improvement of binding and stability properties and the rapid and fine epitope mapping. Moreover, based on the enormous success that yeast surface display technology in rapidly screening and assessing the biophysical properties of hundreds of computationally designed proteins,^28,48,49^ we envisage the tool to be suitable also for effectively characterising *in silico* developed genetically encoded macrocyclic peptide ligands.

Although the design of the herein described yeast-encoded disulfide-tethered macrocyclic peptides restrict their immediate application primarily to extracellular protein targets, this does not preclude that their binding affinity, selectivity, stability, and membrane permeability could be further enhanced and tailored by applying various biocompatible post-translational modifications, medicinal chemistry and rational design strategies.^50–54^ Moreover, the modular structure of peptide and the commercial availability of hundreds of amino acid building blocks simplifies the rapid development of macrocyclic peptides with desired properties. For instance, incorporation of cysteine reacting chemical linkers, N-methylated amino acids, D-amino acids and non-proteinogenic amino acids, N-terminal capping/acetylation, deamination, systematic truncations or extensions of N-termini or C-termini, N- to C-terminal cyclization and amide bond mimetics, are chemical modifications that have often proved key to transform disulfide-cyclized macrocyclic peptide leads into potent analogues with better drug-like properties.^3,55,56^

Although many challenges still remain, the herein described yeast surface display-based approach has the potential to enable facile and cost-effective development of target-tailored disulfide-tethered macrocyclic peptide molecules with desired binding properties that can be readily used in the form of either chemical molecules or as gene fusion products. In the future, it will be important to prove the efficacy and broad applicability of this technology to generate genetically encoded macrocyclic ligands against challenging protein targets and demonstrate their therapeutic uses *in vivo*.

## Materials and Methods

### Bioinformatic analysis of protein targets

The biochemical and biophysical properties of each protein target (PT; **Supplementary table 9**) were determined using different bioinformatic tools. The volume (Å^3^) of each PT was assessed using MOLEonline web interface.^57^ A probe radius of 5 Å and an interior threshold of 1.1 Å were applied. The solvent accessible surface area (ASA) of each PT was calculated using PISA software and a spherical probe of 1.5 Å radius.^58^ The surface charge distribution of each PT was calculated using PyMOL.^59^ The cavities volume and surface representation of each PT were determined using VEGAZZ software setting a minimum and maximum sphere radius of 3 and 6 Å, respectively.^60^ The composition of secondary structure of each PT was determined using PDBsum database.^61^ The three-dimensional structure of each PT was generated and rendered using PyMOL.^59^ Molecular weight (MW), isoelectric point (pI), and protein extinction coefficient (ε) were determined using ProtParam tool.^62^ The properties were determined using the three-dimensional structures of PTs in the absence of bound ligands (apo form; **Supplementary table 10**).

### Purification and chemical biotinylation of protein targets

Protein targets (PT) rabbit aldolase (PT1, GE-Healthcare, Chicago, IL, USA), *Streptomyces avidinii* streptavidin (PT2, Thermo Fisher Scientific, Dreieich, Germany), *Klebsiella pneumoniae* carbapenemase GES-5 (PT3, recombinantly produced),^36^ bovine carbonic anhydrase (PT4, Fluka, Darmstadt, Germany) and bovine α-chymotrypsin (PT5, P00766, Fluka, Darmstadt, Germany) were dissolved in 1X PBS pH 7.4 and further purified by size exclusion chromatography using either a Superdex 75 16/600 GL or a Superdex 200 16/600 GL column (Cytiva, Marlborough, MA, USA) equilibrated with buffer 1X PBS pH 7.4 and connected to an ÄKTA pure 25 M system (Cytiva, Marlborough, MA, USA). The fractions containing the expected monodisperse PT were pulled and further concentrated to a final concentration of 10 µM using 10000 Da MWCO Amicon ultrafiltration devices (Merck, Nottingham, UK) at 4000 g and 4 °C on a Heraeus Multifuge X1R centrifuge (Thermo Fisher Scientific, Dreieich, Germany). Reactive EZ-link sulfo-NHS-LC-biotin (Thermo Fisher Scientific, Dreieich, Germany) was dissolved in 1X PBS pH 7.4 to obtain a final concentration of 10 mM. Protein-biotin conjugates were prepared by incubating each PT, at a concentration of 10 µM in 1X PBS pH 7.4, with ten-fold molar excess of EZ-link sulfo-NHS-LC-biotin (100 µM) for 1 hr at room temperature. The reaction was quenched with 1 M Tris-HCl. Excess of unreacted or hydrolysed biotinylation reagent was removed by gel filtration using a HiPrep 26/10 desalting column (Cytiva, Marlborough, MA, USA) equilibrated with buffer 1X PBS pH 7.4 and connected to an ÄKTA pure 25 M system (Cytiva, Marlborough, MA, USA). The fractions containing the expected monodisperse PT were pulled and concentrated to a final concentration ranging from 10 μM to 50 μM using 10000 Da MWCO Amicon ultrafiltration devices (Merck, Nottingham, UK) at 4000 g and 4 °C on a Heraeus Multifuge X1R centrifuge (Thermo Fisher Scientific, Dreieich, Germany). Final PT concentrations were determined using a BioPhotometer D30 UV spectrophotometer (Eppendorf, Hamburg, Germany). Purified PTs were flash frozen in liquid nitrogen and stored at -80 °C. The monodisperse state of concentrated biotinylated PTs was confirmed by size exclusion chromatography using a Superdex 200 10/300 GL column (Cytiva, Marlborough, MA, USA) equilibrated with buffer 1X PBS pH 7.4 and connected to an ÄKTA pure 25 M system (Cytiva, Marlborough, MA, USA). Purified biotinylated PTs were eluted as a single peak at elution volumes (*V*_e_) that correspond to apparent molecular masses of 160 kDa for aldolase (PT1, tetramer), 50 kDa for streptavidin (PT2, tetramer) and about 25 – 30 kDa for monomeric carbapenemase GES-5 (PT3), carbonic anhydrase (PT4) and α-chymotrypsin (PT5).

### Generation of the yeast-encoded macrocyclic peptide naïve libraries

The yeast-encoded macrocyclic peptide naïve libraries were constructed using homologous recombination-based methods.^22^ Macrocyclic peptides were displayed on the surface of yeast as N-terminal fusion of the cysteine-free glycosylphosphatidylinositol (GPI) cell-surface anchor protein.^33^ Yeast surface display vector was based on a modified version of the pCT-CON backbone,^22^ here renamed pCT-GPI. This vector contains a DNA sequence encoding for a secretory leader sequence, a *Nhe*I restriction site, a sequence encoding for a long and flexible spacer (G_4_S)_3_, a sequence encoding for the hemagglutinin tag (HA; YPYDVPDYA) followed by a sequence encoding for a GPI cell-surface anchor protein which has been modified to include a silent *Bam*HI restriction site. The synthetic genes were codon optimized for expression in *Saccharomyces cerevisiae* cells and obtained from Integrated DNA Technologies (Coralville, IA, USA). Yeast-encoded macrocyclic peptide naïve libraries were created by inserting the DNA sequences encoding the random peptide sequences AC(X)*_n_*CSG (X = any amino acid, *n* = 7 or 9) and AC(X)*_n_*C(X)*_m_*CSG (X = any amino acid, *n* = 3, 6 or 9 and *m* = 9, 6 or 3), the flexible (G_4_S)_3_ linker, the HA tag and a N-terminal portion of the GPI protein into the pCT-GPI vector. The insert was created by appending the DNA encoding the random macrocyclic peptide sequences to the N-terminus of the (G_4_S)_3_ – HA – GPI encoding gene in a PCR reaction (30 cycles) using the forward degenerate primers F1, F2, F3, F4 and F5 and the universal reverse primer R (**Supplementary table 26**). The oligonucleotides were obtained from Integrated DNA Technologies (Coralville, Iowa, IA, USA). The pCT-GPI vector was used as DNA template. The PCR amplifications were performed in a reaction volume of 50 μL containing the oligonucleotides (500 nM each), dNTP mix (250 μM), DNA template (1 ng), 10X DreamTaq buffer (1X), 1 DreamTaq DNA polymerase (1.25 U, Thermo Fisher Scientific, Dreieich, Germany) and H_2_O mQ. The pCT-GPI vector was double digested using *Nhe*I-HF and *Bam*HI-HF restriction enzymes (New England Biolabs, Woburn, MA, USA). The linearized pCT-GPI vector and the PCR products were further purified using ethanol precipitation and concentrated to 1 µg/µL, combined together in a 1:50 molar vector to insert ratio and electroporated into freshly prepared EBY100 competent cells, where the full constructs are reassembled via homologous recombination.^22,63^ Transformed yeast cultures were recovered and expanded in SD-SCAA media. Small portions of transformed cells were serially diluted and titrated on SD-SCAA plates to assess the final library sizes.^22,63^ Library quality and diversity was further assessed by next generation sequencing.

### Sample preparation and analysis of next generation sequencing data

Transformed yeast cultures were grown to mid-log phase (OD_600_ = 2 – 3) in fresh SD-CAA media at 30 °C with shacking (250 rpm) and the DNA plasmid pools extracted using Zymoprep Yeast Plasmid Miniprep II kit following manufacture instructions (Zymo Research, Irvine, CA, USA).^22^ Amplicons for next generation sequencing (NGS) were generated performing two consecutive PCR reactions using yeast-extracted DNA plasmids as template. For each yeast-encoded macrocyclic peptide library a specific ten nucleotide barcode and an Illumina sequencing tag were used (**Supplementary table 11**). The oligonucleotide primers were obtained from Integrated DNA Technologies (Coralville, IA, USA). The first PCR was performed in a reaction volume of 30 µL containing 5X HF buffer (1X), DMSO (30% v/v), dNTPs (200 μM), NGS-F1 oligonucleotide (500 nM), NGS-R1 oligonucleotide (500 nM), Phusion high-fidelity DNA polymerase (New England Biolabs, Woburn, MA, USA), and yeast-extracted DNA plasmids (10 μL). The PCR amplicons (280 bp) were purified using the Clean and Concentrator PCR purification kit (Zymo Research, Irvine, CA, USA) and their quality verified on a 1.5% w/v agarose gel. The second PCR was performed in a reaction volume of 50 µL containing 5X HF buffer (1X), DMSO (30% v/v), dNTPs (200 μM), NGS-F2-X barcode oligonucleotide (500 nM, where X is a letter from A to I) and NGS-R2 oligonucleotide (500 nM), Phusion high-fidelity DNA polymerase (New England Biolabs, Woburn, MA, USA), and initial purified PCR products (10 ng). The PCR products were analysed on a 1% w/v agarose, gel-extracted and purified using Qiagen Gel Extraction Kit (Qiagen, Hilden, Germany). The final DNA concentration and quality were assessed using a BioPhotometer D30 UV spectrophotometer (Eppendorf, Hamburg, Germany). The amplicons were sequenced using a NovaSeq6000 (IGA Technology Services, Udine, Italy), yielding paired-end reads of 150 bp. Raw data were processed using MatLab scripts that have been adapted to the need of this study.^64^ The applied command lines and the codes are reported in **Supplementary information** and **Supplementary table 12**.

### Selection of yeast-encoded macrocyclic peptides

Selection of yeast-encoded macrocyclic peptide binders toward each protein target (PT) was performed using an amount of yeast cells at least ten-fold larger than *i*) the initial estimated naïve library size (ranging from 5 × 10^8^ to 2 × 10^9^ unique clones) or *ii*) the number of cells isolated from the previous round of either magnetic bead screening (MBs) or flow cytometry cell sorting (FACS). Each single yeast cell display naïve macrocyclic peptide library was grown separately in SD-CAA media at 30 °C with shacking (250 rpm) and surface expression of macrocyclic peptides induced in SG-CAA media for 16 hrs at 20 °C with shacking (250 rpm). Before positive selection, separately expanded and induced yeast populations were mixed ensuring to cover at least 10-fold the diversity of each library and subjected to two sequential cycles of negative selection using uncoated Dynabeads biotin binder magnetic beads (Thermo Fisher Scientific, Dreieich, Germany). Ten-fold diversity mixed library depleted of streptavidin-coated beads binders was following screened against highly diverse biotinylated PTs captured on MBs. Two iterative cycles of MB-based selections followed by four cycles of FACS were applied. Each cycle comprises growth of yeast cells, expression of the macrocyclic peptide on the surface of yeast cells, binding to the biotinylated PT, washing and expansion of the isolated bound yeast cells. For each cycle of MB-based selection, yeast cells displaying macrocyclic peptide binders (2 × 10^9^ cells/mL) were washed twice with ice-cold PBSA buffer (1X PBS pH 7.4 supplemented with 0.1% w/v bovine serum albumin fraction V), incubated at 4 °C for 1 hr with 50 pmol biotinylated PT immobilized on 4 × 10^6^ streptavidin-coated MBs, cells washed three times using ice-cold PBSA buffer, rescued with 5 mL of fresh SD-CAA medium and grown for 16 hr at 30 °C with shacking (250 rpm). The use of highly avid reagents such as MB saturated with diverse biotinylated PTs increases the likelihood of isolating low-affinity macrocyclic peptide binders from the naïve library by exploiting the multivalent interaction between yeast cells and the pre-loaded PT. For FACS selection, yeast cells displaying macrocyclic peptide binders were isolated using a two-colour labelling scheme based on fluorescent-conjugated detection reagents for expression of macrocyclic peptides on the surface of yeast cells (anti-HA epitope tag) and binding of the same macrocyclic peptides to PTs (anti-biotin) at recommended dilutions (**Supplementary table 27**). Secondary fluorescent-conjugated detection reagents for FACS were alternated to avoid enrichment of potential streptavidin or neutravidin binders. Sorting was performed on BD FACSAriaIII sorter instrument (BD Life Sciences, Franklin Lakes, NJ, USA) and data evaluated using FlowJo v.10.0.7 software (BD Life Sciences, Franklin Lakes, NJ, USA). After four cycles of iterative FACS-based selections, DNA plasmid was extracted from collected polyclonal yeast cells and sequenced.

### Sequencing analysis of selected yeast-encoded macrocycle peptide ligands

The identity of the selected macrocyclic peptide ligands was revealed by both Sanger and next generation sequencing (NGS). For Sanger sequencing, a fraction of each polyclonal yeast cell population collected after the fourth round of FACS sort were spread and grown on selective SD-CAA solid agar media at 30 °C. Single colonies were further picked and grown to mid-log phase (OD_600_ = 2–3) in fresh SD-CAA media at 30 °C with shacking (250 rpm). The DNA plasmid present within the yeast cells was extracted using Zymoprep Yeast Plasmid Miniprep II kit following manufacture instructions (Zymo Research, Irvine, CA, USA).^22^ Purified yeast-extracted DNA plasmid was then used to transform competent *E. coli* TOP10 cells. Bacteria-extracted DNA plasmids were analysed by Sanger sequencing (BMR Genomics, Padova, Italy) using the following oligonucleotide: 5’-GACTTGGAAGGTGACTTCG-3’. For NGS analysis (IGA Technology Services, Udine, Italy), a fraction of polyclonal yeast cell populations collected after the fourth round of FACS sort, was grown to mid-log phase (OD_600_ = 2–3) in fresh SD-CAA media at 30 °C with shacking (250 rpm) and the DNA plasmid pools extracted using Zymoprep Yeast Plasmid Miniprep II kit following manufacture instructions (Zymo Research, Irvine, CA, USA). Purified yeast-extracted DNA plasmid was then used as template to generate amplicons through two consecutive PCR reactions. For each selected yeast-encoded macrocyclic peptide population a specific ten nucleotide barcode and an Illumina sequencing tag were used. The amplicons were sequenced using a NovaSeq6000 (IGA Technology Services, Udine, Italy), yielding paired-end reads of 150 bp. Raw data were processed using command lines and the codes are reported in **Supplementary information**.

### Generation of unique yeast-encoded macrocycle peptide ligands identified using next generation sequencing

While binding properties of the thirteen most abundant macrocyclic peptide ligands identified using Sanger sequencing (MP1.1, MP2.1, MP2.2, MP3.1, MP3.2, MP4.1, MP4.4, MP5.1, MP5.2, MP5.4, MP5.5, MP5.6 and MP5.7) could be characterised directly as cell-surface fusions starting from the same single picked colonies of sorted yeast cells, those identified by NGS analysis (MP2.3, MP3.3, MP4.2, MP4.3, MP5.3 and MP5.5) required additional cloning steps. DNA constructs encoding NGS-derived macrocyclic peptide ligands fused to the N-terminus of the GPI cell-surface anchor were generated using homologous recombination-based methods. Briefly, DNA inserts encoding the macrocyclic peptide sequences were appended to the N-terminus of the (G_4_S)_3_ – HA – GPI encoding gene in a PCR reaction (30 cycles) using the pCT-GPI vector as template, forward and reverse oligonucleotides (**Supplementary table 15**) and the Phusion high-fidelity DNA polymerase (Thermo Fisher Scientific, Dreieich, Germany). The oligonucleotides were obtained from Integrated DNA Technologies (Coralville, IA, USA). The pCT-GPI vector was double digested using *Nhe*I-HF and *Bam*HI-HF restriction enzymes (New England Biolabs, Woburn, MA, USA). The linearized vector and the PCR products were further purified using ethanol precipitation, concentrated to 100 ng/µL, combined using DNA assembly methods (1:15 vector to insert molar ratio)^65^ and transformed into *E. coli* TOP10 competent cells. All DNA constructs encoding single macrocyclic peptide ligand were verified by Sanger sequencing and further used to transform new competent EBY100 cells using Frozen-EZ Yeast Transformation II Kit (Zymo Research, Irvine, CA, USA).

### Determination of equilibrium binding affinities using yeast surface display titrations

The apparent equilibrium dissociation constant (*K*_D_^app^) of individual selected macrocyclic peptide ligand towards each single PT was determined using yeast surface display titrations ^22^. Individual yeast colonies were inoculated into 5 mL SD-SCAA cultures, grown to mid-log phase (OD_600_ = 2–5) in SD-CAA media at 30 °C with shacking (250 rpm). Cells were induced in SG-CAA media for 16 hrs at 20 °C with shacking (250 rpm). The binding assays were conducted in 96-well conical V-bottom plates (Corning, Tewksbury, MA, USA) containing 2 × 10^5^ induced yeast cells per well. Yeast cells displaying macrocyclic peptide binders were incubated with varying concentrations of soluble biotinylated PT (ranging from 100 pM to 3 μM) overnight at 4 °C with gentle shaking (150 rpm). After PT incubation, yeast cells were pelleted (2200 g for 3 min at 4 °C) and washed twice with 200 μL ice-cold PBSA buffer. Labelling with secondary reagents such as streptavidin or neutravidin conjugated to DyLight dye was performed at recommended dilutions (**Supplementary table 27**). The level of cell surface expression of each macrocyclic peptide binder was estimated by incubating 2 × 10^5^ induced yeast cells with mouse anti-HA epitope tag (1:1000) antibody (Thermo Fischer Scientific, Dreieich, Germany). After incubation, yeast cells were pelleted (2200 g for 3 min at 4 °C) and washed twice with 200 μL ice-cold PBSA buffer. Labelling with secondary goat anti-mouse antibody conjugated to DyLight dye was performed at recommended dilutions (**Supplementary table 27**). The 96-well plates were run on a Attune NxT (Thermo Fischer Scientific, Dreieich, Germany) and data analysed using FlowJo v.10.0.7 software (BD Life Sciences, Franklin Lakes, NJ, USA). To ensure that the difference in binding was not due to variations of number of copies of macrocyclic peptides expressed on the surface of yeast cells, the mean fluorescence intensity (MFI_BIND_) from binding signal was normalized to the mean fluorescence intensity (MFI_DISP_) from display signal. The normalized (binding/display = MFI_BIND_/ MFI_DISP_) geometric mean fluorescence intensity as a function of PT concentration was used to determine the *K*_D_ ^app^ values for all clones of interest. The *K* ^app^ values were determined by fitting a one-site-specific binding curve on GraphPad Prism (GraphPad software Inc., San Diego, CA, USA). Reported values are the results of three independent experiments and are presented as mean (dots) ± s.e.m. (bars).

### Chemical synthesis of macrocyclic peptides

Linear peptides containing both cysteines protected with Trt(trityl), a free amine at the N-terminus and an amide at the C-terminus, were chemically synthetized by standard Fmoc (9-fluorenylmethoxycarbonyl) solid-phase peptide synthesis (SPPS).^66^ Fmoc-protected amino acids, Fmoc-rink amide MBHA resin (100 – 200 mesh, loading 0.4 – 0.9 mmol/g resin, 0.01 mmol scale), N,N-dimethylformamide (DMF) and anisole were purchased from Novabiochem (Darmstadt, Germany). Acetic anhydride, acetonitrile (ACN), formic acid, trifluoroacetic acid (TFA), octanedithiol (ODT), diethyl ether, dichloromethane (DCM), triisopropylsilane (TIS), piperidine, N-methylmorpholine (NMM), dimethyl sulfoxide (DMSO) and 2,6-Lutidine were purchased from Merck (Darmstadt, Germany). Thioanisole and 1,2-ethanedithiol (EDT) were purchased from FlukaChemie GmbH (Buchs, Switzerland). N-methylpirrolidone (NMP) was purchased from VWR (Pennsylvania, PA, USA). O-Benzotriazole-N,N,N’,N’-tetramethyl-uroniumhexafluoro-phosphate (HBTU) and Hexafluorophosphate Azabenzotriazole Tetramethyl Uronium (HATU) were purchased from ChemPep (Wellington, FL, USA). All chemicals were used as received without further purification. Peptides were synthetized using either a ResPepSLi or a MultiPepRSi automated peptide synthesiser (Intavis Bioanalytical Instruments, Köln, Germany). The deprotection step was carried out twice using a 20% v/v solution of Piperidine in DMF for 5 min. The amino acid coupling was carried out twice (60 min × 2) for each Fmoc-amino acid (2.15 eq., 0.18 M solution in DMF) using HATU/NMM coupling mixture (2 eq. 0.17 M / 4.7 eq. 0.4 M/ 4% v/v solution in DMF). The capping step was performed once using a solution of 5% v/v acetic anhydride and 6% v/v 2,6-Lutidine in DMF. Washes in between were performed using DMF (600 µL × 2). At the end of the synthetic process, washes were performed twice with DCM (600 µL × 2). The final peptides were deprotected and cleaved from the resin under reducing conditions using a TFA/H_2_O/thionalisole/anisole/ODT mixture (90/2.5/2.5/2.5/2.5% v/v) for 3 hrs at room temperature with shaking (300 rpm). The resin was removed by vacuum filtration and the peptides were precipitated with cold diethyl ether (50 mL) and centrifugation at 4000 g for 5 min at 4 ◦C either on a Heraeus Multifuge X1R centrifuge (Thermo Fisher Scientific, Waltham, MA, USA) or a Sigma 4-16KS centrifuge (Merck, Darmstadt, Germany) under inert atmosphere. The precipitated linear peptides were washed twice with cold diethyl ether (35 mL × 2) and cyclised. Macrocyclic peptides with ‘one ring’ were generated by dissolving crude linear peptides in 10% v/v DMSO and 90% v/v aqueous buffer (20 mM NH_4_HCO_3_, pH 8.0) at a final concentration of 500 µM. The reaction mixture was incubated at room temperature for 48 hrs with shacking (150 rpm). Macrocyclic peptides with ‘two rings’ were generated using two orthogonal cysteine protecting groups. Linear peptides containing one pair of cysteines protected with Mmt (Monomethoxytrityl) and the second pair with Dpm (1,2-Diphenylmaleyl), a free amine at the N-terminus and an amide at the C-terminus were chemically synthetized by standard Fmoc SPPS as described above. Deprotection of Mmt groups was conducted on resin under mild reducing conditions using DCM/TFA/TIS (93/2/5 % v/v) mixture with shaking (300 rpm) for 10 min at room temperature. The entire procedure was repeated five times. The on-resin Mmt-deprotected linear peptide was sequentially washed with 100% v/v DCM (3 mL × 3), 100% v/v MeOH (3 mL × 3), 100% v/v DCM (3 mL × 3) and 100% v/v DMF (3 mL × 3). The on-resin ‘one ring’ macrocyclic peptides (first disulfide bridge formation) were obtained by adding N-chlorosuccinimide (NCS, 2 eq. 0.04 M in DMF) with shaking (300 rpm) for 15 min at room temperature. The reaction solution was removed by vacuum filtration and the ‘one ring’ macrocyclic peptides washed three times with 100% v/v DMF (3 mL × 3) and 100% v/v DCM (3 mL × 3). ‘One ring’ macrocyclic peptides were further fully deprotected and cleaved from the resin under reducing conditions using TFA/H_2_O/TIS (95/2.5/2.5% v/v) mixture for 3 hrs at room temperature with shaking (300 rpm). The resin was removed by vacuum filtration and the ‘one ring’ macrocyclic peptides precipitated with cold diethyl ether (50 mL) and by centrifugation at 4000 g for 5 min at 4 ◦C on a Heraeus Multifuge X1R centrifuge (Thermo Fisher Scientific, Waltham, MA, USA) or Sigma 4-16KS centrifuge (Merck, Darmstadt, Germany) under inert atmosphere. The precipitated ‘one ring’ macrocyclic peptides were washed twice with cold diethyl ether (35 mL × 2) and the second cyclisation performed. ‘Two rings’ macrocyclic peptides (second disulfide bridge formation) were generated by dissolving ‘one ring’ macrocyclic peptides in 10% v/v DMSO and 90% v/v aqueous buffer (20 mM NH_4_HCO_3_, pH 8.0) at final concentration of 500 µM. The reaction mixture was incubated at room temperature for 48 hrs with shacking (150 rpm). Finally, the ‘two rings’ macrocyclic peptides were dissolved in H_2_O:ACN (1:1), freeze-dried and lyophilized on a LIO-5PDGT (5Pascal, Milan, Italy).

### Purification and characterisation of macrocyclic peptides

The macrocyclic peptides were purified by semi-preparative reversed-phase high performance liquid chromatography (RP-HPLC) using a C18 SymmetryPrep functionalized silica column (7 μm, 19 mm × 150 mm, Waters, Millford, MA, USA) connected to a Waters Delta Prep LC 4000 System equipped with a Waters 2489 dual λ absorbance detector, a Waters 600 pump and a PrepLC Controller (Waters, Millford, MA, USA).^67^ A flow rate of 4 mL/min and a linear gradient (30% to 70% in 25 min) with a mobile phase composed of eluant A (99.9% v/v H_2_O, 0.1% v/v TFA) and eluant B (99.9% v/v ACN and 0.1% v/v TFA) was applied. The purified peptides were freeze-dried. The purity and molecular mass of each macrocyclic peptide (linear, ‘one ring’ and ‘two rings’ forms) was determined by electrospray ionisation mass spectrometry (ESI–MS) performed either on a single quadrupole liquid chromatograph mass spectrometer LCMS-2020 (Shimadzu, Kyoto, Japan) or on an InfinityLab LC/MSD mass spectrometer coupled to a 1260 Infinity II LC system (Agilent Technologies, Santa Clara, CA, USA). Both systems operated with the standard ESI source and in the positive ionisation mode. Peptides were dissolved in DMSO:ACN:H_2_O (1:50:50) solution at a final concentration of 50 µM and run at flow rate of 1 mL/min with a linear gradient (10% - 100% v/v) of solvent B over 5 min (solvent A: 99.95% v/v H_2_O, 0.05% v/v formic acid; solvent B: 99.95% v/v ACN, 0.05% v/v formic acid). The reversed-phase HPLC column was a Nucleosil 100-5 C18 (5 μm, 125 mm × 4 mm; Macherey-Nagel, Dueren, Germany). Data were acquired, processed, and analysed using the Shimadzu LabSolutions software (Kyoto, Japan) and MestReNova (Mestrelab Research, Santiago de Compostela, Spain). Concentrations of macrocyclic peptides were determined using a BioPhotometer D30 UV spectrophotometer (Eppendorf, Hamburg, Germany).

### Determination of equilibrium binding affinities using surface plasmon resonance

The equilibrium binding constant (*K*_D_), association rate constant (‘on rate’, *k*_on_) and dissociation rate constant (‘off rate’, *k*_off_) for the interaction of chemical synthetised macrocyclic peptides with PTs were determined by surface plasmon resonance (SPR) using a Biacore 8K+ instrument (BIAcore Inc., Piscataway, NJ, USA). Protein targets PT1 (aldolase), PT2 (streptavidin), and PT5 (α-chymotrypsin) were diluted in 10 mM sodium acetate (NaAc) pH 5.2 at 200 µg/mL, 100 µg/mL and 30 µg/mL final concentration, respectively. Protein target PT3 (carbapenemase GES-5 from *Klebsiella pneumoniae*) and PT4 (carbonic anhydrase) were instead diluted in 10 mM NaAc pH 4.5 at 10 µg/mL final concentration. All PTs were captured on a series S sensor chip CM5 (Cytiva, Marlborough, US) using amine coupling in 1X PBS pH 7.4 and 0.005% v/v Tween-20. The coupling was performed at 25 °C and a continuous flow rate of 10 µL/min till a protein immobilization level of approximately 1000 resonance units (RUs) per flow cell was reached. The coupling involved four steps: *i*) injection of 70 µL of 0.4 M EDC for 420 s, *ii*) injection of 70 µL of 0.1 M NHS for 420 s, *iii*) injection of 70 µL of PT for 420 s and *iv*) injection of 83 µL of 1 M ethanolamine-HCl pH 8.5 for 500 s. Binding experiments with multiple two-fold dilution of each macrocyclic peptide were performed in running buffer (1X PBS supplemented with 0.005% v/v Tween-20 and 0.2% v/v DMSO) at 25 °C and a continuous flow rate of 30 µL/min. Serial 20 – 50 µL injections of two-fold dilution of the macrocyclic peptides were applied through the sensor surface with a continuous flow rate of 10 µL/min for a time period ranging from 120 – 320 s and with a dissociation time of 600 s. The bound macrocyclic peptides were allowed to dissociate for 120 s before a further analysis was performed. A solvent correction of four points (3.0%, 2.5%, 2.0 %, 1.5% v/v DMSO in 1X PBS supplemented with 0.005% v/v Tween-20) was performed before and after each analysis. In all experiments an untreated flow cell without PT protein was used as reference to correct binding response for bulk refractive index changes and unspecific binding. The association rate constant (*k*_on_) and the dissociation rate constant (*k*_off_) were determined by global fitting mode. Data were fitted to a Langmuir binding model assuming stoichiometric (1:1) interactions by using Biacore 8K Evaluation software (Piscataway, NJ, USA). The equilibrium binding constant values were calculated as the ratio of *k*_off_ to *k*_on_ (*K*_D_ = *k*_off_/*k*_on_). The *K*_D_ were also determined by plotting binding responses in the steady-state region of the sensorgram (R_eq_, 5 s before injection ends with a window of 5 s) versus analyte peptide concentration (*C*). The data were analysed by using Biacore 8K Evaluation software with the predefined evaluation methods LMW multi-cycle or LMW single-cycle kinetics.

### Determination of inhibitory activity of macrocyclic peptides

The inhibitory activity of macrocyclic peptides MP5.4.1, MP5.4.2 and MP5.4.3 was assessed by monitoring the residual activity of enzyme bovine α-chymotrypsin (PT5) in the presence of a chromogenic substrate and different concentrations of inhibitor macrocyclic peptides. The activity assay was performed by incubating 0.5 nM PT5 with 100 μM chromogenic substrate N-Succinyl-Ala-Ala-Pro-Phe *p*-nitroanilide (100 μM; Merck, Darmstadt, Germany) and two-fold macrocyclic peptide dilutions ranging from 0.0005 to 2000 nM. All reagents were diluted in 10 mM Tris-Cl, pH 7.4, 150 mM NaCl, 10 mM MgCl_2_, 1 mM CaCl_2_, 1% w/v BSA and 0.1% v/v Triton-X100 buffer. The measurements were performed on a Tecan Infinite 200 PRO microplate reader (Tecan Trading AG, Switzerland) using standard flat bottom 96-well plate (Thermo Fisher Scientific, Dreieich, Germany). The enzymatic reactions were performed at 25 °C for 1 hr, under shacking with an absorbance wavelength of 410 nm. The initial velocities were monitored as changes in absorbance intensity. The sigmoidal curves were fitted to the data using the following non-linear regression equation for the inhibitory dose-response curves with variable slope (1):

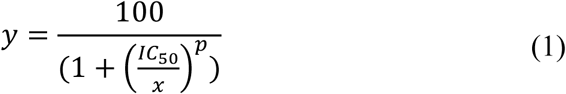

where *x* is the peptide concentration, *y* is the residual percentage of protease activity and *p* is the hill slope. Half maximum inhibitory concentration (IC_50_) values were derived from the fitted curves from GraphPad Prism 8 8.0.0. software (GraphPad software Inc., San Diego, CA, USA). The final inhibitory constant (*K*_i_) was subsequently determined using the Cheng-Prusoff equation (2):

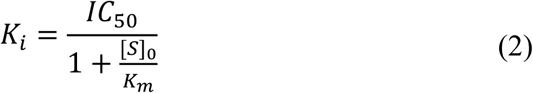

where *K*_m_ (114 μM) is the Michaelis constant for the hydrolysis of substrate N-Succinyl-Ala-Ala-Pro-Phe *p*-nitroanilide catalysed by PT5 which has been determined by standard Michaelis-Menten equation. Values were determined using either OriginPro 8G software (OriginLab Corporation, Northampton, MA, USA) or GraphPad Prism 8.0.0 software (GraphPad software Inc., San Diego, CA, USA). Reported values are the results of three independent experiments and are presented as mean (dots) ± s.e.m. (bars).

### Determination of binding specificity of selected macrocyclic peptides

To assess the specificity of macrocyclic peptides their equilibrium dissociation constants and/or inhibitory activities toward proteins with different extent of sequence and structural identity were evaluated. The apparent equilibrium dissociation constants (*K*_D_^app^) were determined using yeast surface display titrations combined to flow cytometry. The binding assays were conducted in 96-well conical V-bottom plates (Corning, Tewksbury, MA, USA) containing 2 × 10^5^ induced yeast cells per well. Yeast cells displaying macrocyclic peptides (MP1.1, MP2.2 and MP2.3) selected against DyLight 650 conjugated streptavidin (PT2) were incubated with varying concentrations of soluble DyLight 650 conjugated strep-tactin (PH2.1; concentrations ranging from 0.1 to 300 nM) and DyLight 650 conjugated neutravidin (PH2.2; concentrations ranging from 30 to 300 nM) for 1 hr at 4 °C with shaking (150 rpm). Similarly, yeast cells displaying macrocyclic peptide (MP3.2) selected against the carbapenemase GES-5 from *Klebsiella pneumoniae* (PT3) were incubated with varying concentrations of soluble biotinylated carbapenemase KPC-2 from *Klebsiella oxytoca* (PH3.1; concentrations ranging from 100 to 1000 nM) and the extended-spectrum β-lactamase CTX-M-15 from *Escherichia coli* (PH3.2; concentrations ranging from 100 to 1000 nM) for 1 hr at room temperature with shaking (150 rpm). After primary incubation, cells were pelleted (2200 g for 3 min at 4 °C) and washed twice with 200 μL of ice-cold PBSA buffer. In the case of biotinylated PTs, secondary labelling was performed with Neutravidin DyLight 650 at recommended dilution (**Supplementary table 27**). The 96-well plates were run on a Attune NxT (Thermo Fischer Scientific, Dreieich, Germany) and data analysed using FlowJo v.10.0.7 software (BD Life Sciences, Franklin Lakes, NJ, USA). To ensure that the difference in binding was not due to variations of number of copies of macrocyclic peptides expressed on the surface of yeast cell, the mean fluorescence intensity (MFI_BIND_) from binding signal was normalized to the mean fluorescence intensity (MFI_DISP_) from display signal. The normalized (binding/display = MFI_BIND_/MFI_DISP_) geometric mean fluorescence intensity as a function of PT concentration was used to determine the *K*_D_^app^ values for all clones of interest. The *K*_D_^app^ values were determined by fitting a one-site-specific binding curve on GraphPad Prism (GraphPad software Inc., San Diego, CA, USA). Reported values are the results of three independent experiments and are presented as mean (dots) ± s.e.m. (bars). The inhibitory activities (*K*_i_) were assessed by monitoring the residual activity of different enzymes in the presence of either chromogenic or a fluorogenic substrate and different concentrations of inhibitor macrocyclic peptides. Residual activities were measured in 150 μL volume of buffer containing 10 mM Tris-Cl, pH 7.4, 150 mM NaCl, 10 mM MgCl2, 1 mM CaCl2, 0.1% w/v BSA, 0.01% v/v Triton-X100 and 5% v/v DMSO. Final concentrations of serine proteases were the following: 0.5 nM bovine α-chymotrypsin (PT5), 6 nM human trypsin (PH5.1; hTRYP from Molecular Innovations, Novi, MI, USA), 0.05 nM human plasmin (PH5.2; hPLM from Molecular Innovations, Novi, MI, USA) and 10 nM human thrombin (PH5.3; IHT from Molecular Innovations, Novi, MI, USA). Two-fold dilutions of macrocyclic peptide inhibitor MP5.4.3 were prepared ranging from 0.25 to 256 µM for all the proteases. For PT5 additional two-fold dilutions of MP5.4.3 inhibitor was prepared ranging from 0.47 pM to 125 nM. For the determination of the *K*_i_ inhibitory constant of macrocyclic peptide inhibitor MP5.4.3 against PT5 we used chromogenic substrate N-Succinyl-Ala-Ala-Pro-Phe *p*-nitroanilide (100 μM; Merck, Darmstadt, Germany) at final concentration of 100 μM. For human trypsin (PH5.1) and human thrombin (PH5.3) we used the fluorogenic substrate Z-Gly-Gly-Arg-AMC (Bachem, Bubendorf, Switzerland), while for human plasmin (PH5.2) we used the fluorogenic substrate H-D-Val-Leu-Lys-AMC (Bachem, Bubendorf, Switzerland). All fluorogenic substrates were used at final concentration of 50 μM. The initial velocities were monitored as changes in absorbance or fluorescence intensity on a Tecan microplate reader (Tecan infinite 200 pro, Tecan Trading AG, Switzerland) using either standard flat bottom 96-well plate or black microfluor 96-well plate Nunc MicroWell (Thermo Fisher Scientific, Dreieich, Germany). In the case of PT5, the enzymatic reactions were performed at 25 °C for 1 hr, under shacking with an absorbance wavelength of 410 nm, whereas in the case of human PH5.1, PH5.2 and PH5.3 the enzymatic reactions were performed at 25 °C for 1 hr, under shacking with an excitation wavelength of 355 nm and an emission recording at 460 nm. Apparent inhibitory constants *K*_i_^app^ values were determined by non-linear regression analyses of V_i_/V_0_ versus [I]_0_ using equation (1). The final *K*_i_s were subsequently determined by correcting for the competitive effect of the substrate [S]_0_ using equation (2). The kinetic constants *K*_m_s for the hydrolysis of chromogenic or fluorogenic substrate, catalysed by each protease, were determined by standard Michaelis-Menten equation (**Supplementary table 21**).^66^ Values were determined using either OriginPro 8G software (OriginLab Corporation, Northampton, MA, USA) or GraphPad Prism 8.0.0 software (GraphPad software Inc., San Diego, CA, USA). Reported values are the results of three independent experiments and are presented as mean (dots) ± s.e.m. (bars).

### Competitive binding assay on yeast cells

A competitive flow cytometry-based binding assay was performed to rapidly identify the binding site of some selected yeast displayed macrocyclic peptides. The binding assays were conducted in 96-well conical V-bottom plates (Corning, Tewksbury, MA, USA) containing 2 × 10^5^ induced yeast cells per well. Yeast cells displaying the desired macrocyclic peptide were incubated for 1 hr at 4 °C with gentle shacking with a concentration 10-fold higher their *K*_D_^app^ values of soluble protein target (PT) alone or the PT pre-complexed with at least 100-fold molar excess of a well-known site-specific PT-binding soluble ligand. Competitive binding assay of yeast-encoded macrocyclic peptides MP2.1, MP2.2 and MP2.3 for binding to streptavidin (PT2) were conducted in the presence or absence of biotin (Merck, Darmstadt, Germany). Competitive binding assay of yeast-encoded macrocyclic peptide MP3.2 for binding to carbapenemase GES-5 from *Klebsiella pneumoniae* (PT3) were conducted in the presence or absence of the non-β-lactam β-lactamase inhibitor avibactam (Merck, Darmstadt, Germany). Competitive binding assay of yeast-encoded macrocyclic peptides MP5.1, MP5.2, MP5.3, MP5.4, MP5.5 and MP5.6 for binding to bovine α-chymotrypsin (PT5) were conducted in the presence or absence of phenylmethylsulfonyl fluoride (Merck, Darmstadt, Germany) and aprotinin (Cytiva, Marlborough, USA). After the incubation, cells were pelleted (2200 g for 3 min at 4 °C) and washed twice with 200 μL ice-cold PBSA buffer. The 96-well plates were run on a Attune NxT (Thermo Fischer Scientific, Dreieich, Germany) and data analysed using FlowJo v.10.0.7 software (BD Life Sciences, Franklin Lakes, NJ USA). The binding (MFI) values were normalized to the value obtained in the absence of soluble ligand providing us with a percentage value, ranging from 0 to 100 %, that corresponds to the residual binding observed upon incubation with the known site-specific PT-binding soluble ligand. Reported values are the results of three independent experiments and are presented as mean (dots) ± s.e.m. (bars).

### Crystallization and structure determination of a protein target in complex with a macrocyclic peptide

Lyophilized α-chymotrypsin from bovine pancreas (UniProt ID: P00766) was purchased from Fluka (Burlington, MA, USA), dissolved in 50 mM MES, pH 6.0 and further purified by size exclusion chromatography using a HiLoad 26/600 Superdex 200 prep grade column (Cytiva, Marlborough, MA, USA) pre-equilibrated with the same buffer and connected to an ÄKTA pure 25 M system (Cytiva, Marlborough, MA, USA). The fractions containing the monodisperse protein were pooled and further concentrated by ultrafiltration using 10000 Da MWCO Amicon ultrafiltration devices (Merck Life Science, Darmstadt, Germany) at 4000 g and 4 ^◦^C on a Heraeus Multifuge X1R centrifuge (Thermo Fisher Scientific, Waltham, MA, USA) to a final value of 25 mg/mL (1 mM). The protein concentration was determined using a BioPhotometer D30 UV spectrophotometer (Eppendorf, Hamburg, Germany). Crystallization trials of α-chymotrypsin in complex with macrocyclic peptide MP5.4.3 were carried out in SWISSCI MRC 96-well crystallization plates (Hampton Research, Aliso Viejo, CA, US) using the isothermal (293 K) sitting-drop vapor diffusion method and the PACT premier, LMB and Morpheus II screening kits (Molecular Dimensions Ltd., Sheffield, UK). Formation of protein-peptide complex was induced by incubating α-chymotrypsin (1 mM) with the macrocyclic peptide MP5.4.3 (1.5 mM, 5% v/v DMSO) for 2 hrs at 25 ^◦^C. Droplets of 0.6 μL volume (0.3 μL of protein-peptide complex in the presence of 5% v/v DMSO and 0.3 μL of reservoir solution) were dispensed by an Oryx 8 crystallization robot (Douglas Instruments Ltd., Berkshire, UK) and equilibrated against 70 μL of reservoir solution. Best crystals were obtained in 48 – 72 hrs using the following precipitant agent: 70% v/v MPD, 0.1 M HEPES, pH 7.5. For X-ray data collection, crystals were soaked in a solution of 20% v/v ethylene glycol in the precipitant buffer, mounted on LithoLoops (Molecular Dimensions Ltd, Suffolk, UK) and flash-frozen in liquid nitrogen. X-ray diffraction data were collected at the ID30-A beamline of the European Synchrotron Radiation Facility (ESRF, Grenoble, France). Crystals belong to the *P*6_1_ space group. The asymmetric unit contains 1 α-chymotrypsin molecule and a solvent content of 56% of the crystal volume. Reflections were indexed and integrated by the GrenADeS automated processing pipeline,^68^ merged and scaled by AIMLESS ^69^ in the CCP4i2 crystallographic suite.^70^ Phases were determined by molecular replacement with Molrep ^71^ using the PDB entry 4CHA as a template,^72^ via the CCP4i2 interface.^70^ Refinement of the protein model was carried out manually by Coot ^73^ and automatically by Refmac5 ^74^ and PDB_REDO.^75^ The peptide ligand was parametrized by eLBOW ^76^ via the PHENIX interface.^77^ The presence of macrocyclic peptide MP5.4.3 in α-chymotrypsin active site was confirmed by omit map at 2.5 σ. Since the first cycles of refinement, the electron density corresponding to the bound macrocyclic peptide MP5.4.3 was clearly visible in the electron density map and the peptide model was built in a well-defined unique conformation. The final model of the complex contains 1778 protein atoms, 97 macrocyclic peptide atoms and 41 water molecules. The final crystallographic *R* factor is 0.213 (*R*_free_ 0.28). Geometrical parameters of the model are as expected or better for this resolution. Buried surface calculations were performed using PDBePISA.^58,78^ Intra-molecular and inter-molecular interactions were analysed by LIGPLOT+ ^79^ and PLIP.^80^ All figures were made with PyMOL.^59^ The structure of α-chymotrypsin in complex with macrocyclic peptide MP5.4.3 has been deposited in the Protein Data Bank (PDB) under identification code 9F6H.

## Data availability

The data that support the findings of this study are present in the paper and/or the Supplementary information. Additional data related to this study are available from the authors upon request.

## Acknowledgements

The authors would like to thank Dr. Giuseppe Borsato and Dr. Alessandro Bonetto for technical assistance with peptide purification and mass spectrometry analysis. We are grateful to Mr. Federico Cusinato of the Department of Pharmaceutical and Pharmacological Sciences at the University of Padova for technical assistance during fluorescence activated cell sorting experiments. We thank Dr. Kelvin Lau at protein production and structure core facility at EPFL for technical assistance during surface plasmon resonance experiment. The authors would like to thank Dr. Gordon Leonard and the staff of ID30A-3 beamline of the European Synchrotron Radiation Facility (ESRF, Grenoble, France) for assistance with crystal testing and data collection. We are grateful to all the group members for helpful discussions and for critical reading of this manuscript.

## Author contribution

S.L. and A.A. conceived the study; S.L. generated the libraries; S.L. and Y.M. performed the screening, bioinformatic analyses and determined the binding affinities and specificities; Z.R. performed competition studies; S.L., Y.M., L.F.S. and Y.X. synthesised and purified macrocyclic peptides; S.L. and E.W. performed surface plasmon resonance experiments; F.V. and L.C. solved and analysed the x-ray structure; M.S: supervised bioinformatic analyses; S.C., A.S. and C.H. supervised chemical synthesis, modification and purification. All authors analysed the data, discussed the results, and wrote the manuscript.

