## Supplementary Tables and Figures for "Screening macrocyclic peptide libraries by yeast display allows control of selection process and affinity ranking"

| naïve library | | amino acid sequence design | sequence diversity | |
|---|---|---|---|---|
| | | | theoretical | experimental |
| **1** | $CX_7C$ | ACXXXXXXXCSG | $1.3 \times 10^9$ | $4.0 \times 10^8$ |
| **2** | $CX_9C$ | ACXXXXXXXXXCSG | $5.1 \times 10^{11}$ | $2.0 \times 10^9$ |
| **3** | $CX_3CX_9C$ | ACXXXCXXXXXXXXXCSG | $4.1 \times 10^{15}$ | $6.0 \times 10^8$ |
| **4** | $CX_6CX_6C$ | ACXXXXXXCXXXXXXCSG | $4.1 \times 10^{15}$ | $3.0 \times 10^8$ |
| **5** | $CX_9CX_3C$ | ACXXXXXXXXXCXXXCSG | $4.1 \times 10^{15}$ | $5.0 \times 10^8$ |

**Supplementary table 1**. Sequence design and diversity of yeast-encoded naïve macrocycle peptide libraries. Fixed cysteine residues (C) are underlined. The theoretical (left) and experimentally determined (right) sequence diversity of each library are reported.

| naïve library | | DNA sequence (5' – 3') |
|---|---|---|
| **1** | $CX_7C$ | GCTTGTNNKNNKNNKNNKNNKNNKNNKTGTTCCGGT |
| **2** | $CX_9C$ | GCTTGTNNKNNKNNKNNKNNKNNKNNKNNKNNKTGTTCCGGT |
| **3** | $CX_3CX_9C$ | GCTTGTNNKNNKNNKTGTNNKNNKNNKNNKNNKNNKNNKNNKNNKTGTTCCGGT |
| **4** | $CX_6CX_6C$ | GCTTGTNNKNNKNNKNNKNNKNNKTGTNNKNNKNNKNNKNNKNNKTGTTCCGGT |
| **5** | $CX_9CX_3C$ | GCTTGTNNKNNKNNKNNKNNKNNKNNKNNKNNKTGTNNKNNKNNKTGTTCCGGT |

**Supplementary table 2**. DNA sequences (5' to 3') designed to generate the naïve macrocycle peptide libraries. Random amino acid positions (X) in all libraries are encoded by 'NNK' codons. Fixed cysteine residues (C) are encoded by the underlined 'TGT' codon. The 'GCT', 'TCC' and 'GGT' codons encode for fixed amino acids alanine (A), serine (S) and glycine (G), respectively.

| naïve library | | frequency (%) additional or missing cysteine residues | | | |
|---|---|---|---|---|---|
| | | - 1 cys | + 0 cys | + 1 cys | + 2 cys |
| **1** | $CX_7C$ | 0% | 74% | 22% | 4% |
| **2** | $CX_9C$ | 0% | 65% | 28% | 7% |
| **3** | $CX_3CX_9C$ | 2% | 60% | 30% | 8% |
| **4** | $CX_6CX_6C$ | 2% | 60% | 30% | 8% |
| **5** | $CX_9CX_3C$ | 2% | 60% | 30% | 8% |

**Supplementary table 3**. Theoretical probability to identify additional or fewer cysteine residues (C) in the five yeast-encoded macrocyclic peptide naïve libraries.

| **MATLAB script** | **script description** |
|---|---|
| >> *Step1 ('indelmut','on','bc',{'...','...','...','...'})* | Separation of sequence containing the correct barcode from others. The indelmut option allows the presence of only one modification (mutation, insertion or deletion) inside the barcode sequence. |
| >> *Step2 ('start', 'GAAGC', 'end', 'TCCGGT', 'uplimit', 19, 'downlimit', 11,'keepqf','on','translate','on')* | Translation of the peptide sequence included between the DNA regions 'GAAGC' and 'TCCGGT'. Peptide shorter than 11 amino acids and longer than 19 were discarded. The script was modified to translate the amber codon (TAG) as letter 'Z' and not 'N'. |
| >>*LoopLengthsZ* | This code allows the identification and counting of amber codons present within the polyclonal population. Adapted from LoopLenghts script [1]. |
| >>*LoopLengths* | This code allows the identification and counting of cysteine residues present within the polyclonal population. The sequences were further divided into different outputs based on the position and total number of cysteine residues. |
| >>*AminoCount* | Counting the occurrence of amino acids within the random peptide ring. Definition of amino acid enrichment or depletion versus theoretical probability of occurrence of the same amino acids when applying the degenerate NNK codon. |

**Supplementary table 4**. MATLAB scripts used for bioinformatic analysis of next generation sequencing (NGS) data of yeast-encoded macrocyclic peptide naïve libraries. The command lines (left) and their description (right) are reported for each step.

| | naïve macrocyclic peptide libraries | | | | |
|---|---|---|---|---|---|
| | $CX_7C$ | $CX_9C$ | $CX_3CX_9C$ | $CX_6CX_6C$ | $CX_9CX_3C$ |
| nº barcode DNA sequences | 1162740 | 764677 | 564635 | 655656 | 854067 |
| nº high quality peptide sequences * | 572952 | 188577 | 229524 | 281134 | 370332 |

**Supplementary table 5**. Statistics of next generation sequencing (NGS) analyses of yeast-encoded macrocyclic peptide naïve libraries. The number (nº) of barcode DNA sequences and identified high quality macrocyclic peptide sequences are reported for each naïve library. *: after the application of a quality filter to exclude readings with more than two bases with a quality score lower than Q18.

| naïve libraries | | frequency (%) additional or missing cysteine residues | | | |
|---|---|---|---|---|---|
| | | **-1 cys** | **+ 0 cys** | **+1 cys** | **+2 cys** |
| **1** | $CX_7C$ | 0.63% | 74.22% | 22.11% | 2.8% |
| **2** | $CX_9C$ | 1.29% | 68.69% | 25.48% | 4.09% |
| **3** | $CX_3CX_9C$ | 2.30% | 59.19% | 29.80% | 7.98% |
| **4** | $CX_6CX_6C$ | 1.85% | 56.62% | 31.49% | 9.30% |
| **5** | $CX_9CX_3C$ | 1.76% | 55.95% | 31.94% | 9.53% |

**Supplementary table 6**. Experimental probability of identifying additional or fewer cysteine residues in the yeast-encoded macrocyclic peptide naïve libraries. One (+ 1) or two (+ 2) additional cysteine residues occur at a rate of approximately 22 – 31% and 2 – 8%, respectively. Macrocycle peptides lacking one cysteine (- 1) are also present, thought at a low percentage (< 2%). Macrocyclic peptide sequences with fewer cysteine residues, as well as shorter or longer amino acid sequences, are often the result of inadequate nucleotide chemical coupling in the production of degenerated oligonucleotides used for the generation of the naive libraries.

| naïve libraries | | frequency (%) amber stop codon ('TAG') | | | |
|---|---|---|---|---|---|
| | | **0** | **1** | **2** | **3** |
| **1** | $CX_7C$ | 50.03% | 39.49% | 9.47% | 1.00% |
| **2** | $CX_9C$ | 50.06% | 37.14% | 11.23% | 1.55% |
| **3** | $CX_3CX_9C$ | 50.20% | 32.29% | 14.37% | 3.10% |
| **4** | $CX_6CX_6C$ | 50.18% | 32.90% | 14.03% | 2.85% |
| **5** | $CX_9CX_3C$ | 50.15% | 33.58% | 13.61% | 2.63% |

**Supplementary table 7**. Experimental probability of identifying additional amber stop codon ('TAG') in yeast-encoded macrocyclic peptide naïve libraries. The occurrence of 'TAG' codons is 1.5 times higher than the theoretical value expected for 'NNK'. One (1), two (2) and three (3) additional 'TAG' codons occur at a rate of approximately 32 – 39%, 9 – 14% and 1 – 3%, respectively. The presence of 'TAG' codons results in reduced effective library diversity as these sequences will not be displayed on the surface of yeast cells.

| amino acid | ratio experimental and theoretical frequency of single amino acid | | | | |
|---|---|---|---|---|---|
| | **$CX_7C$** | **$CX_9C$** | **$CX_3CX_9C$** | **$CX_6CX_6C$** | **$CX_9CX_3C$** |
| C | 1.30 | 1.27 | 1.28 | 1.40 | 1.42 |
| A | 0.95 | 0.93 | 0.88 | 0.89 | 0.92 |
| L | 0.88 | 0.87 | 1.07 | 1.06 | 1.01 |
| V | 1.29 | 1.25 | 1.42 | 1.44 | 1.45 |
| I | 0.97 | 1.02 | 0.94 | 0.99 | 0.98 |
| N | 0.95 | 1.03 | 0.74 | 0.78 | 0.79 |
| D | 1.12 | 1.09 | 0.91 | 0.97 | 1.01 |
| R | 1.02 | 1.03 | 0.98 | 0.94 | 0.96 |
| E | 1.06 | 1.07 | 0.98 | 0.92 | 0.94 |
| Q | 0.82 | 0.81 | 0.80 | 0.76 | 0.75 |
| Z | 1.06 | 1.05 | 1.16 | 1.11 | 1.06 |
| G | 1.34 | 1.37 | 1.27 | 1.23 | 1.29 |
| H | 0.87 | 0.83 | 0.72 | 0.79 | 0.81 |
| K | 0.10 | 1.05 | 0.91 | 0.85 | 0.84 |
| M | 1.08 | 1.08 | 1.19 | 1.14 | 1.10 |
| F | 0.99 | 1.02 | 1.17 | 1.22 | 1.15 |
| P | 0.60 | 0.59 | 0.57 | 0.58 | 0.59 |
| S | 0.92 | 0.92 | 0.92 | 0.94 | 0.93 |
| T | 0.83 | 0.83 | 0.74 | 0.75 | 0.75 |

| | | | | | |
|---|---|---|---|---|---|
| W | 1.20 | 1.19 | 1.40 | 1.31 | 1.29 |
| Y | 1.05 | 1.05 | 0.98 | 1.06 | 1.05 |

**Supplementary table 8**. Ratio between experimental (NGS data) and theoretical (codon 'NNK') frequency of single amino acid. The amino acids are indicated as one letter code. The amber stop codon (‘TAG’) is shown as ‘Z’. A ratio less than 1 indicates that the amino acid is underrepresented, while a ratio greater than 1 indicates that the amino acid is enriched. Some amino acids are overrepresented (C, cysteine; V, valine; W, tryptophan; G, glycine), while others are underrepresented (Q, glutamine; H, histidine; P, proline; T, threonine) with an experimental/theoretical frequency ratio of 1.2 – 1.5 and 0.8 – 0.5, respectively.

| protein target (PT) | | | |
|---|---|---|---|
| **ID code** | **protein name** | **UniProt code** | **source** |
| PT1 | aldolase | P00883 | GE Healthcare (cod. n. 28403842) |
| PT2 | streptavidin | P22629 | Thermo Fisher Scientific (cod. n. 84547) |
| PT3 | carbapenemase | Q09HD0 | Produced in house [2] |
| PT4 | carbonic anhydrase | P00921 | Fluka (cod. n. C2624) |
| PT5 | α-chymotrypsin | P00766 | Fluka (cod. n. C4129) |

**Supplementary table 9**. Protein targets (PTs) used during the combinatorial high-throughput screening process. ID code, name, UniProt code and source are reported for each protein target tested.

| protein target (PT) | | | | | |
|---|---|---|---|---|---|
| **ID code** | **M.W. (kDa)** | **ε ($M^{-1}$ $cm^{-1}$)** | **pI** | **state** | **PDB code** |
| PT1 | 157 | 34880 | 8.2 | tetrameric | 6ALD |
| PT2 | 52 | 41940 | 6.1 | tetrameric | 7EK8 |
| PT3 | 31 | 24075 | 6.2 | monomeric | 6TS9 |
| PT4 | 29 | 50420 | 6.4 | monomeric | 1VE9 |
| PT5 | 25 | 50585 | 8.3 | monomeric | 6DI8 |

**Supplementary table 10**. Biochemical properties of proteins targets (PTs) used during the combinatorial high-throughput screening process. ID code, molecular weight (M.W., kDa), pI, extinction coefficient (ε, $M^{-1}$ $cm^{-1}$), state in solution and PDB code are reported for each PT tested.

| ID code | barcode | DNA sequence (5' – 3') |
|---|---|---|
| NGS-F1 | n.i. | GGAAGAAGGTGTTCAATTGGACAAGAG AGAAGC |
| NGS-R1 | n.i. | CCAGGCAGTCCGAGAGGGTGAGGATGT TTGAGCGTAATCTGG |
| NGS-F2-A | CTGCACATCGAT | TCGTCGGCAGCGTCAGATGTGTATAAG AGACAGCTGCACATCGATGGAAGAAGG TGTTCAATTGGACAAGAG |
| NGS-F2-B | ATGTCCAACTAC | TCGTCGGCAGCGTCAGATGTGTATAAG AGACAGATGTCCAACTACGGAAGAAGG TGTTCAATTGGACAAGAG |
| NGS-F2-C | GCGTACACTCAC | TCGTCGGCAGCGTCAGATGTGTATAAG AGACAGGCGTACACTCACGGAAGAAGG TGTTCAATTGGACAAGAG |
| NGS-F2-D | TCTGAGGATCCA | TCGTCGGCAGCGTCAGATGTGTATAAG AGACAGTCTGAGGATCCAGGAAGAAGG TGTTCAATTGGACAAGAG |
| NGS-F2-E | CCTTACGATGTG | TCGTCGGCAGCGTCAGATGTGTATAAG AGACAGCCTTACGATGTGGGAAGAAGG TGTTCAATTGGACAAGAG |
| NGS-F2-F | AGTCAAGATGTT | TCGTCGGCAGCGTCAGATGTGTATAAG AGACAGAGTCAAGATGTTGGAAGAAGG TGTTCAATTGGACAAGAG |
| NGS-F2-G | GACACTGATGGT | TCGTCGGCAGCGTCAGATGTGTATAAG AGACAGGACACTGATGGTGGAAGAAG GTGTTCAATTGGACAAGAG |

| | | |
|---|---|---|
| NGS-F2-H | CACAGAGATACT | TCGTCGGCAGCGTCAGATGTGTATAAG AGACAGCACAGAGATACTGGAAGAAG GTGTTCAATTGGACAAGAG |
| NGS-F2-I | TGCTTGGATTCG | TCGTCGGCAGCGTCAGATGTGTATAAG AGACAGTGCTTGGATTCGGGAAGAAGG TGTTCAATTGGACAAGAG |
| NGS-F2-L | AGTGTACACTGC | TCGTCGGCAGCGTCAGATGTGTATAAG AGACAGAGTGTACACTGCGGAAGAAGG TGTTCAATTGGACAAGAG |
| NGS-R2 | n.i. | GTCTCGTGGGCTCGGAGATGTGTATAA GAGACAGCCAGGCAGTCCGAGAGGG |

**Supplementary table 11.** Oligonucleotides used for the NGS analysis of clones selected during the combinatorial high-throughput screening process against five different PTs. The ID code, barcode and DNA sequences (5' – 3') of each oligonucleotide is reported. Oligonucleotides NGS-F1 and NGS-R1 are universal and have been used in the first PCR reaction. Forward oligonucleotides NGS-F2-A, NGS-F2-B, NGS-F2-C, NGS-F2-D, NGS-F2-E, NGS-F2-F, NGS-F2-G, NGS-F2-H, NGS-F2-I and NGS-F2-L have been instead used in the second PCR reaction in combination with the universal NGS-R2 reverse oligonucleotide. Legend: n.i. = barcode not included.

| MATLAB script | script description |
| --- | --- |
| *>> Step1 ('indelmut','on','bc',{'...','...','...','...'})* | Separation of sequence containing the correct barcode from others. The indelmut option allows the presence of only one modification (mutation, insertion or deletion) inside the barcode sequence. |
| *>> Step2 ('start', 'GAAGC', 'end', 'TCCGGT', 'uplimit', 19, 'downlimit', 11,'keepqf','on','translate','on')* | Translation of the peptide sequence included between the DNA regions 'GAAGC' and 'TCCGGT'. Peptides shorter than 11 amino acids and longer than 19 were discarded. The script was modified to translate the amber codon (TAG) as letter 'Z' and not 'N'. |
| *>>CommonSeq ('cut off', 10)* | This code allows identification of unspecific peptide sequences among different protein targets. |
| *>> Clustering75('number_dif', 1000,'min_clustersize', 2)* | This code allows clustering of the 1000 most abundant sequences in families with at least 2 components and a sequence homology equal or higher than 75%. The script was modified to raise the homology from 50% to 75%. |
| *>>LoopLengths* | This code allows the identification and counting of cysteine residues present within the polyclonal population. The sequences were further divided into different outputs based on the position and total number of cysteine residues. |
| *>>AminoCount* | Counting the occurrence of amino acids within the random peptide ring. Definition of amino acid enrichment or depletion versus probability of occurrence of the same amino acids in the naïve libraries. |

**Supplementary table 12**. MATLAB scripts used for bioinformatic analysis of NGS data of clones selected during the combinatorial high-throughput screening process against five different PTs. The command lines (left) and their description (right) are reported for each step.

| | NGS analysis of selected clones | | | | |
|---|---|---|---|---|---|
| | **PT1** | **PT2** | **PT3** | **PT4** | **PT5** |
| nº unique DNA sequences* | 182300 | 28425 | 39947 | 451266 | 43139 |
| nº unique amino acid sequences* | 2810 | 840 | 757 | 87 | 895 |

**Supplementary table 13**. NGS analyses and statistics of clones selected during the combinatorial high-throughput screening process against five different PTs. *: after the application of a quality filter to exclude readings with more than two bases with a quality score lower than Q18.

| amino acid | ratio experimental and theoretical frequency of single amino acid | | | | |
|---|---|---|---|---|---|
| | **PT1** | **PT2** | **PT3** | **PT4** | **PT5** |
| C | 0.23 | 2.68 | 3.67 | 1.92 | 2.03 |
| A | 0.14 | 0.94 | 1.23 | 0.31 | 0.59 |
| L | 1.20 | 0.05 | 0.05 | 1.35 | 0.70 |
| V | 1.28 | 0.40 | 1.68 | 1.25 | 0.69 |
| I | 0.19 | 0.95 | 2.42 | 0.13 | 1.76 |
| N | 0.12 | 0.01 | 2.62 | 0.14 | 0.27 |
| D | 3.24 | 1.83 | 0 | 2.90 | 2.17 |
| R | 0.05 | 0.11 | 0.11 | 0 | 1.44 |
| E | 3.36 | 0.10 | 0.02 | 0 | 0.45 |
| Q | 0.03 | 3.28 | 0 | 0 | 1.54 |
| Z | 0 | 0 | 0 | 0 | 0 |
| G | 1.19 | 0.67 | 0.07 | 0.87 | 0.41 |
| H | 0.22 | 4.40 | 0.19 | 0.70 | 0.88 |
| K | 0.04 | 1.13 | 0.21 | 0 | 0.16 |
| M | 0.04 | 1.68 | 0.07 | 2.02 | 0.91 |
| F | 0.21 | 2.39 | 0.31 | 0.11 | 0.91 |
| P | 0.21 | 2.22 | 1.92 | 4.02 | 0.61 |
| S | 1.10 | 0.62 | 0.06 | 0 | 0.55 |
| T | 2.09 | 0.01 | 1.49 | 0.75 | 1.57 |

| | | | | | |
|---|---|---|---|---|---|
| W | 2.74 | 1.46 | 3.68 | 2.01 | 3.64 |
| Y | 3.42 | 3.36 | 4.65 | 4.38 | 0.77 |

**Supplementary table 14**. Ratio experimental (NGS data) and theoretical (naïve libraries) frequency of single amino acid in clones selected during the combinatorial high-throughput screening process against five different PTs. The amino acids are indicated as one letter code. The amber stop codon (‘TAG’) is shown as ‘Z’. A ratio lower than 1 indicates that the amino acid is underrepresented, while a ratio greater than 1 indicates that the amino acid has been enriched during the selection process.

| ID code | amino acid sequence | DNA sequence (5' – 3') |
|---|---|---|
| **MP2.3** | ACIMVHYDRVECSG | TCAATTGGACAAGAGAGAAGCTTGT ATTATGGTGCATTATGATCGGGTTGA GTGTTCCGGTGGTGGTGGCTCTGGTG G |
| **MP3.3** | ACYVTCFGWYCYRMSCSG | TCAATTGGACAAGAGAGAAGCTTGT TACGTTACTTGTTTCGGTTGGTACTG TTACAGAATGTCTTGTTCTGGTTCCG GTGGTGGTGGCTCTGGTGG |
| **MP4.2** | ACHVGPYYMLACSG | TCAATTGGACAAGAGAGAAGCTTGT CATGTGGGTCCTTATTATATGCTGGC TTGTTCCGGTGGTGGTGGCTCTGGTG G |
| **MP4.3** | ACYGPYYVLCSG | TCAATTGGACAAGAGAGAAGCTTGT TATGGTCCTTATTATGTTTTGTGTTCC GGTGGTGGTGGCTCTGGTGG |
| **MP5.3** | ACCYRIYCPCSG | GGAAGAAGGTGTTCAATTGGACAAG AGAGAAGCTTGTTGTTATAGGATTTA TTGTCCGTGTTCCGGTGGTGGTGGCT CTGGTGG |
| **MP5.5** | ACLHVCLERGQCLFMCSG | GGAAGAAGGTGTTCAATTGGACAAG AGAGAAGCTTGTCTGCATGTGTGTTT GGAGCGTGGGCAGTGTCTGTTTATGT GTTCCGGTGGTGGTGGCTCTGGTGG |
| **R** | | GCCAGATGTTGTCGAACTTTCTGATT TAGTGG |

**Supplementary table 15**. Oligonucleotides used for PCR amplification and cloning of additional macrocyclic peptide (MP) sequences selected during the combinatorial high-throughput screening process against five different PTs identified by next generation sequencing analysis. The ID code, amino acidic (N- to C-terminus) and DNA (5' – 3') sequences of each oligonucleotide is reported. MP-R reverse oligonucleotide is universal and has been used in all PCR reactions in combination with sequence-specific forward oligonucleotides.

| binding affinity of macrocyclic peptides using yeast surface titration | | |
|---|---|---|
| **ID code** | **amino acid sequence** | **$K_D^{app}$ ± S.E. (nM)** |
| **MP1.1** | ACEVLSYDWTGCSG | 0.6 ± 0.1 |
| **MP2.1** | ACSMCHPQGDFCYWACSG | 2.1 ± 0.3 |
| **MP2.2** | ACHYHPQFYVICSG | 0.2 ± 0.1 |
| **MP2.3** | ACIMVHYDRVECSG | 1.9 ± 0.2 |
| **MP3.1** | ACYIYCANPWVCWVTCSG | >1000 |
| **MP3.2** | ACRYICLSFRHCGWFCSG | 2.2 ± 0.5 |
| **MP3.3** | ACYVTCFGWYCYRMSCSG | >1000 |
| **MP4.1** | ACLVGPYYMLDCSG | 41.0 ± 4.9 |
| **MP4.2** | ACHVGPYYMLACSG | 172.0 ± 10.5 |
| **MP4.3** | ACYGPYYVLCSG | 369.0 ± 2.3 |
| **MP4.4** | ACCPTCPDWWVCSG | 680 ± 4.7 |
| **MP5.1** | ACRDWTWIQCSG | 1.9 ± 0.6 |
| **MP5.2** | ACHRWGVRFSLCSG | 3.7± 0.5 |
| **MP5.3** | ACCYRIYCPCSG | 2.4 ± 0.7 |
| **MP5.4** | ACAWCLRGTICCSG | 0.2 ± 0.01 |
| **MP5.5** | ACLHVCLERGQCLFMCSG | 9.2 ± 1.2 |
| **MP5.6** | ACVLMLRRCRILGWCCSG | 1.5 ± 0.25 |

**Supplementary table 16**. Binding affinities of the most abundant selected macrocyclic peptide ligands displayed on the surface of yeast. Apparent equilibrium binding constant ($K_D^{app}$) of each yeast-encoded macrocyclic peptide (MP) with its respective protein target (PT) were determined by flow cytometry at 25 °C and physiological pH 7.4. The indicated values are means of at least three independent experiments. S.E., standard error.

| | binding affinity of macrocyclic peptides using surface plasmon resonance | | |
|---|---|---|---|
| **ID code** | **$k_{on}$ ± S.E. ($M^{-1}\cdot s^{-1}$)** | **$k_{off}$ ± S.E. ($s^{-1}$)** | **$K_D$ ± S.E. (M)** |
| **MP1.1** | $2.6 \times 10^{4}$ | $1.5 \times 10^{-2}$ | $576 \times 10^{-9}$ |
| **MP2.2** | $6.4 \times 10^{1}$ | $1.1 \times 10^{-8}$ | $0.2 \times 10^{-9}$ |
| **MP3.2.1** | $1.3\times 10^{5}$ | $4.1 \times 10^{-4}$ | $3.2 \times 10^{-9}$ |
| **MP3.2.2** | n.b. | n.b. | n.b. |
| **MP3.2.3** | n.b. | n.b. | n.b. |
| **MP4.2** | $1.8 \times 10^{3}$ | $3.0 \times 10^{-4}$ | $171 \times 10^{-9}$ |
| **MP5.4.3** | $4.9 \times 10^{7}$ | $4.3 \times 10^{-3}$ | $0.08 \times 10^{-9}$ |

**Supplementary table 17**. Binding kinetics of seven chemically synthesised macrocyclic peptide ligands toward their respective immobilised PTs were determined by surface plasmon resonance at 25 °C and physiological pH (7.4). S.E., standard error; n.b., not binding.

| inhibitory activity of chemically synthesised macrocyclic peptides | |
|---|---|
| **ID code** | **$K_i$ ± S.E. (nM)** |
| **MP5.4.1** | 51.0 ± 1.2 |
| **MP5.4.2** | 11.0 ± 0.7 |
| **MP5.4.3** | 0.36 ± 0.2 |

**Supplementary table 18**. The inhibitory activity ($K_i$) values of three synthetic macrocyclic peptides (MP5.4.1, MP5.4.2 and MP5.4.3) towards bovine α-chymotrypsin (PT5) protease were determined at 25 °C and physiological pH 7.4 using the chromogenic N-Succinyl-Ala-Ala-Pro-Phe *p*-nitroanilide substrate at a concentration of 100 µM. The $K_m$ value (114 µM) of bovine α-chymotrypsin protease was determined by standard Michaelis-Menten kinetics and used in the calculation of the reported $K_i$ values. The shown values are the means of three independent experiments. S.E., standard error.

| binding affinity of macrocyclic peptides using yeast surface titration – $K_D^{app}$ ± S.E. (nM) | | | |
|---|---|---|---|
| | **PT2** | **PH2.1** | **PH2.2** |
| **MP2.1** | 2.1 ± 0.2 | 0.2 ± 0.1 | n.b. |
| **MP2.2** | 0.2 ± 0.1 | 0.8 ± 0.2 | n.b. |
| **MP2.3** | 1.9 ± 0.2 | 0.8 ± 0.2 | n.b. |

**Supplementary table 19**. Binding affinities of macrocyclic peptide MP2.1, MP2.2 and MP2.3 displayed on the surface of yeast against streptavidin (PT2) and two homologue proteins, namely strep-tactin (PH2.1) and neutravidin (PH2.2). Apparent equilibrium binding constant ($K_D^{app}$) of each yeast-encoded MP with protein targets PT2, PH2.1 and PH2.2 were determined by flow cytometry at 25 °C and physiological pH 7.4. The indicated values are means of at least three independent experiments. S.E., standard error. n.b., not binding.

| binding affinity of macrocyclic peptides using yeast surface titration – $K_D^{app}$ ± S.E. (nM) | | | |
|---|---|---|---|
| | **PT3** | **PH3.1** | **PH3.2** |
| **MP3.2** | 2.2 ± 0.5 | n.b. | n.b. |

**Supplementary table 20**. Binding affinities of macrocyclic peptide MP3.2 displayed on the surface of yeast against carbapenemase GES-5 from *Klebsiella pneumoniae* (PT3) and two homologue proteins, namely carbapenemase KPC-2 from *Klebsiella oxytoca* (PH3.1) and the extended-spectrum β-lactamase CTX-M-15 from *Escherichia coli* (PH3.2). Apparent equilibrium binding constant ($K_D^{app}$) of yeast-encoded macrocyclic peptide MP3.2 with protein targets PT3, PH3.1 and PH3.2 were determined by flow cytometry at 25 °C and physiological pH 7.4. The indicated values are means of at least three independent experiments. S.E., standard error. n.b., not binding.

| inhibitory activity of chemically synthesised macrocyclic peptides – $K_i$ ± S.E. (nM) | | | | |
|---|---|---|---|---|
| | **PT5** | **PH5.1** | **PH5.2** | **PH5.3** |
| **MP5.4.3** | 0.36 ± 0.2 | n.i. | n.i. | n.i. |

**Supplementary table 21**. Inhibitory activity ($K_i$) values of synthetic 'two rings' macrocyclic peptide MP5.4.3 towards bovine a-chymotrypsin (PT5) and three homologue serine proteases, namely trypsin (PH5.1), plasmin (PH5.2) and thrombin (PH5.3) were determined at 25 °C and physiological pH 7.4. The chromogenic substrate N-Succinyl-Ala-Ala-Pro-Phe *p*-nitroanilide at final concentration of 100 µM was used for bovine a-chymotrypsin (PT5). The fluorogenic substrate Z-Gly-Gly-Arg-AMC at final concentration of 50 µM was used for human trypsin (PH5.1) and human thrombin (PH5.3). The fluorogenic substrate H-D-Val-Leu-Lys-AMC at final concentration of 50 µM was used for human plasmin (PH5.2). The $K_m$ value of bovine a-chymotrypsin (PT5; $K_m$ = 114 µM), human trypsin (PH5.1; $K_m$ = 54 µM), human plasmin (PH5.2; $K_m$ = 610 µM) and human thrombin (PH5.3; $K_m$ = 199 µM) were determined by standard Michaelis-Menten kinetics and used in the calculation of the reported $K_i$ values [3]. The shown values are the means of three independent experiments. S.E., standard error. n.i., not inhibition.

| PDB ID | 9F6H |
|---|---|
| **Data collection** | |
| Diffraction source | ID30-A (ESRF) |
| Wavelength (Å) | 0.9677 |
| Temperature (K) | 100 |
| Detector | Dectris Eiger X 4M |
| Crystal-detector distance (mm) | 100 |
| Rotation range per image (°) | 0.05 |
| Exposure time per image (s) | 0.013 |
| Space group | $P6_1$ |
| No. of molecules/ASU | 1 |
| *a, b, c* (Å) | 103.47, 103.47, 46.42 |
| *α, β, γ* (°) | 90, 90, 120 |
| Total no. of reflections | 65600 (5533) |
| No. of unique reflections | 10807 (1132) |
| Completeness (%) | 98.0 (98.6) |
| Redundancy | 6.1 (4.9) |
| CC1/2 | 0.997 (0.439) |
| $\langle I/\sigma(I) \rangle$ | 10.5 (1.6) |
| $R_{mrg}$ | 0.137 (0.986) |
| **Refinement statistics** | |

| | |
|---|---|
| Resolution range (Å) | 44.80 – 2.42 |
| No. of reflections, working set | 10807 |
| No. of reflections, test set | 488 |
| Final $R_{cryst}$ | 0.213 |
| Final $R_{free}$ | 0.280 |
| No. of non-H atoms | |
| protein | 1778 (chains A, B and C) |
| water | 41 |
| others | 97 (macrocyclic peptide, chain L) |
| total | 1916 |
| R.m.s. deviations | |
| bonds (Å) | 0.0081 |
| angles (°) | 1.426 |
| Average $B$ factors (Å$^2$) | 34.0 |
| Ramachandran plot | |
| Most favoured (%) | 96 |
| Allowed (%) | 4 |

**Supplementary table 22**. X-ray data collection and refinement statistics of bovine α-chymotrypsin (PT5) in complex with the 'two rings' macrocycle peptide MP5.4.3. A single crystal was used to collect all diffraction data. Highest-resolution shell statistics are shown within brackets.

| | **PT5** | **MP5.4.3** |
|---|---|---|
| nº of residues at the interface | 19 | 9 |
| nº of salt bridges | 1 | 1 |
| nº of hydrogen bonds | 10 | 10 |
| nº of hydrogen bonds water mediated | 1 | 1 |
| nº of no-bonded contacts | 51 | 51 |
| buried surface area (Å$^2$) | 784 | 911 |

**Supplementary table 23.** Inter-molecular interactions between bovine α-chymotrypsin (PT5) and the 'two rings' macrocyclic peptide MP5.4.3. Number of residues of PT5 and MP5.4.3 at the interface, total number of inter-molecular salt bridges, hydrogen bonds (direct or water mediated) and non-polar interactions have been defined using the software LIGPLOT+ [4]. Buried surface areas (Å$^2$) were calculated using the software PDBsum [5] with a probe of 1.4 Å radius and are reported here for both PT5 and MP5.4.3. The designation 'buried' implies that the residues are at least partially inaccessible to bulk solvent because of the proximity of the interface surfaces of both protein target and macrocyclic peptide.

| PT5 | MP5.4.3 | distance interaction |
|---|---|---|
| O / Ser217 | NE1 / Trp4 | 3.1 Å [HB] |
| O / Gly216 | N / Cys2 | 3.0 Å [HB] |
| N / Gly216 | O / Cys2 | 3.1 Å [HB] |
| N / Gly193 | O / Trp4 | 3.0 Å [HB] |
| O / Phe41 | N / Leu6 | 4.1 Å [PI] |
| O / Cys58 | NH1 / Arg7 | 3.8 Å [HB] |
| N / Asp194 | O / Trp4 | 3.8 Å [HB] |
| N / Ser195 | O / Trp4 | 3.4 Å [HB] |
| OG / Ser195 | N / Cys5 | 3.2 Å [HB] |
| O / Ser214 | N / Trp4 | 3.5 Å [HB] |
| N / Ser218 | N / Ala1 | 2.9 – 3.5 Å [HB]* |
| OD1 / Asp64 | NH2 / Arg7 | 3.8 Å [HB] |

**Supplementary table 24.** Unique polar inter-molecular interactions between bovine α-chymotrypsin (PT5) and the 'two rings' macrocycle peptide MP5.4.3. Atoms (left) and residues (right) of PT5 forming polar inter-molecular interactions with MP5.4.3 are reported. Optimal hydrogen bonds [HB], polar interactions [PI] and distances (Å) have been defined using PROFUNC and LIGPLOT+ [4,6]. *: water mediated contacts: distance atom/residue MP5.4.3 – $H_2O$ (left); atom/residue (PT5) – $H_2O$ (right).

| PT5 | MP5.4.3 | distance interaction |
|---|---|---|
| CB / Ser218 | SG / Cys2 | 3.3 Å |
| CA / Ser218 | SG / Cys2 | 3.6 Å |
| O / Ser217 | CZ2 / Trp4 | 3.9 Å |
| O / Ser217 | CE2 / Trp4 | 3.8 Å |
| N / Ser217 | CZ2 / Trp4 | 3.4 Å |
| C / Gly216 | C / Ala1 | 3.7 Å |
| C / Gly216 | CA / Ala1 | 3.3 Å |
| C / Gly216 | CZ2 / Trp4 | 3.6 Å |
| C / Gly216 | CE2 / Trp4 | 3.9 Å |
| CA / Gly216 | CZ2 / Trp4 | 3.3 Å |
| CA / Gly216 | CH2 / Trp4 | 3.8 Å |
| CA / Gly216 | CE2 / Trp4 | 3.7 Å |
| N / Gly216 | CZ2 / Trp4 | 3.8 Å |
| N / Gly216 | CE2 / Trp4 | 3.6 Å |
| CB / Trp215 | CA / Ala3 | 3.8 Å |
| CB / Trp215 | O / Cys2 | 3.2 Å |
| CB / Trp215 | C / Cys2 | 3.8 Å |
| CA / Trp215 | O / Cys2 | 3.6 Å |
| O / Ser214 | CA / Ala3 | 3.8 Å |
| OG / Ser195 | CA / Cys5 | 3.7 Å |

| | | |
|---|---|---|
| OG / Ser195 | C / Trp4 | 2.8 Å |
| OG / Ser195 | CB / Trp4 | 3.7 Å |
| OG / Ser195 | CA / Trp4 | 3.4 Å |
| CB / Ser195 | O / Trp4 | 3.2 Å |
| CB / Ser195 | C / Trp4 | 3.6 Å |
| CA / Ser195 | O / Trp4 | 3.8 Å |
| C / Met192 | O / Trp4 | 3.7 Å |
| SD / Met192 | CB / Cys12 | 3.8 Å |
| SD / Met192 | CD1 / Trp4 | 3.6 Å |
| CG/Met192 | O / Cys5 | 3.4 Å |
| CB / Met192 | CD/Leu6 | 3.8 Å |
| CB / Met192 | O / Cys5 | 3.5 Å |
| CA / Met192 | O / Trp4 | 3.3 Å |
| CA / Met192 | C / Trp4 | 3.8 Å |
| O / Ser190 | CZ2 / Trp4 | 3.6 Å |
| O / Ser190 | CH2 / Trp4 | 3.5 Å |
| O / Ser190 | CZ3 / Trp4 | 3.7 Å |
| O / Ser190 | CE3 / Trp4 | 3.6 Å |
| OG / Ser189 | CH2 / Trp4 | 3.7 Å |
| CZ3 / Trp172 | CB / Ala1 | 3.8 Å |
| CG1 / Ile99 | CD1 / Ile10 | 3.8 Å |

| | | |
|---|---|---|
| OH / Tyr94 | CD1 / Ile10 | 3.7 Å |
| CD2 / His57 | SG / Cys5 | 3.5 Å |
| CD2 / His57 | CB / Ala3 | 3.8 Å |
| NE2 / His57 | CB / Cys5 | 3.8 Å |
| NE2 / His57 | CB / Ala3 | 3.5 Å |
| CE2 / Phe41 | NH1 / Arg7 | 3.5 Å |
| CD2 / Phe41 | NH1 / Arg7 | 3.6 Å |

**Supplementary table 25.** Unique non-polar inter-molecular interactions between bovine α-chymotrypsin (PT5) and the 'two rings' macrocycle peptide MP5.4.3 with a distance below 4.0 Å. Optimal interactions and distances (Å) have been defined using PROFUNC and LIGPLOT+ [4,6].

| naïve library | | | DNA sequence (5' – 3') |
|---|---|---|---|
| **1** | $CX_7C$ | F1 | GGAAGAAGGTGTTCAATTGGACAAGAGAGAAGCTTG<br>TNNKNNKNNKNNKNNKNNKNNKTGTTCCGGTGGTGG<br>TGGCTCTGGTGG |
| **2** | $CX_9C$ | F2 | GGAAGAAGGTGTTCAATTGGACAAGAGAGAAGCTTG<br>TNNKNNKNNKNNKNNKNNKNNKNNKNNKTGTTCCGG<br>TGGTGGTGGCTCTGGTGG |
| **3** | $CX_3CX_9C$ | F3 | GGAAGAAGGTGTTCAATTGGACAAGAGAGAAGCTTG<br>TNNKNNKNNKTGTNNKNNKNNKNNKNNKNNKNNKN<br>NKNNKTGTTCCGGTGGTGGTGGCTCTGGTGG |
| **4** | $CX_6CX_6C$ | F4 | GGAAGAAGGTGTTCAATTGGACAAGAGAGAAGCTTG<br>TNNKNNKNNKNNKNNKNNKTGTNNKNNKNNKNNKN<br>NKNNKTGTTCCGGTGGTGGTGGCTCTGGTGG |
| **5** | $CX_9CX_3C$ | F5 | GGAAGAAGGTGTTCAATTGGACAAGAGAGAAGCTTG<br>TNNKNNKNNKNNKNNKNNKNNKNNKNNKTGTNNKN<br>NKNNKTGTTCCGGTGGTGGTGGCTCTGGTGG |
| | | R | GCCAGATGTTGTCGAACTTTCTGATTTAGTGG |

**Supplementary table 26**. Synthetic oligonucleotides (5' to 3') used for the generation of yeast-encoded naïve macrocycle peptide libraries.

| flow cytometry reagents | applied dilution | |
|---|---|---|
| | FACS sorting | flow cytometry analysis |
| mouse anti-HA IgG1 (1 mg $mL^{-1}$) | 1:1000 | 1:1000 |
| goat anti-mouse IgG-DyLight 488 (1 mg $mL^{-1}$) | 1:200 | 1:500 |
| neutravidin-DyLight 650 (1 mg $mL^{-1}$) | 1:200 | 1:500 |
| streptavidin-DyLight 650 (1 mg $mL^{-1}$) | 1:200 | 1:500 |

**Supplementary table 27**. Commercial recombinant proteins and antibodies used for yeast surface display selection (FACS sorting) and equilibrium binding titration (flow cytometry analysis). Mouse anti-HA IgG1 (clone 2-2.2.14; 1 mg $mL^{-1}$) has been used as primary reagent for the display detection. Goat anti-mouse IgG-Alexa Fluor 488 (2 mg $mL^{-1}$), neutravidin-DyLight 650 (1 mg $mL^{-1}$) and streptavidin-DyLight 650 (1 mg $mL^{-1}$) have been used as secondary reagents. All reagents were purchased from Thermo Fisher Scientific.

### Supplementary figures

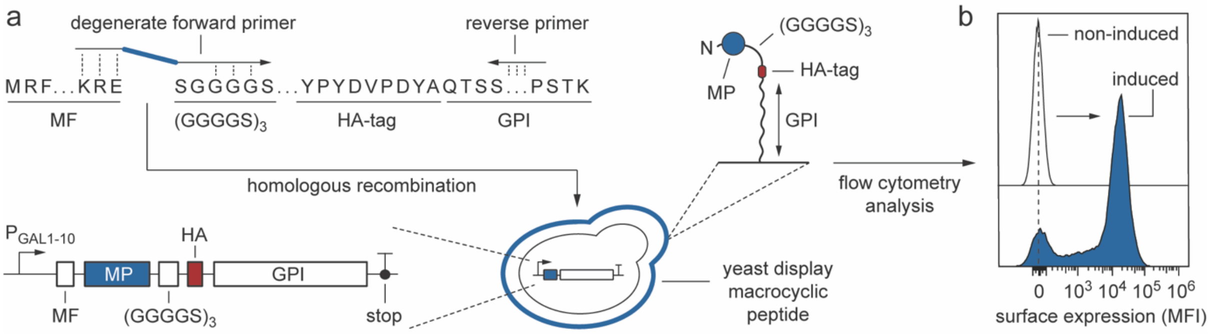


**Supplementary figure 1**. Generation of yeast-encoded macrocyclic peptide libraries. **a**) The insert is created by appending the DNA encoding the randomised macrocyclic peptide (MP) sequence (blue) to the N-terminus of the $(G_4S)_3$ – HA – GPI encoding gene in a PCR reaction using a forward degenerate primer ‘F’ and a universal reverse primer ‘R’. The amplified insert and the double digested pCT-GPI backbone are further assembled via homologous recombination in yeast cells. The final construct contains the following elements: a GAL1-10 promoter, an α-mating factor (MF) secretion signal, the randomised macrocyclic peptide sequence (blue), a long and flexible spacer $(G_4S)_3$, the hemagglutinin tag (HA; YPYDVPDYA, red) and the cysteine-free glycosylphosphatidylinositol (GPI) cell-surface anchor protein. The level of expression of the MP on the surface of yeast cells is assessed by flow cytometry; **b**) Histograms representing the flow-cytometry analysis of expression of macrocyclic peptide displayed on the surface of induced (blue) and not induced (white) yeast cells after staining with mouse anti-HA antibody (primary) and goat anti-mouse IgG-DyLight 488 antibody (secondary).

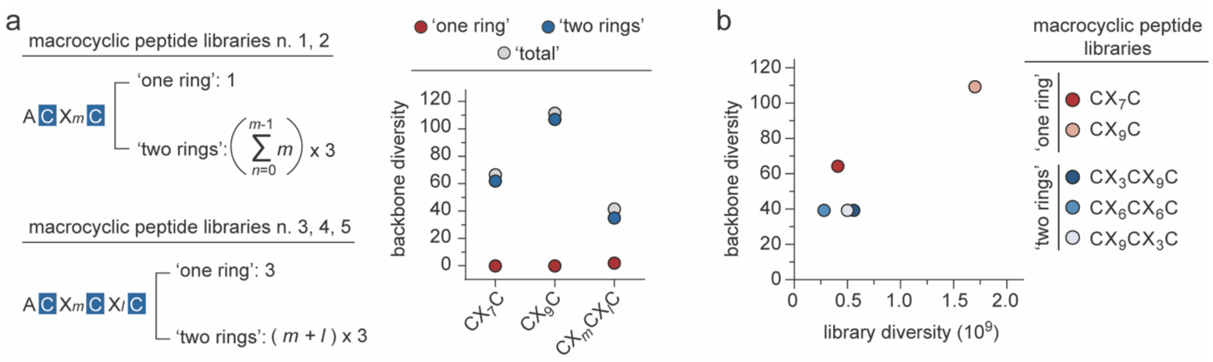


**Supplementary figure 2**. Size and backbone diversity of ‘one ring’ and ‘two rings’ macrocyclic peptide libraries. The backbone diversity is the sum of all possible unique combinations of ‘one-ring’ (two cysteines, one disulfide bridge) or ‘two rings’ (four cysteines, two disulfide bridges) macrocyclic peptide ligands that can rise from the CX$_7$C, CX$_9$C and CX$_m$CX$_l$C libraries (where $m$ = 3, 6 or 9 and $l$ = 9, 6 or 3). The ‘one ring’ macrocycle peptide sequences can form only one isomer, while ‘two rings’ macrocycle peptide sequences can yield three different isomers. Theoretically, the CX$_7$C and CX$_9$C libraries have a higher probability to form ‘one ring’ macrocyclic peptide ligands (>95%), while CX$_m$CX$_l$C libraries have higher probability to yield ‘two rings’ macrocyclic peptide ligands (>30%). Experimental next generation sequencing (NGS) analyses of the naïve libraries confirmed the presence of all possible theoretically predicted unique combinations of ‘one ring’ and ‘two rings’ macrocyclic peptides, albeit with lower percentages than those predicted. **a**) ‘one ring’ (red), ‘wo ring’ (blue) and ‘total’ (grey) backbone diversity of library CX$_7$C, CX$_9$C and CX$_m$CX$_l$C (wherein $m$ = 3, 6 or 9 and $l$ = 9, 6 or 3) are reported; **b**) Library diversity and ‘total’ backbone diversity of ‘one ring’ and ‘two rings’ macrocyclic peptide libraries CX$_7$C (red), CX$_9$C (light red), CX$_3$CX$_9$C (dark blue), CX$_6$CX$_6$C (blue) and CX$_9$CX$_3$C (light blue) are reported in the $x$-axis and $y$-axis, respectively.

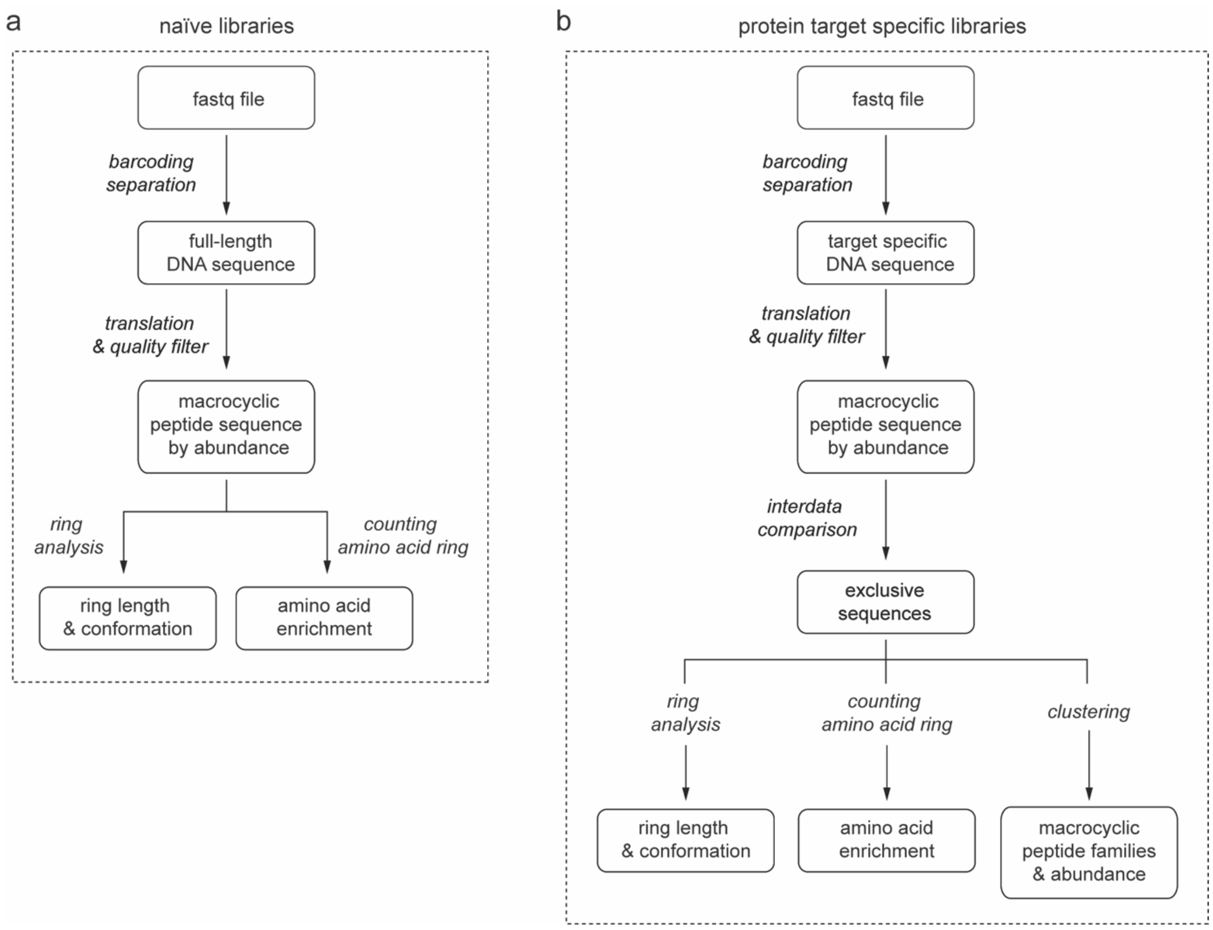


**Supplementary figure 3**. Schematic representation of the pipelines applied to analyse the next generation sequencing (NGS) data derived from either the naïve libraries (**a**) or the target specific selected clones (**b**). Analysis was performed on FastQ files input. The reads were initially separated into several files according to their barcode for the identification of target specific DNA sequences. Following the removal of low-quality sequences from the dataset, the remaining sequences were translated and sorted by abundance. The peptide sequence datasets were compared to each other for the identification of target exclusive motifs. Starting from these datasets, three simultaneous analyses were performed: distribution of the isolated macrocyclic peptide sequences based on loop length and topologies (*loop analyses*); definition of enrichment or depletion of amino acids distribution inside the rings (*counting amino acids loop*) and the most abundant macrocyclic peptides are compared and clustered in consensus groups or sub-families of consensus groups (*clustering*).

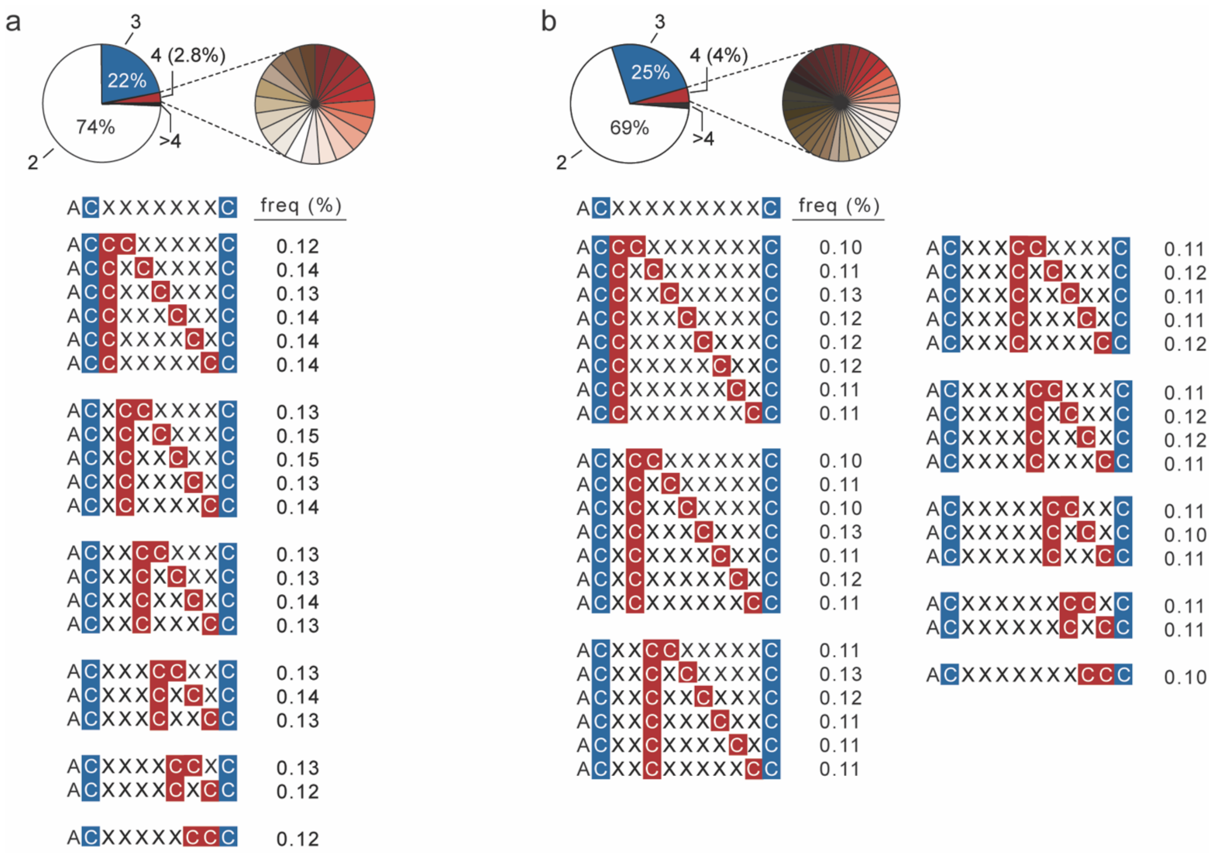


**Supplementary figure 4**. The distribution and frequency of appearance of the third and fourth cysteine residue in the naïve 'one ring' macrocyclic peptide libraries $CX_7C$ (**a**) and $CX_9C$ (**b**) obtained by NGS analyses. The position of N-terminal alanine (A, black), fixed cysteines (C, blue), random fourth cysteines (C, red) and random amino acids (X, black) are shown.

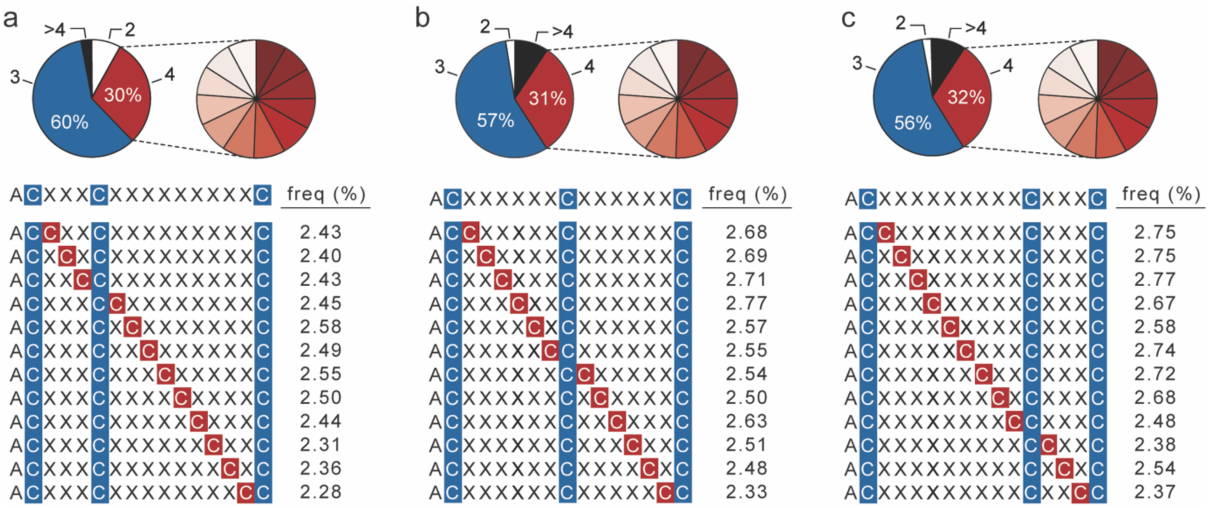


**Supplementary figure 5**. The distribution and frequency of appearance of the fourth cysteine residue in the naïve 'two rings' macrocyclic peptide libraries $CX_3CX_9C$ (**a**), $CX_6CX_6C$ (**b**) and $CX_9CX_3C$ (**c**) obtained by NGS analyses. The position of N-terminal alanine (A, black), fixed cysteines (C, blue), random fourth cysteines (C, red) and random amino acids (X, black) are shown.

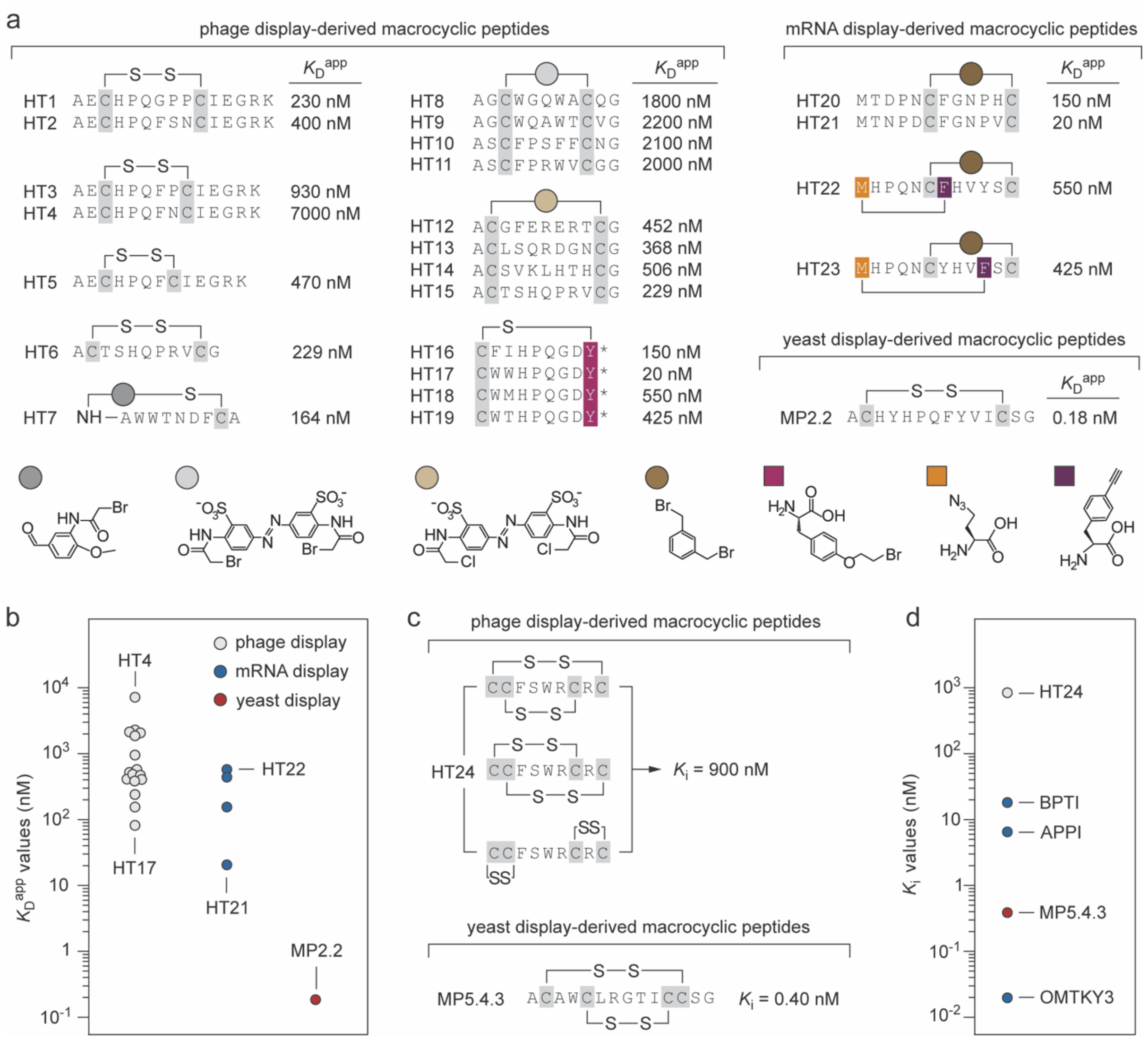


**Supplementary figure 6**. Phage and mRNA-derived macrocyclic peptide ligands selected against streptavidin and α-chymotrypsin. **a**) Amino acid sequence and apparent dissociation constant ($K_D^{app}$) values of some macrocyclic peptide ligands selected against streptavidin. Disulfide-tethered macrocyclic peptide HT1, HT2, HT3, HT4, HT5 were isolated using phage display technology (format naïve libraries: $CX_4C$, $CX_5C$ and $CX_6C$; libraries diversity: ~$10^8$). $K_D^{app}$ values were determined using surface plasmon resonance.[7] Disulfide-tethered macrocyclic peptide HT6 was identified using phage display technology (format naïve library: $CX_7C$; library diversity: ~$10^9$). $K_D^{app}$ value was determined by ESI-MS binding assay.[8] Macrocyclic peptide HT7 was discovered using phage display technology (format naïve library: $AX_8C$; library diversity: ~$10^8$). Post-translational macrocyclization of linear peptide sequence was performed by using an asymmetric bifunctional scaffold (dark grey circle) that selectively bridges the free N-terminal amine of alanine with the nearby unique cysteine residue. $K_D^{app}$ value was determined using a 4′-hydroxy-azobenzene-2-carboxylic acid (HABA) based competition assay. The binding affinity of an analogue of HT7 has also been measured using isothermal titration calorimetry ($K_D$ = 352 nM).[8] Macrocyclic peptides HT8, HT9, HT10 and HT11 were isolated using phage display technology (format naïve library: $ACX_7CG$; library diversity: ~$10^8$). Post-translational macrocyclization of linear peptide sequences has been performed using the bifunctional cyclization linker 3,3′-bis (sulfonato)-4,4′-bis(bromoacetamido) azobenzene (BSBBA, light grey circle) that selectively bridges the two free cysteine residues. $K_D^{app}$ values were determined using a fluorescence polarization-based assay.[9] Macrocyclic peptides HT12, HT13, HT14 and HT15 have been discovered using phage display technology (format naïve library: $CX_7C$; library diversity: ~$10^9$). Post-translational macrocyclization of linear peptide sequence has been performed using the bifunctional cyclization linker 3,3′-bis(sulfonato)-4,4′-bis(chloroacetamido)-azobenzene (BSBCA, light brown circle) that selectively bridges the two free cysteine residues. $K_D^{app}$ values were determined using an ESI-MS-based binding assay.[8] Macrocyclic peptides HT16, HT17, HT18 and HT19 constrained by a nonreducible thioether bridge were isolated using phage display technology (format naïve library: $CX_2HPQX_2Y$*; library diversity: ~$10^7$. Y* is the genetically incorporated cysteine-reactive noncanonical amino acid O-(2-bromoethyl)-tyrosine (O2beY). Macrocyclic peptides are constrained by a nonreducible, inter-side-chain-to-side-chain thioether linkage formed through a hemo- and regioselective reaction involving the genetically incorporated Y* (light purple) and a proximal cysteine residue. $K_D^{app}$ values were determined by ELISA assay.[10] Macrocyclic peptides HT20, HT21, HT22 and HT23 were identified using mRNA display technology starting from >$10^{13}$ member libraries. 'one ring' macrocyclic peptides HT20 and HT21 have been generated by cysteine alkylation using bifunctional α,α′-dibromo-*m*-xylene (dark brown circle) that selectively bridges the two free cysteine residues. The 'two rings' macrocyclic peptides HT22 and HT23 were generated by using the following orthogonal chemistries *i)* cysteine alkylation using bifunctional α,α′-dibromo-*m*-xylene and the *ii)* copper-mediated azide–alkyne cycloaddition (CuAAC) that selectively link the L-β-azidohomoalanine (AzHA, orange) to p-ethynyl phenylalanine (F-yne, dark purple) that were genetically incorporated in place of methionine (M) and phenylalanine (F), respectively. $K_D^{app}$ values were determined using a magnetic bead-based version of the spin filter binding-inhibition assay.[11] **b**) Plot of the $K_D^{app}$ values (nM) of unique streptavidin-binding macrocyclic peptide ligands (MP) isolated using phage display (grey filled circle), mRNA display (blue filled circle) and yeast display (this work, red filled circle); **c**) Amino acid sequence and inhibitory constant ($K_i$) values of some macrocyclic peptide inhibitors of α-chymotrypsin reported in literature. Macrocyclic peptides HT24 was discovered using phage display technology (format naïve library: $CX_7C$; library diversity: ~$10^9$). The inhibitory activity of the macrocyclic peptides was assessed by monitoring the residual activity of α-chymotrypsin in the presence of the chromogenic substrate N-Succinyl-Ala-Ala-Pro-Phe-*p*-nitroanilide and different concentrations of inhibitor macrocyclic peptides.[12] **d**) Plot of the $K_i$ values (nM) of unique macrocyclic peptide inhibitors (MP) of α-chymotrypsin isolated using phage display (grey filled circle) and yeast display (this work, red filled circle). As reference the $K_i$ values of the cysteine-rich Kunitz and Kazal small protein domains (~55 amino acids) such as the inhibitor domain of Alzheimer's amyloid P-protein precursor (APPI; $K_i$ = 7.1 nM), the basic pancreatic trypsin inhibitor (BPTI; $K_i$ = 16 nM) and the turkey ovomucoid third domain (OMTKY3; $K_i$ = 20 pM) are also plotted (blue dots).[13,14]

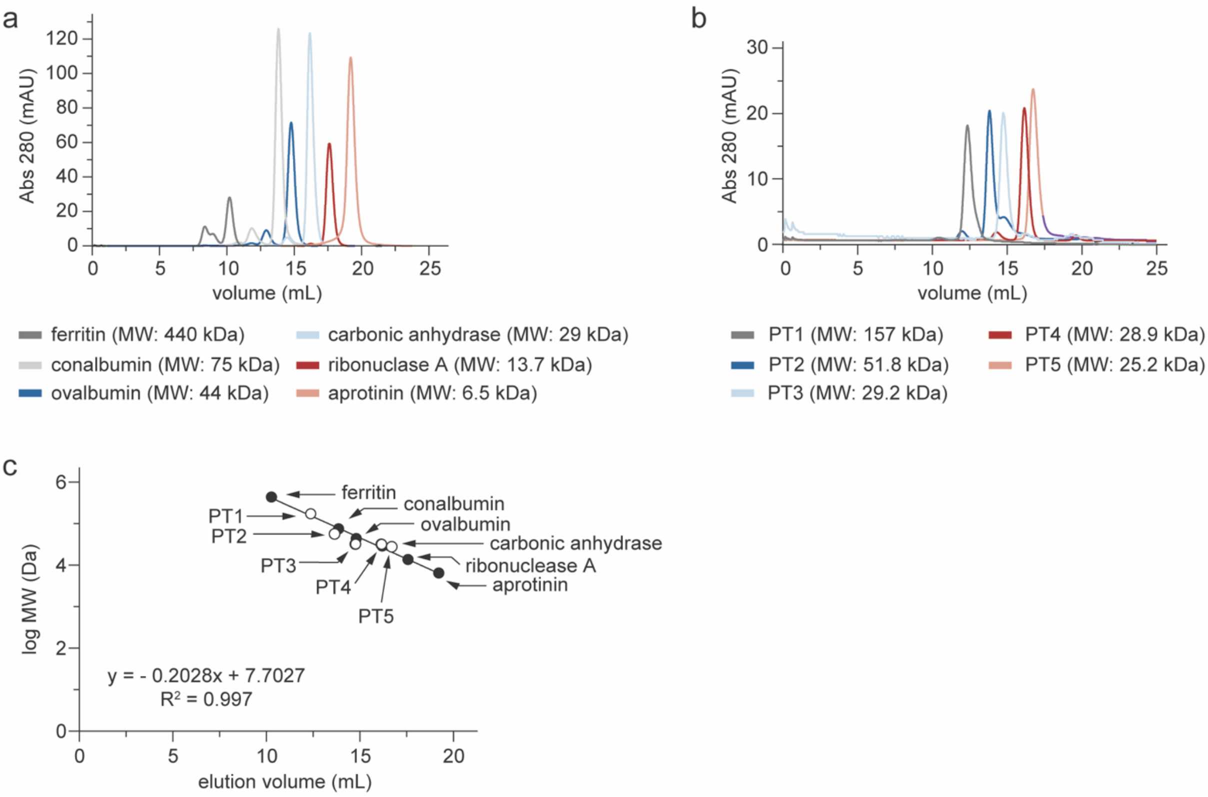


**Supplementary figure 7**. Size exclusion chromatography analyses of the five highly diverse protein targets used to validate the technology. **a**) Superdex 200 Increase 10/300 GL (GE, Healthcare) elution profiles of the calibration proteins ferritin (MW 440 kDa, dark grey), conalbumin (75 kDa, light grey), ovalbumin (44 kDa, dark blue), carbonic anhydrase (29 kDa, light blue), ribonuclease (13.7 kDa, dark red) and aprotinin (6.5 kDa, light red); **b**) Superdex 200 Increase 10/300 GL (GE, Healthcare) elution profiles of the protein target: PT1 (MW 157,4 kDa, dark grey), PT2 (51.8 kDa, dark blue), PT3 (29.2 kDa, light blue), PT4 (28.9 kDa, dark red) and PT5 (25.2 kDa, light red); **c**) Correlation between the logarithm of protein molecular weight (*y* axis) and the elution volumes (*x* axis). Calibration proteins are indicated by black circles, while grey circles represent the protein targets.

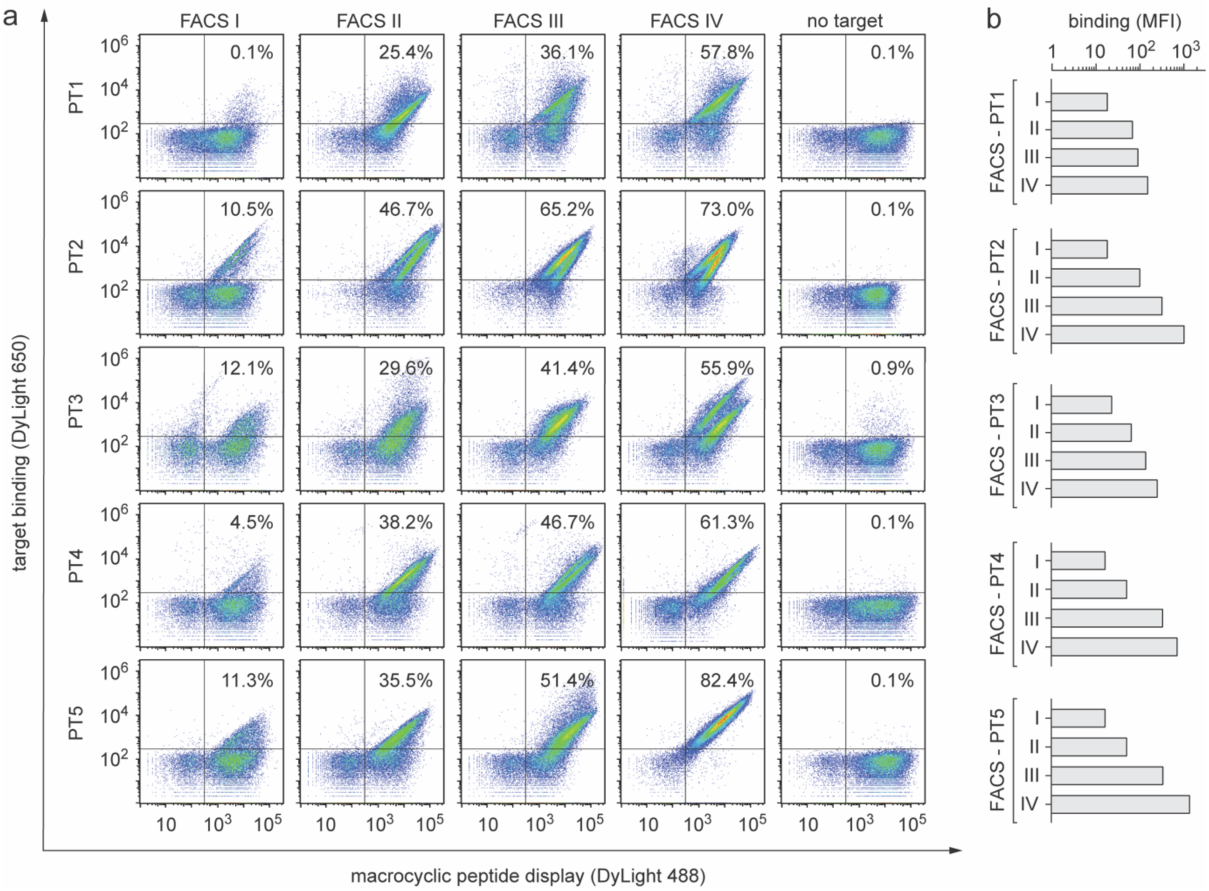


**Supplementary figure 8**. Sequential cycles of equilibrium-based fluorescence-activated cell sorting (FACS) selections. **a**) Density plots of polyclonal population of yeast cells encoding different macrocyclic peptides selected against multiple PTs that have been enriched through four cycles of FACS. Each dot represents two fluorescent signals of a single yeast cell. The fluorescence intensity on the *y*-axis is a measure of the amount of biotinylated PT bound to the surface of a yeast cell (DyLight 650; 'target binding') whereas the fluorescence intensity on the *x*-axis is a measure of the number of macrocyclic peptide molecules expressed on the surface of a yeast cell (DyLight 488; 'macrocyclic peptide display'); **b**) Columns graph reporting the geometric mean fluorescence (MFI) measured for the polyclonal population of yeast cells encoding different macrocyclic peptides against each PT through four cycles of FACS.

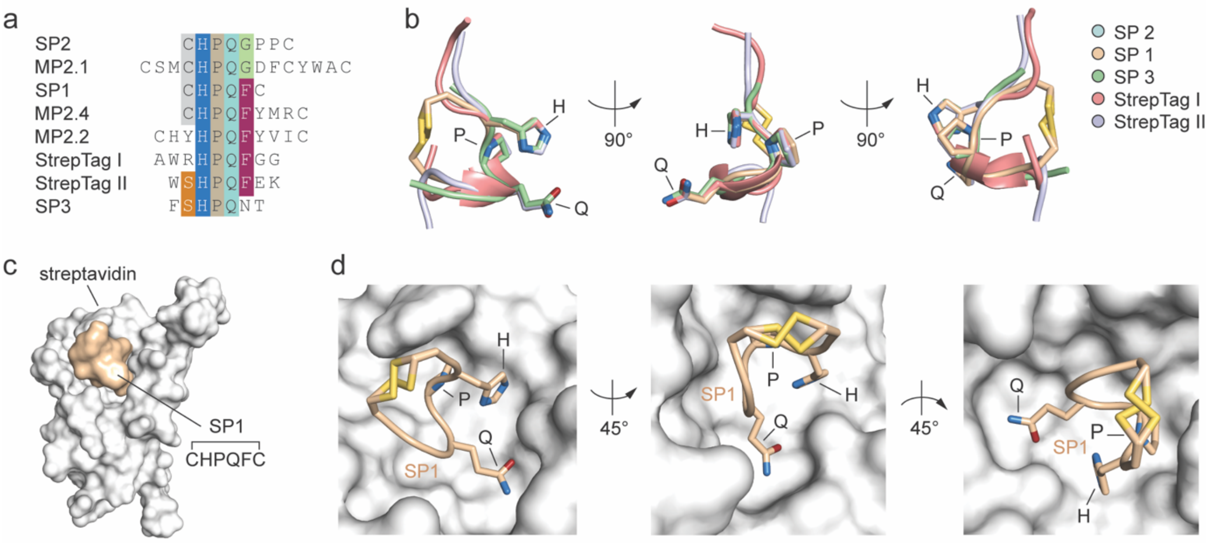


**Supplementary figure 9**. Structure of peptides including 'HPQ' motif in complex with streptavidin. **a**) Alignment of amino acids sequence of streptavidin-binding linear and cyclic peptide bearing the 'HPQ' motif. The amino acids are indicated as one letter code. Identical or similar amino acids between different peptide sequences are highlighted in colour (C: grey; H: indigo; P: light brown; Q: light blue; F: purple; G: light green; S: light orange); **b**) Detailed view of the three-dimensional structure superimposition of streptavidin-binding macrocyclic peptide SP1 (CHPQFC, wheat; PDB ID: 1SLD),[15] macrocyclic peptide SP2 (CHPQGPPC, pale cyan; PDB ID: 1SLE),[15] linear peptide SP3 (FSHPQNT, pale-green; PDB ID: 1VWA),[16] linear peptide StrepTag I (AWRHPQFGG, salmon; PDB ID: 1RST)[17] and linear peptide StrepTag II (WSHPQFEK, blue-white; PDB ID: 1RSU)[17] shown in three different orientations (90° rotation). The side chains of histidine (H), proline (P) and glutamine (Q) residues forming the 'HPQ' motif are shown as sticks; **c**) Molecular surface representation of streptavidin (grey) in complex with macrocyclic peptide SP1 (wheat; PDB ID: 1SLD);[15] **d**) Zoomed-in view of SP1 (wheat) bound to streptavidin (grey) shown in three different orientations (45° rotation). Amino acid side chains of the 'HPQ' motif are shown as sticks and coloured by atom type (carbon: wheat, oxygen: firebrick, nitrogen: deep blue, sulphur: yellow orange).

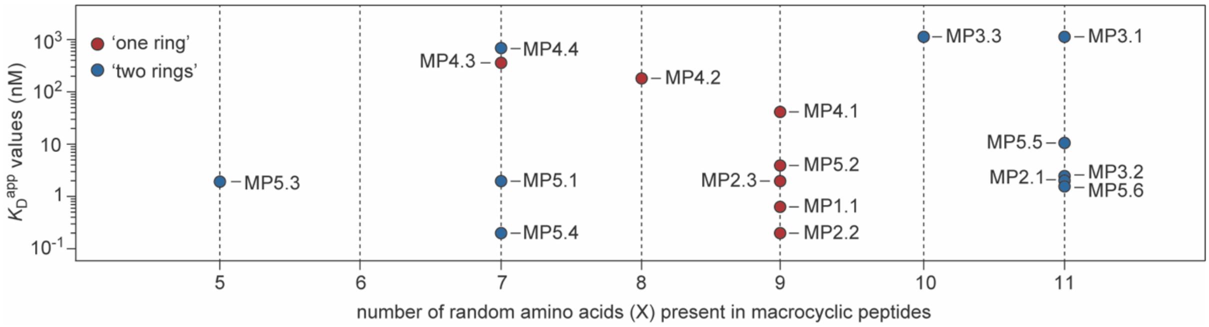


**Supplementary figure 10**. Plot of the binding affinities of seventeen unique yeast-encoded macrocyclic peptide ligands (MP) selected against five diverse protein targets versus the number of randomised amino acids present within the peptide sequences. Each colored filled circle indicates the apparent equilibrium binding constant value ($K_D^{app}$; nM) of a given MP. 'one ring' and 'two rings' macrocycle peptides are reported as red and blue filled colored circles, respectively.

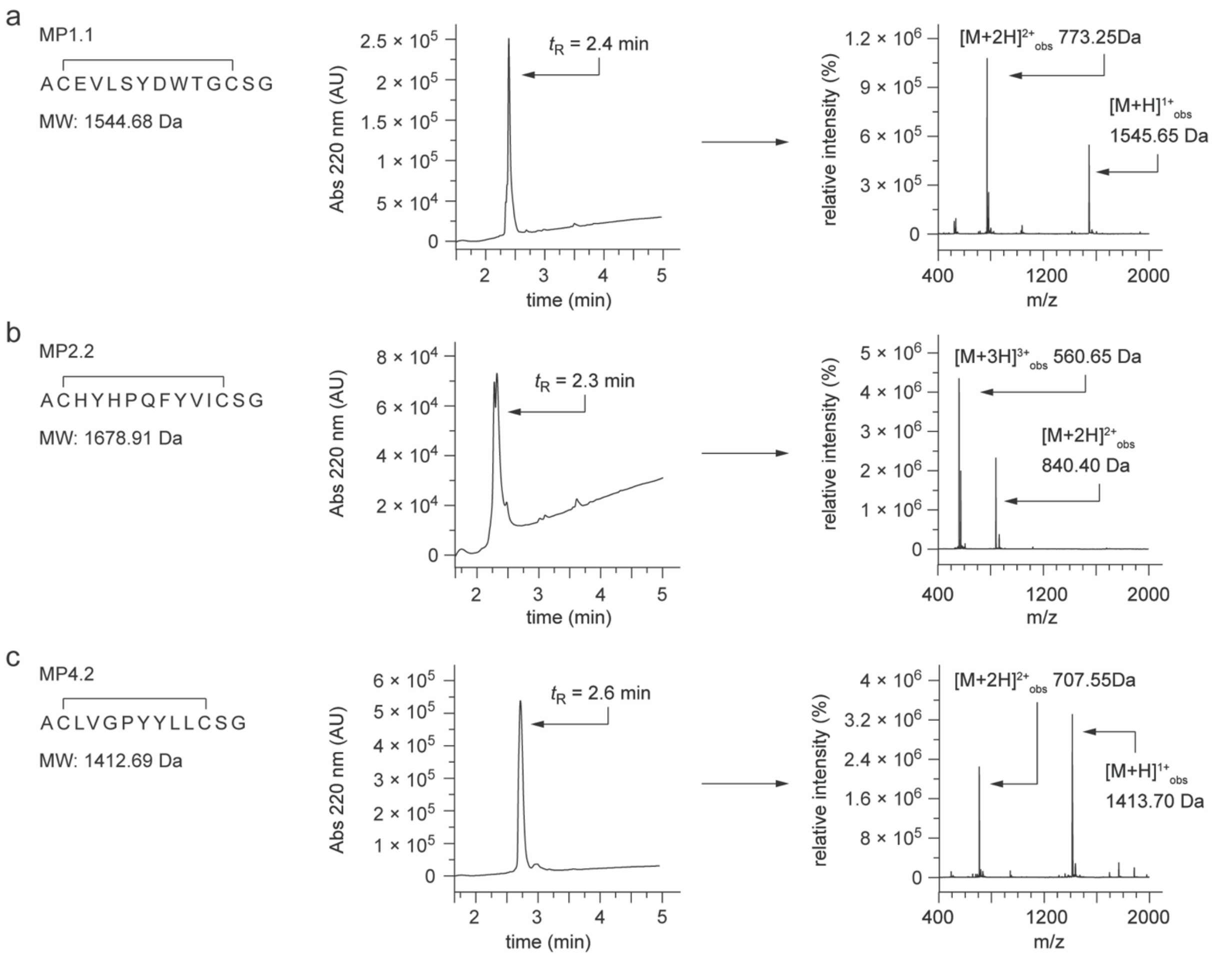


**Supplementary figure 11**. Synthesis and characterisation of 'one ring' macrocyclic peptides. HPLC (left) and mass spectra (right) analysis of macrocycles MP1.1 (**a**), MP2.2 (**b**) and MP4.2 (**c**). The measured molecular weight of each macrocyclic peptide corresponds to expected mass. Name, elution retention time ($t_R$), expected molecular weight (MW) and observed molecular ions of each macrocyclic peptide are indicated.

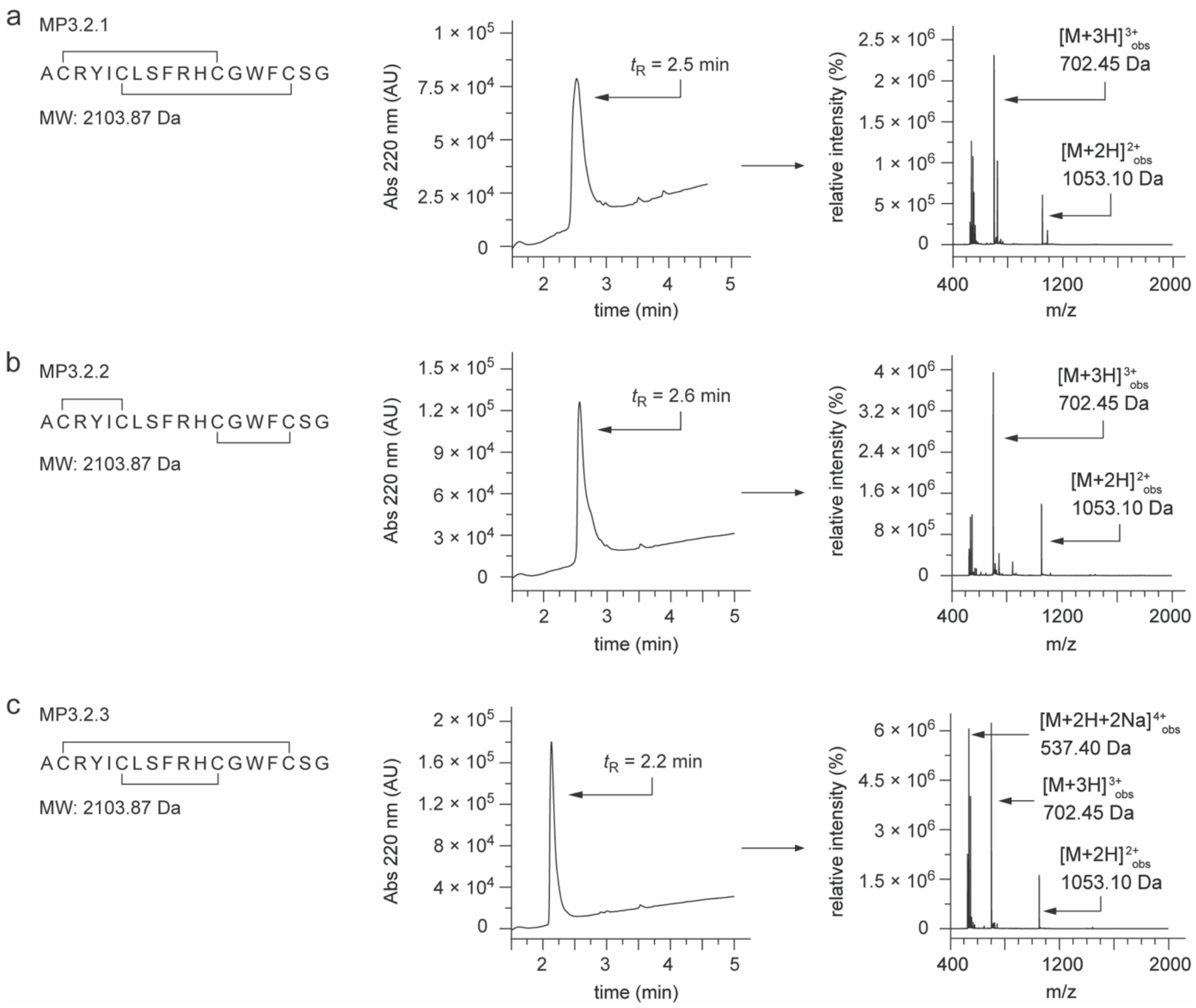


**Supplementary figure 12**. Synthesis and characterisation of ‘two rings’ macrocyclic peptides MP3.2.1, MP3.2.2 and MP3.2.3. HPLC (left) and mass spectra (right) analysis of MP3.2.1 (**a**), MP3.2.2 (**b**) and MP3.2.3 (**c**). The measured molecular weight of each ‘two rings’ macrocyclic peptide corresponds to expected mass. Name, elution retention time ($t_R$), expected molecular weight (MW) and observed molecular ions of each macrocyclic peptide are indicated.

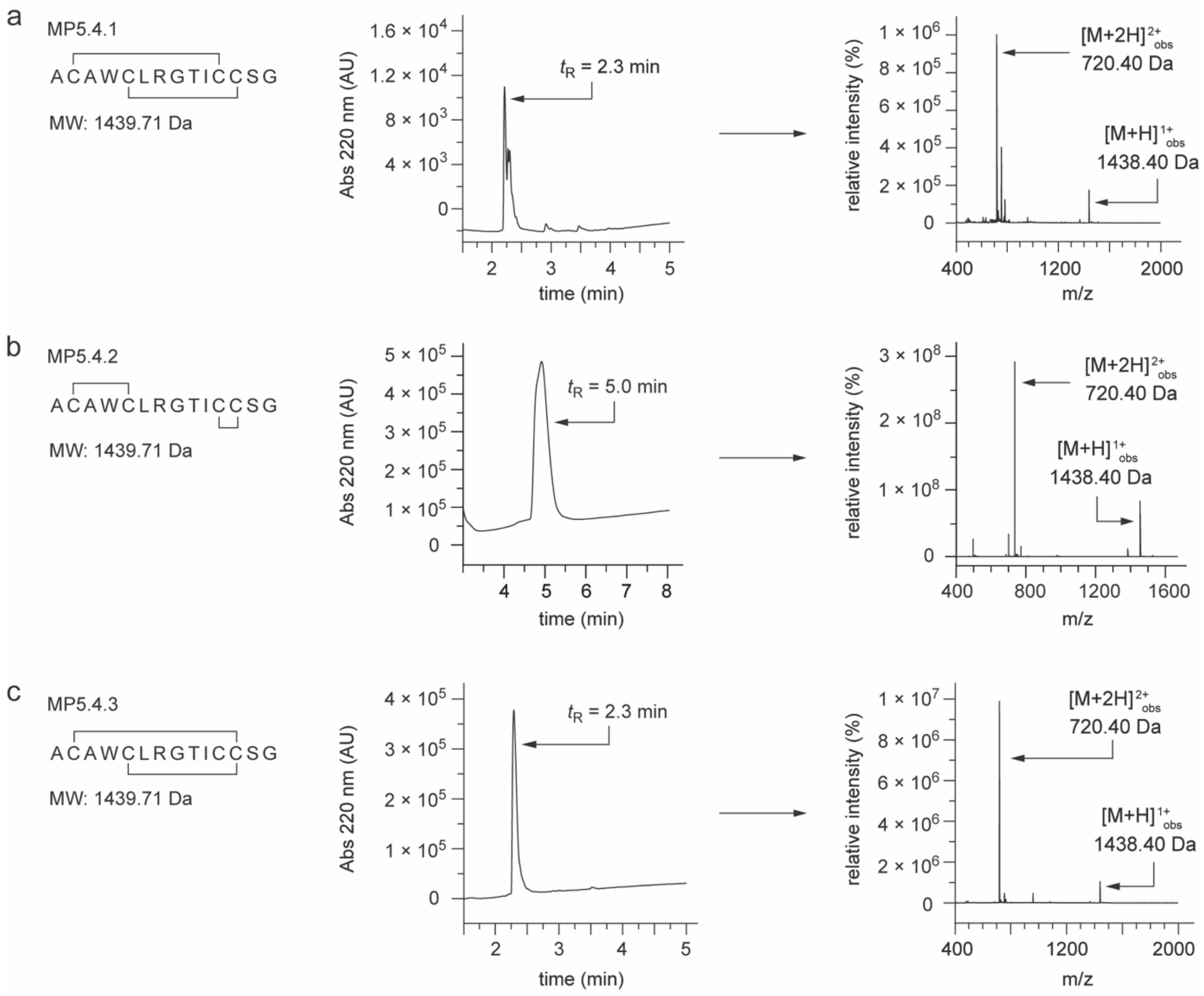


**Supplementary figure 13**. Synthesis and characterisation of 'two rings' macrocyclic peptides MP5.4.1, MP5.4.2 and MP5.4.3. HPLC (left) and mass spectra (right) analysis of MP5.4.1 (**a**), MP5.4.2 (**b**) and MP5.4.3 (**c**). The measured molecular weight of each 'two rings' macrocyclic peptide corresponds to expected mass. Name, elution retention time ($t_R$), expected molecular weight (MW) and observed molecular ions of each macrocyclic peptide are indicated.

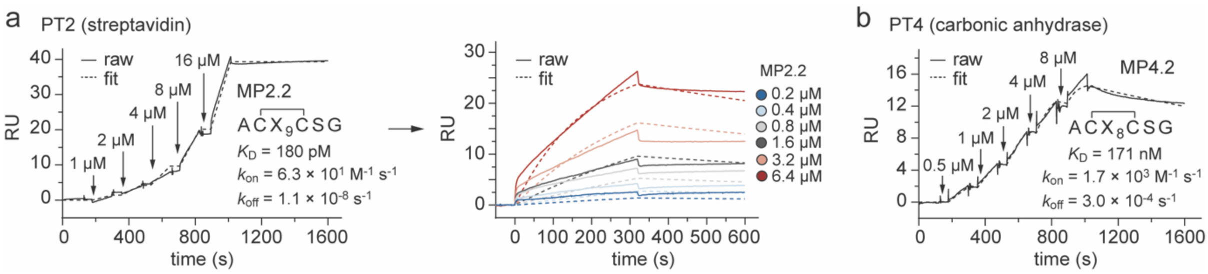


**Supplementary figure 14**. Surface plasmon resonance (SPR) sensorgram traces for the interaction of the immobilised biotinylated PT2 with soluble 'one ring' macrocycle peptide MP2.2 (**a**) and immobilised biotinylated PT4 with soluble 'one ring' macrocycle peptide MP4.2 (**b**). The injection and flow fill ranges have been removed from the presented sensorgram traces. Data were fitted using a 1:1 binding model. Kinetic constants $k_a$ (association constant), $k_d$ (dissociation constant) and $K_D$ (equilibrium binding affinity) are calculated applying LMW single-cycle kinetics 1:1 binding fitting or LMW multi cycle fitting. Raw data are shown as solid lines, while fit data are shown as dashed lines.

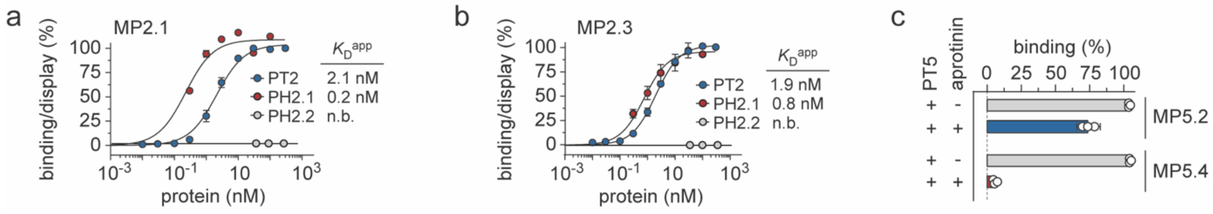


**Supplementary figure 15**. Binding isotherms of yeast-encoded MP2.1 (**a**) and MP2.3 (**b**) macrocycle peptides towards streptavidin (PT2, blue), strep-tactin (PH2.1, red) and neutravidin (PH2.2, grey). Apparent equilibrium binding affinities ($K_D^{app}$) values were determined only for PT2 and PH2.1. Data are presented as mean (dots) ± s.e.m. (bars). n.b., not binding; **c**) Competitive binding assay of yeast-encoded macrocyclic peptides MP5.2 and MP5.4 for binding to PT5 (bovine α-chymotrypsin) in the presence (+) or absence (-) of aprotinin.

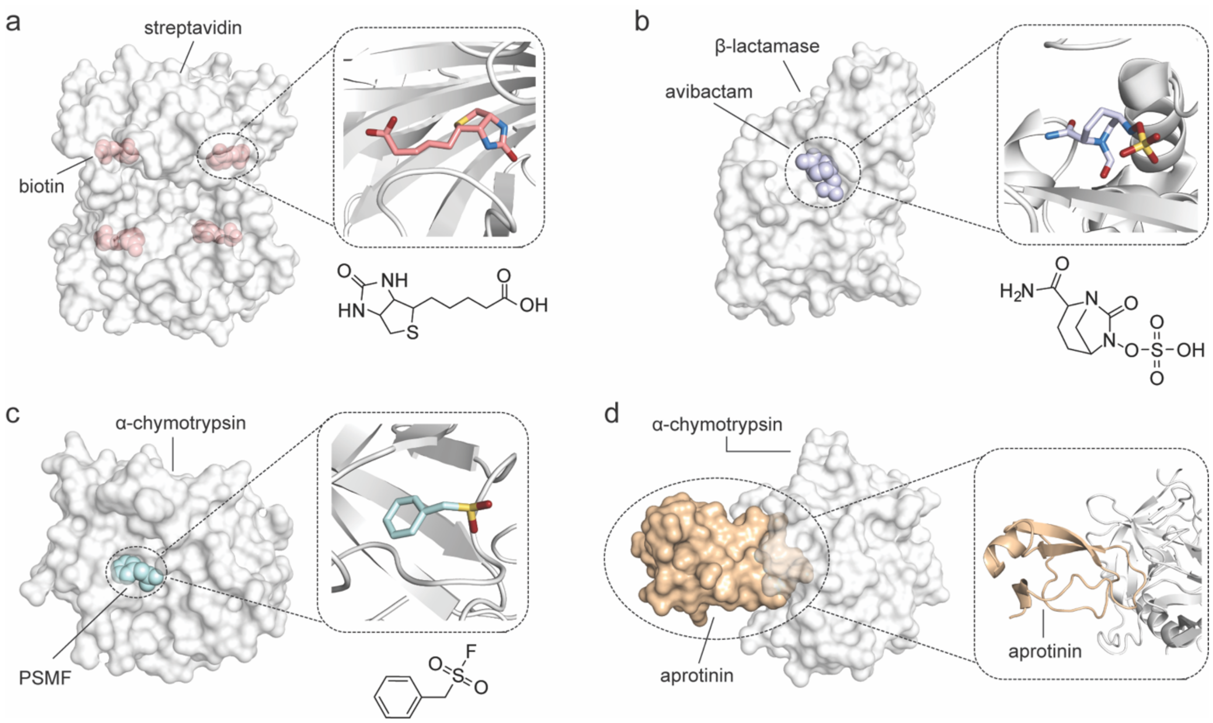


**Supplementary figure 16**. Crystal structure of some protein targets in complex with known site-specific and soluble binding ligand. **a**) Left, molecular surface representation of streptavidin (PT2; grey) in complex with biotin (carbon: salmon, nitrogen: sky-blue; oxygen: firebrick, sulphur: yellow-orange; PDB ID: 6J6J).[18] Right, zoomed-in view of the biotin molecule bound to streptavidin. The chemical structure of the biotin is shown; **b**) Left, molecular surface representation of β-lactamase CTX-M-15 (grey) in complex with avibactam (carbon: blue-white, nitrogen: sky-blue; oxygen: firebrick, sulphur: yellow-orange; PDB ID: 4S2I).[19] Right, zoomed-in view of avibactam bound to CTX-M-15. The chemical structure of avibactam is shown; **c**) Left, molecular surface representation of fire ant α-chymotrypsin (grey) in complex with phenylmethylsulfonyl fluoride (PSMF; carbon: pale cyan, nitrogen: sky-blue; oxygen: firebrick, sulphur: yellow-orange; PDB ID: 1EQ9).[20] Right, zoomed-in view of the PMSF structure bound to fire ant α-chymotrypsin. The chemical structure of PMSF is reported; **d**) Left, molecular surface representation of bovine α-chymotrypsin (grey) in complex with aprotinin (wheat; PDB ID: 1CA0).[13] Right, zoomed-in view of aprotinin structure bound to fire ant α-chymotrypsin.

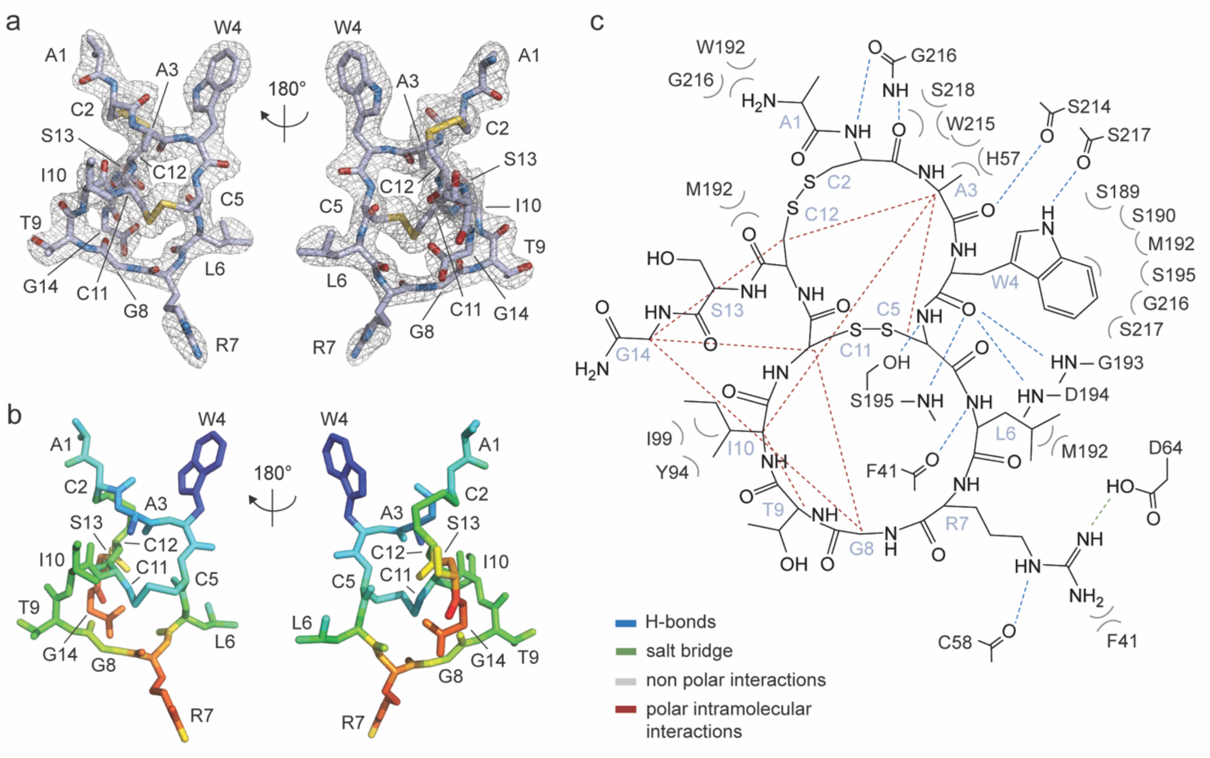


**Supplementary figure 17**. Electron density map and B-factor diagram of 'two rings' macrocyclic peptide MP5.4.3. **a**) Conformation and electron density map of MP5.4.3 shown in two orientations (180° rotation). The side chains of the amino acid residues are shown as sticks. Carbon, oxygen, nitrogen, and sulphur atoms are shown in light blue, red, sky-blue and yellow, respectively. The $F_O - F_C$ omit map is shown and contoured at the 2.5 σ level; **b**) B-factor diagram of MP5.4.3 shown in two orientations (180° rotation). The side chains of the residues are shown as sticks. The B-factor values are illustrated by colour, ranging from low (blue) to high (red); **c**) Schematic representation of molecular interactions between PT5 (bovine α-chymotrypsin) and MP5.4.3. Inter-molecular salt bridges and hydrogen bonds are shown as green and blue dashed lines, respectively. Macrocycle peptide intra-molecular polar interactions are shown as red dashed lines. Bent grey lines indicate residues of MP5.4.3 in close contact with PT5 (distances shorter than 4.0 Å that are not polar inter-molecular interactions). The three-dimensional structures were generated and rendered using PHENIX [21] and PyMOL.[22]

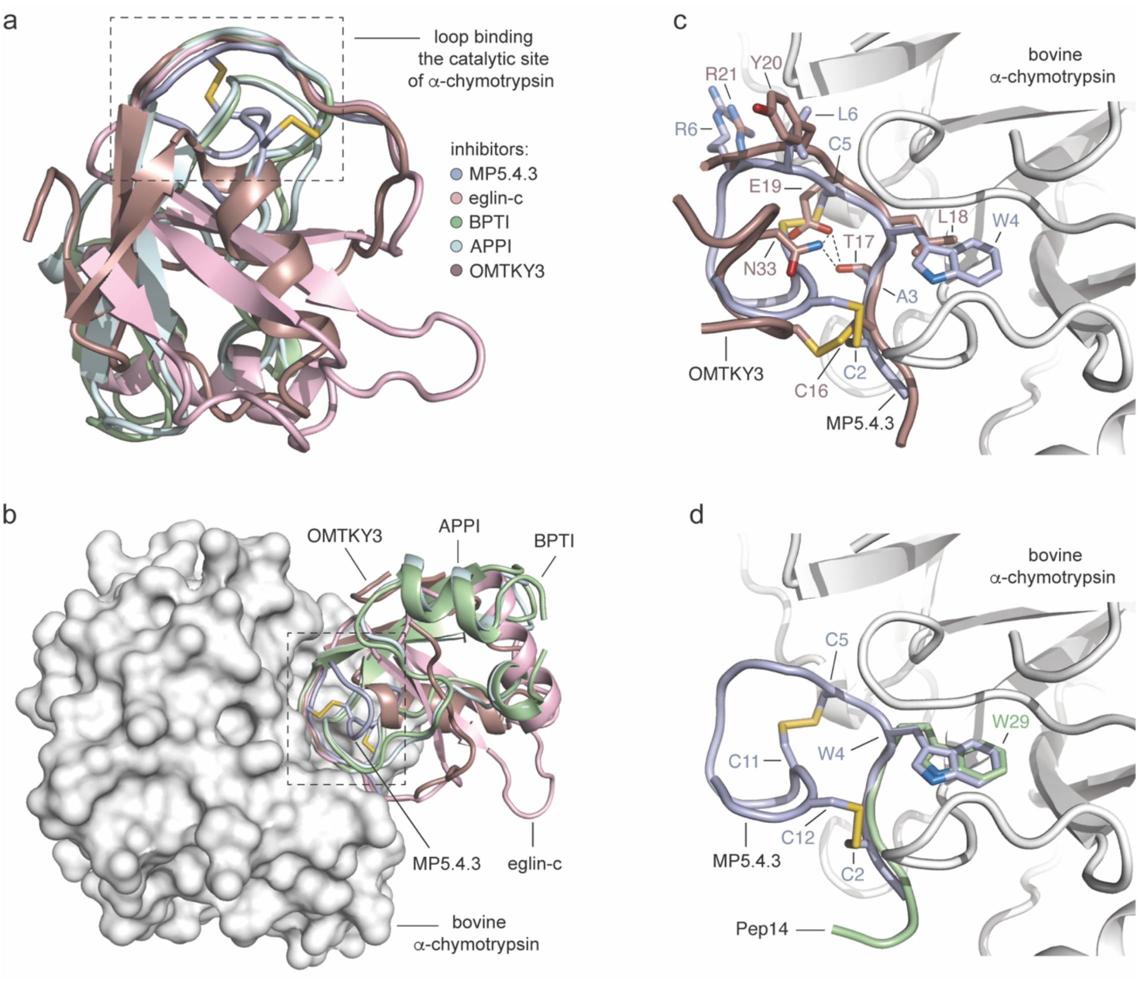


**Supplementary figure 18**. Superimposition of bovine α-chymotrypsin in complex with 'two rings' macrocyclic peptide MP5.4.3, natural and designed small protein-based inhibitors. **a**) Superimposition of aligned bovine α-chymotrypsin in complex with yeast-encoded macrocyclic peptide MP5.4.3 (PDB ID: 9F6H; light blue), eglin-c, a small thermostable protein isolated from the leech Hirudo medicinalis (PDB ID: 1ACB; light pink),[23] the bovine pancreatic trypsin inhibitor (BPTI; PDB ID: 1CBW; pale green), a small member of the protein family of Kunitz-type serine protease inhibitors,[13] the Kunitz protease inhibitor domain of the amyloid precursor protein (APPI; PDB ID: 1CA0; pale cyan)[13] and the third Kazal domain of the turkey ovomucoid protein (OMTKY3; PDB ID: 1CHO; dark salmon)[24]; **b**) Superimposition of aligned bovine α-chymotrypsin in complex with a six amino acid portion ($^{24}$PGSWPW$^{29}$) of Pep14, a native fourteen amino acid fragment, from Ile16 to Trp29 ($^{16}$IVNGEEAVPGSWPW$^{29}$), that is generated by the autolysis of α-chymotrypsin and that is known to bind to the active site of the same enzyme (PDB ID: 1OXG; pale green)[25]. The three-dimensional structures were generated and rendered using PyMOL.[22]

**Amino acid count MATLAB code**

```
function AminoCount(varargin)
%% INPUT SECTION
inname = '';
outdir = ''; % default save directory is a new directory inside the input directory
outname= 'amino';   % default save name
indir = '';
Cter = '';
CUTOFF = 0;

% check for input variable
if exist('varargin','var')
    L = length(varargin);
    if rem(L,2) ~= 0, error('Parameters/Values must come in pairs.');
    end

    % read input variables
    for ni = 1:2:L
        switch lower(varargin{ni})
            case 'inname', inname = varargin{ni+1};
            case 'outdir', outdir = varargin{ni+1};
            case 'indir', indir=varargin{ni+1};
            case 'cutoff', CUTOFF=varargin{ni+1};
            case 'cter', Cter=varargin{ni+1};
        end
    end
end

% check if inname was defined
if strcmp(inname,'')
    [inname,indir,~] = uigetfile('*.txt','Select .txt file');
```

```
else
    [~,message] = fopen(fullfile(indir, inname));
    if strcmp(message,'') == 0
        display('File not found, a dialog box will open...');
        [inname,indir,~] = uigetfile('*.txt','Select .txt file');
    end;
end;



if strcmp(outdir,'')
    mkdir(indir, ['amino_'  strrep(inname,'.txt','')] );
    outdir = ['amino_'  strrep(inname,'.txt','')];
end;
%mkdir(fullfile(indir, outdir, 'other'));



% check if Cter was specified
if strcmp(Cter,'')
    display('No C-terminus specified');
else
    display(['Considering sequences whose C-terminus is ' Cter]);
end;

% check if cutoff was specified
if CUTOFF == 0
    display('No abundance cutoff applied');
else
    display(['Considering sequences with a minimum abundance of ' num2str(CUTOFF)]);
end;

%% DATA READING
%open file and read data
```

```
file = fopen(fullfile(indir, inname));
AllVar = textscan(file, '%s %d %s %*[^\n]');
fclose('all');

AllSeq = AllVar{1}; %Sequences are stored as a cell array of strings
AllOccur = AllVar{2};
AllNtd = AllVar{3};

KEEP = find(AllOccur>=CUTOFF); % Discard sequences that appeared less than 'CUTOFF' times)
AllSeq = AllSeq(KEEP);
AllOccur = AllOccur(KEEP);
AllNtd = AllNtd(KEEP);

if strcmp(Cter,'')
else
   KEEP = ~cellfun('isempty',(strfind(AllSeq,Cter)));
   AllSeq = AllSeq(KEEP);
   AllOccur = AllOccur(KEEP);
   AllNtd = AllNtd(KEEP);
end;

if numel(AllSeq)>1 && ischar(AllSeq{1}) && isnumeric(AllOccur(1)) && ischar(AllNtd{1})
   total_seq_considered = sum(AllOccur);
   display(['Total sequences considered = ' num2str(total_seq_considered)]);
else
   display('Error: file chosen not suitable for this script');
   display('Choose a file of the format: peptide seq - abundance - nucleotide seq');
   return
end;
```

```
clear('AllVar');

%% DATA ANALYSIS: counting aminoacids

total_seq = sum(AllOccur);
cys = strfind(AllSeq,'C');
ala = strfind(AllSeq,'A');
leu = strfind(AllSeq,'L');
val = strfind(AllSeq,'V');
ile = strfind(AllSeq,'I');
asn = strfind(AllSeq,'N');
asp = strfind(AllSeq,'D');
arg = strfind(AllSeq,'R');
glu = strfind(AllSeq,'E');
gln = strfind(AllSeq,'Q');
azm = strfind(AllSeq,'Z');
gly = strfind(AllSeq,'G');
his = strfind(AllSeq,'H');
lys = strfind(AllSeq,'K');
met = strfind(AllSeq,'M');
phe = strfind(AllSeq,'F');
pro = strfind(AllSeq,'P');
ser = strfind(AllSeq,'S');
thr = strfind(AllSeq,'T');
trp = strfind(AllSeq,'W');
tyr = strfind(AllSeq,'Y');
star = strfind(AllSeq,'*');
total_cys = 0;
total_ala = 0;
total_leu = 0;
total_val = 0;
```

```
total_ile = 0;
total_asn = 0;
total_asp = 0;
total_arg = 0;
total_glu = 0;
total_gln = 0;
total_azm = 0;
total_gly = 0;
total_his = 0;
total_lys = 0;
total_met = 0;
total_phe = 0;
total_pro = 0;
total_ser = 0;
total_thr = 0;
total_trp = 0;
total_tyr = 0;
total_star = 0;

for i=1: size(AllSeq)
  total_cys = total_cys + numel(cys{i})*AllOccur(i);
  total_ala = total_ala + numel(ala{i})*AllOccur(i);;
  total_leu = total_leu + numel(leu{i})*AllOccur(i);;
  total_val = total_val + numel(val{i})*AllOccur(i);;
  total_ile = total_ile + numel(ile{i})*AllOccur(i);;
  total_asn = total_asn + numel(asn{i})*AllOccur(i);;
  total_asp = total_asp + numel(asp{i})*AllOccur(i);;
  total_arg = total_arg + numel(arg{i})*AllOccur(i);;
  total_glu = total_glu + numel(glu{i})*AllOccur(i);;
  total_gln = total_gln + numel(gln{i})*AllOccur(i);;
  total_azm = total_azm + numel(azm{i})*AllOccur(i);;
  total_gly = total_gly + numel(gly{i})*AllOccur(i);;
```

```
    total_his = total_his + numel(his{i})*AllOccur(i);;

    total_lys = total_lys + numel(lys{i})*AllOccur(i);;

    total_met = total_met + numel(met{i})*AllOccur(i);;

    total_phe = total_phe + numel(phe{i})*AllOccur(i);;

    total_pro = total_pro + numel(pro{i})*AllOccur(i);;

    total_ser = total_ser + numel(ser{i})*AllOccur(i);;

    total_thr = total_thr + numel(thr{i})*AllOccur(i);;

    total_trp = total_trp + numel(trp{i})*AllOccur(i);;

    total_tyr = total_tyr + numel(tyr{i})*AllOccur(i);;

    total_star = total_star + numel(star{i})*AllOccur(i);

end;

%total_amino = total_cys + total_ala + total_leu + total_val + total_ile + total_asn + total_asp +
total_arg + total_glu + total_gln + total_azm + total_gly +  total_his + total_lys + total_met +
total_phe + total_pro +  total_ser + total_thr + total_trp + total_tyr + total_star;



%% WRITE FILES from adaptor sequences analysis


%Printing stats files


fh = fopen(fullfile(indir,outdir,'', [outname '_counting.csv']),'w');

fprintf(fh, '%s%d%s\n', 'tot. sequences: ', total_seq, ';');

fprintf(fh, '%s\n','Aminoacid;count;count/tot_sequences;');

fprintf(fh, '%s%d%s%.4f%s\n', 'cys;', total_cys, ';', double(total_cys)/double(total_seq), ';');

fprintf(fh, '%s%d%s%.4f%s\n', 'ala;', total_ala, ';', double(total_ala)/double(total_seq), ';');

fprintf(fh, '%s%d%s%.4f%s\n', 'leu;', total_leu, ';', double(total_leu)/double(total_seq), ';');

fprintf(fh, '%s%d%s%.4f%s\n', 'val;', total_val, ';', double(total_val)/double(total_seq), ';');

fprintf(fh, '%s%d%s%.4f%s\n', 'ile;', total_ile, ';', double(total_ile)/double(total_seq), ';');

fprintf(fh, '%s%d%s%.4f%s\n', 'asn;', total_asn, ';', double(total_asn)/double(total_seq), ';');

fprintf(fh, '%s%d%s%.4f%s\n', 'asp;', total_asp, ';', double(total_asp)/double(total_seq), ';');

fprintf(fh, '%s%d%s%.4f%s\n', 'arg;', total_arg, ';', double(total_arg)/double(total_seq), ';');

fprintf(fh, '%s%d%s%.4f%s\n', 'glu;', total_glu, ';', double(total_glu)/double(total_seq), ';');

fprintf(fh, '%s%d%s%.4f%s\n', 'gln;', total_gln, ';', double(total_gln)/double(total_seq), ';');
```

```
fprintf(fh, '%s%d%s%.4f%s\n', 'azm;', total_azm, ';', double(total_azm)/double(total_seq), ';');
fprintf(fh, '%s%d%s%.4f%s\n', 'gly;', total_gly, ';', double(total_gly)/double(total_seq), ';');
fprintf(fh, '%s%d%s%.4f%s\n', 'his;', total_his, ';', double(total_his)/double(total_seq), ';');
fprintf(fh, '%s%d%s%.4f%s\n', 'lys;', total_lys, ';', double(total_lys)/double(total_seq), ';');
fprintf(fh, '%s%d%s%.4f%s\n', 'met;', total_met, ';', double(total_met)/double(total_seq), ';');
fprintf(fh, '%s%d%s%.4f%s\n', 'phe;', total_phe, ';', double(total_phe)/double(total_seq), ';');
fprintf(fh, '%s%d%s%.4f%s\n', 'pro;', total_pro, ';', double(total_pro)/double(total_seq), ';');
fprintf(fh, '%s%d%s%.4f%s\n', 'ser;', total_ser, ';', double(total_ser)/double(total_seq), ';');
fprintf(fh, '%s%d%s%.4f%s\n', 'thr;', total_thr, ';', double(total_thr)/double(total_seq), ';');
fprintf(fh, '%s%d%s%.4f%s\n', 'trp;', total_trp, ';', double(total_trp)/double(total_seq), ';');
fprintf(fh, '%s%d%s%.4f%s\n', 'tyr;', total_tyr, ';', double(total_tyr)/double(total_seq), ';');
fprintf(fh, '%s%d%s%.4f%s\n', '*;', total_star, ';', double(total_star)/double(total_seq), ';');

fclose('all');
```
